# Direct molecular evidence for an ancient, conserved developmental toolkit controlling post-transcriptional gene regulation in land plants

**DOI:** 10.1101/2021.03.04.433974

**Authors:** Haiyan Jia, Kelsey Aadland, Oralia Kolaczkowski, Bryan Kolaczkowski

## Abstract

RNA interference (RNAi) plays important roles in organism development through post-transcriptional regulation of specific target mRNAs. Target specificity is largely controlled by base-pair complementarity between micro-RNA (miRNA) regulatory elements and short regions of the target mRNA. The pattern of miRNA production in a cell interacts with the cell’s mRNA transcriptome to generate a specific network of post-transcriptional regulation that can play critical roles in cellular metabolism, differentiation, tissue/organ development and developmental timing. In plants, miRNA production is orchestrated in the nucleus by a suite of proteins that control transcription of the pri-miRNA gene, post-transcriptional processing and nuclear export of the mature miRNA. In the model plant, *Arabidopsis thaliana*, post-transcriptional processing of miRNAs is controlled by a pair of physically-interacting proteins, HYL1 and DCL1. However, the evolutionary history of the HYL1-DCL1 interaction is unknown, as is its structural basis. Here we use ancestral sequence reconstruction and functional characterization of ancestral HYL1 *in vitro* and *in vivo* to better understand the origin and evolution of the HYL1-DCL1 interaction and its impact on miRNA production and plant development. We found the ancestral plant HYL1 evolved high affinity for both double-stranded RNA (dsRNA) and its DCL1 partner very early in plant evolutionary history, before the divergence of mosses from seed plants (~500 Ma), and these high-affinity interactions remained largely conserved throughout plant evolutionary history. Structural modeling and molecular binding experiments suggest that the second of two double-stranded RNA-binding motifs (DSRMs) in HYL1 may interact tightly with the first of two C-terminal DCL1 DSRMs to mediate the HYL1-DCL1 physical interaction necessary for efficient miRNA production. Transgenic expression of the nearly 200 Ma-old ancestral flowering-plant HYL1 in *A. thaliana* was sufficient to rescue many key aspects of plant development disrupted by HYL1^−^ knockout and restored near-native miRNA production, suggesting that the functional partnership of HYL1-DCL1 originated very early in and was strongly conserved throughout the evolutionary history of terrestrial plants. Overall, our results are consistent with a model in which miRNA-based gene regulation evolved as part of a conserved plant ‘developmental toolkit’; its role in generating developmental novelty is probably related to the relatively rapid evolution of miRNA genes.

## INTRODUCTION

Understanding the factors that have contributed to similarities and differences among living species has intrigued humanity since archaic times, informing cultural myths and generating much of the impetus for biological science. Before the genomic era, it was widely believed that changes in gene complement were primarily responsible for phenotypic differences across species (Pertea and Salzberg 2010). However, studies of the evolution of development have highlighted the fact that changes in gene regulation can be just as important for generating novel biodiversity (Romero et al. 2012). Evolution of transcription factors controlling when, where and how much of a gene’s messenger RNA (mRNA) is produced - or of the DNA regulatory elements transcription factors recognize - can alter the patterns of gene regulation across species, leading to differences in developmental timing, body plan, tissue/organ development and cell differentiation (Quattrocchio et al. 1999; Graham et al. 2000; Davidson and Erwin 2006; He and Deem 2010; Mansfield 2013; Hudry et al. 2014; Jill Harrison 2017; Szövényi et al. 2019).

Transcription factors are not the only layer of regulation controlling gene function (Martinez and Walhout 2009). RNA interference (RNAi) is an additional gene-regulatory mechanism that affects mRNAs after they have been transcribed (Agrawal et al. 2003). Small RNAs interfere with gene-specific protein synthesis by guiding Argonaute (AGO) proteins to specific mRNA targets via sequence complementarity (Hutvagner and Simard 2008; Sheu-Gruttadauria and MacRae 2017). RNAi is found in all major eukaryote lineages (Cerutti and Casas-Mollano 2006), with various lineage-specific secondary losses (Obbard et al. 2009; Zhang et al. 2011; Billmyre et al. 2013; Mukherjee et al. 2013; Jeseničnik et al. 2019). Although recent studies suggest that endogenous mRNA regulation by RNAi may be facilitated by unique mechanisms in protozoa and fungi (Dang et al. 2011; Billmyre et al. 2013; Zhang et al. 2018), the most well-studied mechanism of endogenous RNAi-facilitated gene regulation occurs via micro-RNAs encoded in animal and plant genomes (Moran et al. 2017).

Mature micro-RNAs (miRNAs) are short (~22nt) single-stranded RNAs produced from genomic pri-miRNA genes (Ha and Kim 2014), many of which can mediate various aspects of developmental programming (Ge et al. 2012; Alberti and Cochella 2017). Animals and plants both use miRNAs for mRNA regulation but generate mature miRNAs using different processes (Axtell et al. 2011). Animals generally export pre-miRNA ‘hairpins’ from the nucleus to the cytoplasm, where they are further processed into mature miRNAs by a complex formed by a physical interaction between a double-stranding RNA-binding protein (DRB) and the endoribonuclease, Dicer (Kim et al. 2016; O’Brien et al. 2018). In contrast, mature plant miRNAs are typically generated completely in the nucleus by homologs of animal DRB-Dicer, before export to the cytoplasm (Bollman et al. 2003; Kurihara and Watanabe 2004; Kurihara et al. 2006; Axtell et al. 2011; Bologna et al. 2018). Although animals and plants both utilize a DRB-Dicer complex for the final step in mature miRNA production, the structural interface facilitating complex formation is different (Dias et al. 2017), suggesting the DRB-Dicer complex may have evolved convergently in animals and plants. Coupled with the lack of clearly-shared pri-miRNA genes between animal and plant lineages, marked differences in miRNA biogenesis between the two groups has led researchers to suggest that miRNA-based gene regulation probably evolved independently in animals and plants via convergent elaboration of a more simplified ancestral RNAi mechanism (Shabalina and Koonin 2008; Axtell et al. 2011; Tarver et al. 2012).

While many of the details of miRNA biogenesis have been documented in the model plant, *Arabidopsis thaliana*, much less is known about when the various components required for endogenous miRNA-based gene regulation arose or how the plant miRNA biogenesis system evolved. Recent examinations of miRNA-gene evolution suggest that the earliest plant genomes encoded ~14 conserved miRNA genes, although the large number of identified species-specific miRNAs suggests that early plants could have had a richer miRNA-gene complement, many of which were lost or altered beyond recognition in modern species (Willmann and Poethig 2007). MicroRNA biogenesis in *A. thaliana* is orchestrated by the Dicer homolog, DCL1 and the DRB homolog, HYL1 (Hammond et al. 2000; Martinez et al. 2002; Kurihara and Watanabe 2004; Hiraguri et al. 2005; Kurihara et al. 2006; Franco-Zorrilla et al. 2007; Yang et al. 2010; Yang et al. 2014). Studies of the evolution of these protein families suggest that HYL1 originated in very early plants (Axtell et al. 2011; Dias et al. 2017), and although DCL1 homologs are found throughout eukaryotes, many of the key features of plant DCL1 responsible for efficient miRNA processing are thought to have evolved around the origin of multicellular plants (Jia et al. 2017). Whether the HYL1-DCL1 interaction responsible for *A. thaliana* miRNA biogenesis arose early in plants or was a later elaboration of a simpler RNAi system remains unknown, and the evolutionary dynamics of the HYL1-DCL1 complex have not been investigated.

Here we use ancestral protein resurrection *in vitro* and in live *A. thaliana* to directly investigate the evolution of HYL1’s role in plant miRNA biogenesis. We found that HYL1’s capacity to mediate miRNA biogenesis arose early in plants and was largely maintained over the hundreds of millions of years of evolutionary history between its origin and modern plant species. Interestingly, the reconstructed ancestral HYL1 of flowering plants - which existed nearly 200 Ma (Silvestro et al. 2021) - was sufficient to recover many of the aspects of the HYL1^−^ knockout phenotype in modern *A. thaliana* by orchestrating near-native miRNA biogenesis, despite low overall sequence similarity, strongly suggesting that the HYL1-DCL1 miRNA-processing complex evolved early in plants and remained strongly conserved throughout plant evolutionary history.

## RESULTS AND DISCUSSION

### HYL1 evolved high affinity for dsRNA and DCL1 early in plant evolutionary history

To begin examining how plant HYL1 evolved the capacity to mediate miRNA biogenesis by forming a complex with DCL1 and its pri-miRNA targets, we identified plant and animal double-stranded RNA-binding proteins (DRBs) and inferred a maximum-likelihood consensus tree for this protein family, integrating results from different alignment strategies (see Methods). Consistent with our previous analysis (Dias et al. 2017), we found that plant DRBs were monophyletic with high statistical confidence (>0.97 SH-like aLRT, depending on the alignment), and plant HYL1+DRB6 formed a monophyletic lineage with support >0.92 (Figure 1; Supplementary Information Figure S1). The HYL1-DRB6 duplication event appears to have occurred before the divergence of mosses from seed plants. Although the support for HYL1 monophyly was sometimes low (e.g., SH-like aLRT=0.77 using the mafft alignment), the HYL1 lineage was recovered in the maximum-likelihood tree using all sequence alignments, with support reaching 0.99 in one case and averaging 0.85 across all alignments (see Supplementary Information Figure S1). The major plant lineages were all recovered in the HYL1 consensus tree, with the overall branching pattern generally congruent with recent plant species phylogenies (Ruhfel et al. 2014; Wickett et al. 2014).

**Figure 1.**
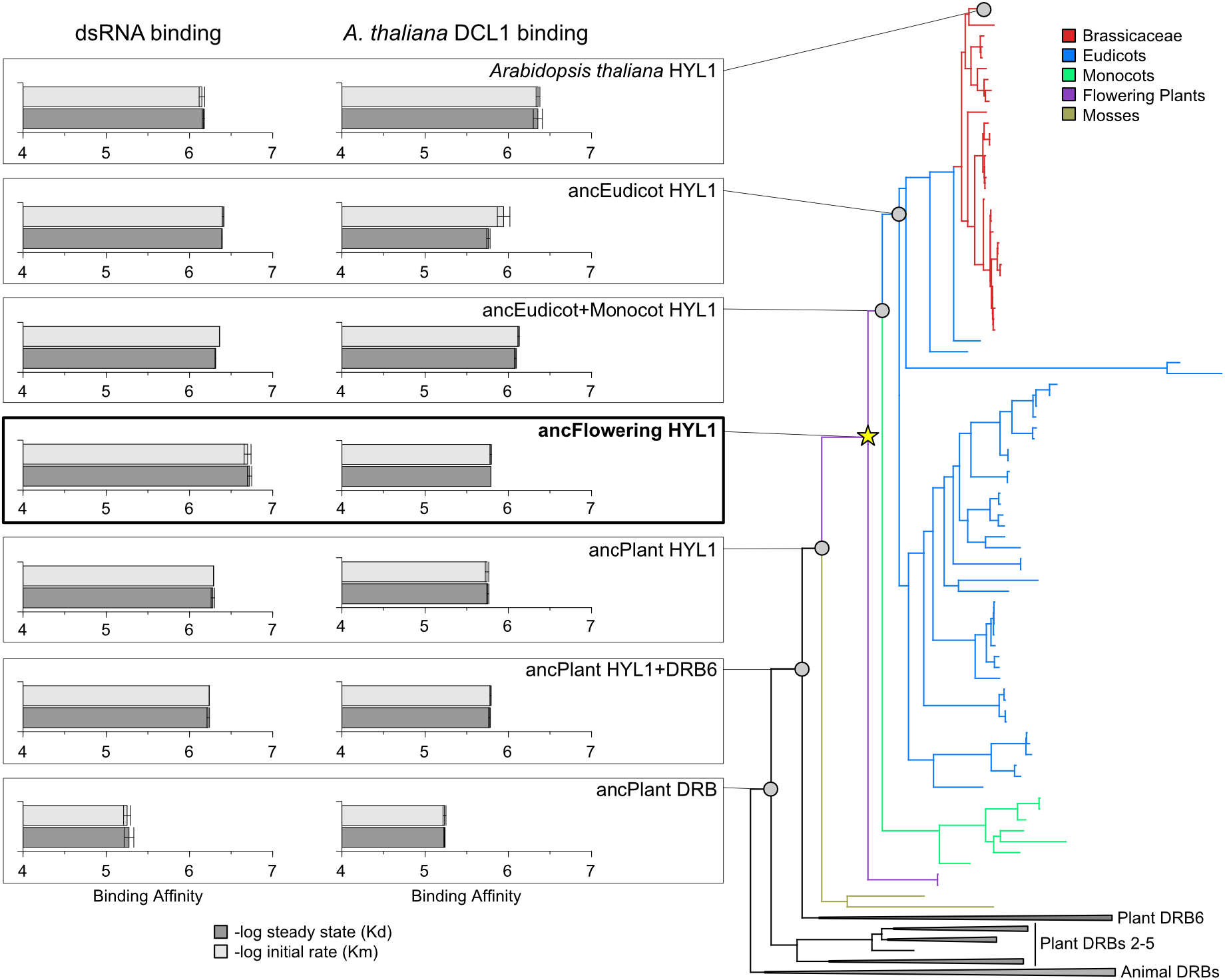
Ancestral double-stranded RNA binding proteins (DRBs) evolved high affinity for dsRNA and *Arabidopsis thaliana* DCL1 early in plant evolutionary history. Maximum likelihood phylogeny of HYL1 and related double-stranded RNA-binding proteins (DRBs) is shown (right), with major taxa colored and key ancestral nodes indicated by grey circles. Ancestral proteins were resurrected, and binding affinities for dsRNA (left) and the *A. thaliana* DCL1 partner were measured by replicate *in vitro* kinetics assays (see Methods). Affinities are reported in –log_10_ units, with longer bars indicating tighter molecular binding, and standard errors are indicated. Dark bars indicate steady-state binding affinity (Kd), and light bars indicate initial binding rates (Km). Yellow star indicates that results are shown for the alternative ancFlowering HYL1 protein, with ambiguous Serine residues replaced by plausible alternative amino-acids (see Supplementary Information Figure S3 for details).

Plant HYL1 has been shown to mediate interactions with double-stranded RNA (dsRNA) and with the twin C-terminal double-stranded RNA-binding motifs (DSRMs) of DCL1 to facilitate RNA interference (Hiraguri et al. 2005; Kurihara et al. 2006; Yang et al. 2010; Yang et al. 2014). To determine when these interactions evolved, we used an alignment-integrated approach (Aadland and Kolaczkowski 2020) to reconstruct maximum-likelihood ancestral DRB protein sequences at key nodes on the HYL1 phylogeny and measured their affinity for dsRNA and for the C-terminal DSRM+DSRM domains of *Arabidopsis thaliana* DCL1 *in vitro* (see Methods). Ancestral protein sequences were reconstructed with high statistical support and low ambiguity (Supplementary Information Figure S1). For each sequence, >84% of reconstructed sites had posterior probability >0.95. In all cases except for the ancestral plant DRB, <1.1% of sites had an alternative reconstruction with posterior probability >0.3, and ancPlant DRB had <2% such sites.

We found that the ancestral plant DRB had relatively low affinity for both dsRNA and for *A. thaliana* DCL1 DSRM+DSRM (Figure 1; Supplementary Information Figure S2). Following the gene duplication giving rise to the HYL1+DRB6 lineage in early plants, ancPlant HYL1+DRB6 increased its affinity for dsRNA by ~9.3-fold and its affinity for *A. thaliana* DCL1 by ~3.5-fold (Figure 1; p<0.0072). After this early functional shift, dsRNA- and DCL1-affinities remained relatively stable across the HYL1 lineage, varying by at most ~2.2-fold, except for the ancestral flowering-plant HYL1, which appeared to have lost affinity for *A. thaliana* DCL1 (~10.9-fold decrease; p<0.008; see Supplementary Information Figure S3). However, the apparent loss of DCL1-affinity in ancFlowering HYL1 is likely due to ancestral reconstruction error; affinity for the *A. thaliana* DCL1 construct was restored when we introduced plausible alternative reconstructions at three positions in the second DSRM of ancFlowering HYL1. All three positions were ambiguously Serine in the maximum-likelihood ancestral sequence (posterior probability <0.57); all three had plausible alternative states with posterior probability >0.3, and in all cases, the plausible alternative residue was conserved in subsequent maximum-likelihood ancestral sequences in the phylogeny. Introducing all three plausible alternative residues increased affinity for *A. thaliana* DCL1 by ~9.9-fold (p<0.012; see Supplementary Information Figure S3).

The initial increase in dsRNA- and DCL1-affinity observed between ancPlant DRB and ancPlant HYL1+DRB6 does appear to be robust to ancestral reconstruction ambiguity. We reconstructed plausible alternative versions of ancPlant DRB and ancPlant HYL1+DRB6 by replacing every maximum-likelihood residue or gap state within the N-terminal DSRM+DSRM region with the next-most-probable residue, provided it had posterior probability >0.3 (see Methods). We observed an ~8.8-fold increase in dsRNA affinity (p<0.007) and an ~4.1-fold increase in affinity for *A. thaliana* DCL1 (p<0.011) between alternative ancPlant DRB and alternative ancPlant HYL1+DRB6, similar to what we observed using maximum-likelihood ancestral sequences (Supplementary Information Figure S4).

Taken together, these results suggest that the ancestral plant HYL1+DRB6 protein increased affinity for dsRNA and for its DCL1 partner immediately after it diverged from the other plant DRB lineages, and that these affinities remained relatively stable across the entire HYL1 lineage. That all ancestral plant HYL1s exhibited relatively high affinity for the derived *A. thaliana* DCL1 DSRM+DSRM construct suggests that the initial HYL1-DCL1 interface may have evolved very early and was subsequently maintained throughout the plant lineage.

### The second DSRM of HYL1 interacts with the first DSRM of DCL1 in vitro

Studies in model systems have shown that the twin N-terminal DSRMs of HYL1 interact directly with the twin C-terminal DSRMs of DCL1 to facilitate RNA interference (Reis et al. 2015). However, the structural interface mediating this interaction is unknown, as is its evolutionary origin. To better understand the HYL1-DCL1 interaction within an evolutionary context, we measured the affinities of individual DSRM pairs from *Arabidopsis thaliana* HYL1-DCL1 and ancestral plant HYL1+DRB6 DSRMs interacting with DSRMs from *A. thaliana* DCL1.

We found that the second DSRM from *A. thaliana* HYL1 had the highest affinity for any *A. thaliana* DCL1 DSRM (p<0.048; Figure 2A; Supplementary Information Figure S5A). Specifically, *A. thaliana* HYL1 DSRM2 had ~3.4-fold higher affinity than DSRM1 for *A. thaliana* DCL1 DSRM1. HYL1 DSRM2 and DSRM1 had equivalent affinity for DCL1 DSRM2 (p>0.21). We observed the same pattern of DSRM-DSRM affinities for ancPlant HYL1+DRB6 domains interacting with *A. thaliana* DCL1 domains (Figure 2B; Supplementary Information Figure S5B). AncPlant HYL1+DRB6 DSRM2 had >5.2-fold higher affinity than DSRM1 for *A. thaliana* DCL1 DSRMs (p<0.022) and ~2.3-fold higher affinity for DCL1 DSRM1, compared to DCL1 DSRM2 (p<0.050).

**Figure 2.**
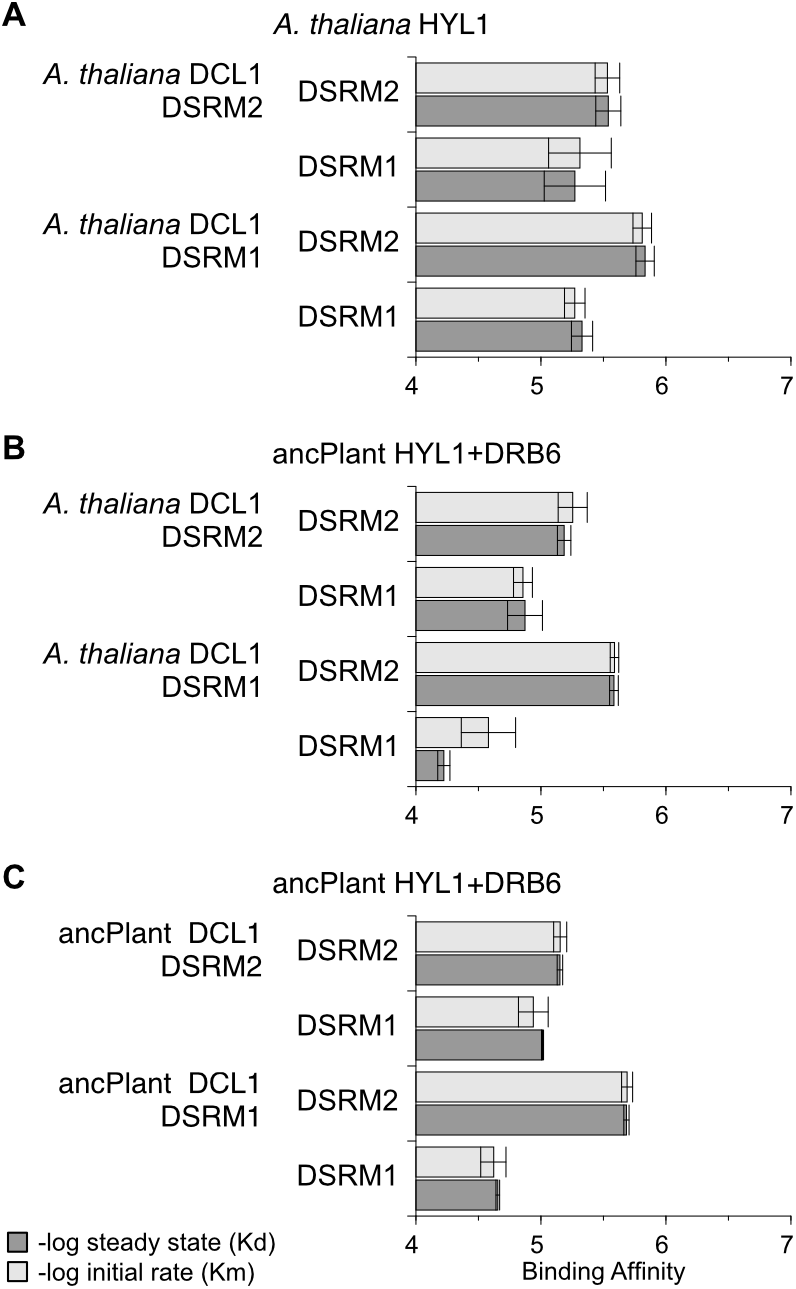
The second DSRM of HYL1 and ancestral plant HYL1+DRB6 interacts strongly with the first DSRM of plant DCL1 *in vitro*. We measured the binding affinities of individual DSRM domains from *Arabidopsis thaliana* HYL1 (A), and ancestral plant HYL1+DRB6 (B,C), interacting with individual DSRM domains from *A. thaliana* DCL1 (A,B) and the ancestral plant DCL1 (C), using an *in vitro* kinetics assay (see Methods). Affinities are reported in –log_10_ units, with longer bars indicating tighter molecular binding, and standard errors are shown. Dark bars indicate steady-state binding affinity (Kd), and light bars indicate initial binding rates (Km).

We reconstructed maximum-likelihood ancestral plant DCL1 DSRM1 and DSRM2 from a consensus phylogeny of plant DCL1 (Supplementary Information Figure S6). DCL1 DSRMs were reconstructed with high confidence; >83% of sites had posterior probability >0.9, and <0.73% of sites had plausible alternative reconstructions (Supplementary Information Figure S6). Ancestral plant HYL1+DRB6 DSRMs exhibited the same affinity profiles for ancPlant DCL1 DSRMs that we observed for DCL1 DSRMs from *A. thaliana* (Figure 2C; Supplementary Information Figure S5C). The ancestral plant HYL1+DRB6 DSRM2 had the highest affinity for any ancestral plant DCL1 DSRM overall, with ~3.4-fold higher affinity for ancPlant DCL1 DSRM1 than any other combination of DSRM-DSRM interactions (p<0.023). Together, these results suggest a specific HYL1-DCL1 interaction mediated by HYL1’s DSRM2 domain and DCL1’s DSRM1 domain may have evolved very early in the HYL1+DRB6 lineage and was maintained over the evolutionary history of land plants.

To further evaluate this hypothesis, we inferred structural models of various HYL1-DCL1 DSRM-DSRM complexes by homology modeling, optimized protein-protein interactions using short molecular dynamics simulations and predicted DSRM-DSRM binding affinities using a structure-based statistical machine learning approach (see Methods). Structure-based affinity predictions were consistent with a model in which HYL1 DSRMs interact with DCL1’s DSRM1 via two main contact regions (Supplementary Information Figures S7,S8). First, the HYL1 DSRM α1-β1 loop appears to interact favorably with parts of DCL1 α1 and α2 via a largely-conserved network of hydrophobic and polar interactions (Supplementary Information Figure S8). Similarly, a conserved network of favorable contacts was observed between regions of HYL1 β1 and β3 interacting with DCL1’s α2 and β2-3 loop (Supplementary Information Figure S8).

Overall, DSRMs from ancestral plant HYL1+DRB6 and *A. thaliana* HYL1 bound DCL1 DSRM1s with >2.36-fold higher predicted affinities than other structural combinations (p<1.4e^−6^; Supplementary Information Figure S7). Affinity prediction was not able to distinguish whether HYL1 DSRM1 or DSRM2 had higher affinity for DCL1 DSRM1 (p=0.71), which is not unexpected, given the relatively low accuracy of structure-based protein-protein affinity prediction (Dias and Kolazckowski 2015; Dias and Kolaczkowski 2017). Although other structural complexes typically had much lower predicted affinities (Supplementary Information Figure S7), we did observe two cases in which alternative HYL1-DCL1 complexes had high affinities (p>0.20). The ancestral plant DCL1 DSRM2 had relatively high predicted affinities for *A. thaliana*’s HYL1 DSRM1 and DSRM2 (Supplementary Information Figure S7). We note that these predicted high-affinity interactions only occur in complexes of ancestral DCL1 DSRM2 and extant *A. thaliana* HYL1 DSRMs and not in ancestral-ancestral or extant-extant complexes, suggesting they are probably not indicative of a long-term functional HYL1-DCL1 interface.

Together, these results suggest that HYL1 may bind DCL1 DSRM1 through an evolutionarily conserved structural interface that likely evolved very early in plants.

### Ancestral HYL1 partially rescues HYL1^−^ knockout phenotype

To examine the potential of ancestral-reconstructed HYL1 to function *in vivo*, we created replicate transgenic *A. thaliana* stably expressing the ancestral flowering plant HYL1 (hereafter, ancFpHYL1), with the native HYL1^−^ gene knocked out (see Methods). On average, ancFpHYL1 was expressed ~5-fold higher in transgenic *A. thaliana* plants, compared to wild-type HYL1 gene expression (p=0.02); HYL1 expression was not detected in the HYL1^−^ knockout (Figure 3A). Confocal imaging of seedling hypocotyl cells suggested that an ancFpHYL1+GFP fusion protein appears to accumulate in the nucleus, similar to native HYL1 (Vazquez et al. 2004), suggesting it could interact with the native *A. thaliana* DCL1 protein (Figure 3B). Visual inspection of ancFpHYL1, HYL1^−^ knockout and wild-type *A. thaliana* seedlings, rosettes, mature plants and mature siliquae suggested that, in general, ancFpHYL1-expressing plants appeared more similar to wild-type HYL1 than to HYL1^−^ knockout plants, which display the characteristic ‘hyponastic leaf’ phenotype (Lu and Fedoroff 2000) and reduced growth rates (Figure 3C-F).

**Figure 3.**
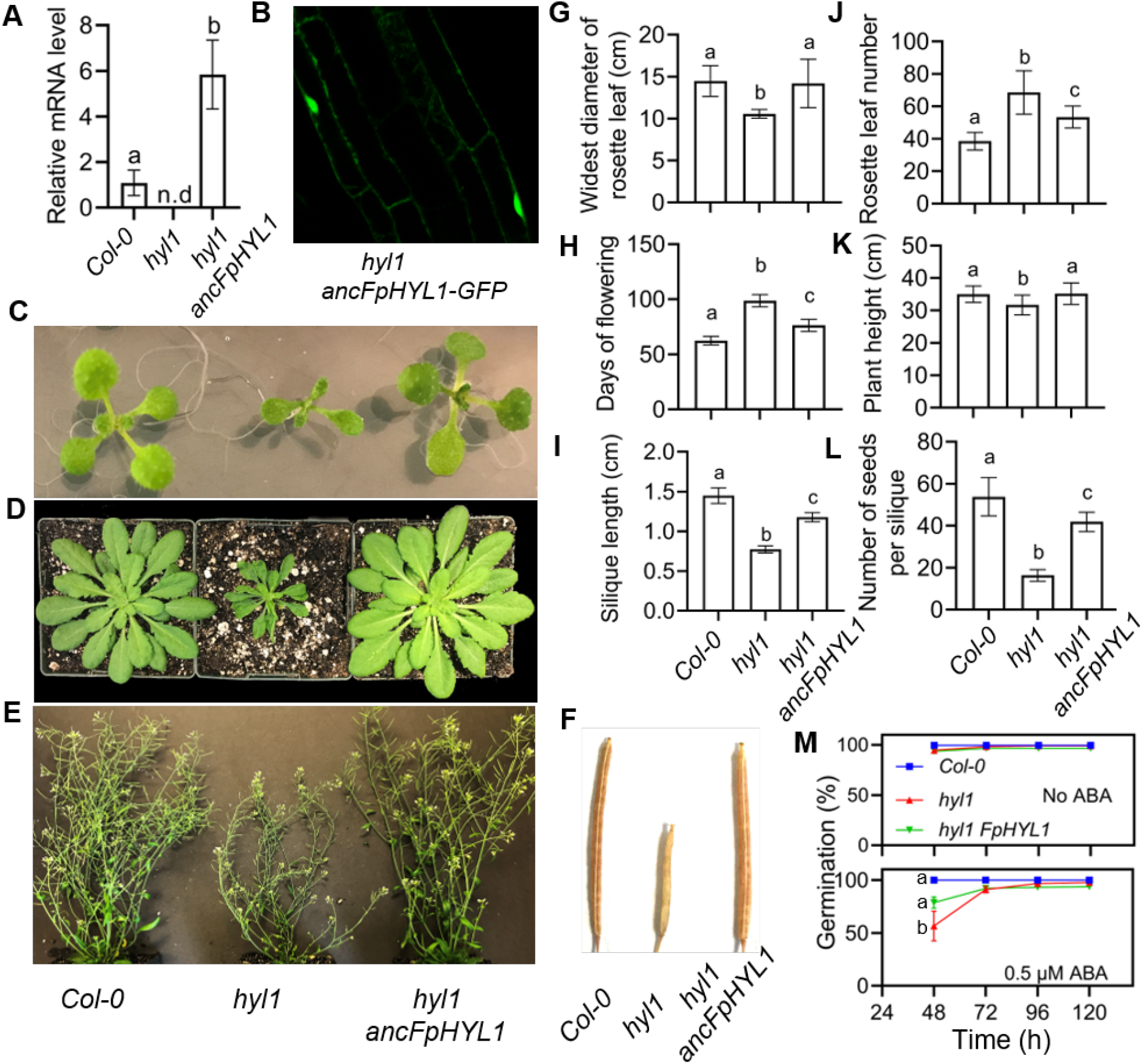
Transgenic expression of ancestral flowering-plant HYL1 (ancFpHYL1) rescues many aspects of the HYL1^−^ knockout phenotype in *Arabidopsis thaliana*. We transformed replicate HYL1^−^ knockout *A. thaliana* plants to express the ancestral flowering-plant HYL1 (ancFpHYL1) as a transgene and compare various aspects of plant phenotype across wild-type (Col-0), HYL1^−^ knockout (hyl1) and ancFpHYL1 plants. **A.** HYL1 transcript levels from 11-day-old seedlings. Statistical differences were evaluated by 2-tailed unpaired *t* test, assuming unequal variances. **B.** Confocal image of root cells from young seedlings expressing an ancFpHYL1-GFP fusion protein. **C.** Representative seedlings of wild-type Col-0 (left), HYL1^−^ knockout (middle) and ancFpHYL1 (right); **D** shows representative plants at the rosette stage, and **E** shows representative mature plants. **F.** Representative siliquae from wild-type Col-0 (left), HYL1^−^ knockout (middle) and ancFpHYL1 (right) plants. **G.** Quantification of rosette diameter. **H.** Comparison of average days to flowering. **I.** Average silique length. **J.** Average number of leaves in rosette stage plants. **K.** Comparison of mature plant heights. **L.** Average number of seeds per silique. **M.** Cold-stratified seeds germinated in the absence (top) or presence of abscisic acid (ABA; bottom) on MS media. We report the average and standard-error in %-germination at each time point. Statistical significance was assessed in (G-L) and at 48h in (M) using one-way ANOVA, Tukey’s multiple comparisons test and Dunnett’s multiple comparisons test. Different letters (a,b,c) indicate statistically significant differences at p<0.05.

The ancFpHYL1 transgene either completely or partially recovered many key quantitative aspects of the HYL1^−^ knockout phenotype (Figure 3G-M). Average rosette leaf diameter was severely reduced in the HYL1^−^ knockout, compared to wild-type (p<1.0e^−4^; Figure 3G) but was completely rescued by ancFpHYL1 (p=0.84). Similarly, plant height was reduced in HYL1^−^ (p=0.01; Figure 3K) but recovered by ancFpHYL1 (p=0.98). Other aspects of the HYL1^−^ knockout phenotype were partially recovered by ancFpHYL1, with quantitative phenotypes having intermediate values between HYL1^−^ and wild-type, including rosette leaf number (p<1.0e-^4^; Figure 3J), days to flowering (p<1.0e^−4^; Figure 3H), silique length (p<1.0e^−4^; Figure 3I) and seeds/silique (p<1.0e^−4^; Figure 3L). The ancFpHYL1 transgene also partially rescued HYL1^−^ hypersensitivity to abscisic acid (Vazquez et al. 2004), which reduced time to seed germination in the HYL1^−^ knockout but had a lesser effect on ancFpHYL1 seeds (p=0.021 at 48 hours; Figure 3M).

That ancFpHYL1 appears to function reasonably well within the modern *A. thaliana* genomic context is surprising, given the relatively high degree of sequence dissimilarity between ancFpHYL1 and the native HYL1 protein (Supplementary Information Figure S9). We observed 92 amino-acid differences between ancFpHYL1 and *A. thaliana* HYL1 out of 392 aligned sequence positions, 19 of which were considered radical substitutions changing biochemical classes (Supplementary Information Figure S9). Thirteen substitutions (5 radical) occurred within HYL1 DSRM1, and 15 (4 radical) were within the DSRM2 domain. In addition, we observed five large deletions in the C-terminal region of *A. thaliana* HYL1, relative to ancFpHYL1, which encodes no annotated structural or functional domains.

### Ancestral HYL1 rescues native microRNA production in HYL1^−^ knockout plants

To better understand how the ancestral flowering-plant HYL1 (ancFpHYL1) functions in a modern genomic context, we quantified global miRNA production in wild-type, HYL1^−^ knockout and transgenic *A. thaliana* plants expressing ancFpHYL1, using small-RNA sequencing (see Methods). Although the total numbers of small-RNA sequencing reads obtained from wild-type, HYL1^−^ knockout and ancFpHYL1 plants was consistent after quality filtering (p=0.41), the number of reads mapping to annotated *A. thaliana* miRNAs was severely reduced in the HYL1^−^ knockout, compared to wild-type and ancFpHYL1 plants (>5.8-fold less; p<1.21e^−5^; Supplementary Information Figure S10), consistent with a model in which HYL1^−^ knockout reduces the efficiency of miRNA production (Hiraguri et al. 2005; Kurihara et al. 2006; Dong et al. 2008; Yang et al. 2010; Yang et al. 2014).

Plant miRNA biogenesis is a complex process that produces a number of intermediate products that may not necessarily function directly in RNAi, including pri-miRNA transcripts, pre-miRNA hairpins and mature miRNA duplexes; only the mature single-stranded miRNA guide strand loaded onto AGO can be assumed to potentially function in RNAi directly (Hutvagner and Simard 2008; Sheu-Gruttadauria and MacRae 2017). To determine which specific miRNA products are being captured by our small-RNA sequencing protocol, we first mapped sequencing reads directly to pri-miRNA transcripts, which are typically the first RNA products produced via transcription of genomic pri-miRNA genes. In all cases, we found that the vast majority of reads mapped to the annotated mature miRNA strands embedded within each pri-miRNA, with very few reads mapping to intervening hairpin regions or other parts of the pri-miRNA (Supplementary Information Figure S11). Over 76% of pri-miRNA transcripts had >80% of sequencing reads mapping to annotated mature miRNA regions within the pri-miRNA sequence (>70% of pri-miRNAs had >90% of reads mapping to mature miRNA regions; Supplementary Information Figure S12). These results suggest that our small-RNA sequencing protocol is likely to be overwhelmingly sequencing mature miRNAs which could function directly in RNAi, with very few reads coming from primary miRNA transcripts or pre-miRNA hairpins.

To determine if our small-RNA sequencing approach was sequencing miRNA duplexes or primarily the mature single-stranded guide strand, we calculated the proportion of reads mapping to each pri-miRNA transcript that matched each of the two possible miRNA guide strands. If small-RNA sequencing is primarily sequencing miRNA duplexes, we would expect the density of read proportions to have a strong peak at 0.5. Alternatively, a peak density skewed toward 1.0 would indicate sequencing of primarily miRNA guide strands. The observed density of read proportions had a strong mode at 0.855±7.703e^−03^ (median 0.888), suggesting very strong strand bias indicative of sequencing reads coming overwhelmingly from mature single-stranded miRNA guide strands with the potential to be directly loaded onto AGO proteins to function in RNAi (Supplementary Figure S13). Although many plant miRNAs exhibit guide-strand selection preference, in which one of the miRNA duplex strands is preferentially loaded onto AGO, in most cases strand selection preference is not absolute (Rajagopalan et al. 2006; Takeda et al. 2008; Eamens et al. 2009). To further evaluate the extent to which our small-RNA sequencing protocol was sequencing mature miRNA guide strands, we calculated miRNA guide strand / pri-miRNA read proportions for those pri-miRNAs with only one annotated mature miRNA guide strand, indicative of cases in which strand-selection preference is very strongly biased. In these cases, nearly all the small-RNA sequencing reads mapping to each pri-miRNA mapped to the single annotated mature miRNA guide strand, further suggesting that the vast majority of sequencing reads were coming from mature single-stranded miRNA guide strands with the strong potential to be functioning in RNAi (Supplementary Information Figure S13).

Previous studies have shown that HYL1 may affect the distribution of miRNA lengths produced by the HYL1-DCL1 complex (Dong et al. 2008). To examine the impact of HYL1 on miRNA fidelity, we locally-aligned all reads mapping to each specific annotated mature miRNA and characterized the variation in miRNA lengths at both the 5’ and 3’ ends (see Methods). We observed no strong differences in 5’ or 3’ miRNA fidelity in the HYL1^−^ knockout, compared to wild-type plants (FDR-corrected K-S and t-test p>0.72). Transgenic ancFpHYL1 plants generated only one miRNA (miR408-5p) with significantly different 5’ or 3’ distributions, compared to wild-type HYL1 (FDR-corrected p<0.02). These results suggest that HYL1^−^ knockout and ancFpHYL1 has little, if any, impact on the length fidelity of miRNA production.

To quantify the impact of the HYL1^−^ knockout and ancFpHYL1 transgene on *A. thaliana* miRNA production, we compared the normalized ‘expression’ of each annotated miRNA from HYL1^−^ and ancFpHYL1 genotypes to wild-type expression (see Methods). We found that ancFpHYL1 miRNA expression profiles were extremely correlated with wild-type HYL1 miRNA expression (*r^2^*=0.998), while HYL1^−^ miRNA expression was only weakly correlated with wild-type miRNA expression (*r^2^*=0.337; Figure 4A). As expected, knocking out HYL1 resulted in differential expression of a number of annotated miRNAs, compared to wild-type plants (Figure 4B). At a false discovery rate (FDR) of 0.05, 37 annotated miRNAs were differentially expressed in HYL1^−^ knockout plants vs wild-type, with 18 being up-regulated and 19 being down-regulated (Supplementary Information Tables S1,S2). Eighteen of these differentially-expressed miRNAs had a fold-change in expression >3 (10 up-regulated; 8 down-regulated). At FDR=0.01, 12 miRNAs were up-regulated in HYL1^−^, and 13 were down-regulated. Consistent with the observed impact of HYL1^−^ knockout on the *A. thaliana* phenotype (see Figure 3), we found that a number of the annotated miRNAs identified as differentially-expressed between HYL1^−^ and wild-type plants had developmental-related functional annotations in miRbase (Kozomara and Griffiths-Jones 2011). For example, miR159a, miR165a, miR168b and miR172c target mRNAs that regulate meristem initiation and development, and miR160a, miR167d, miR170-5p and miR396a/b are associated with the regulation of plant growth hormones auxin and gibberellin. In contrast, ancFpHYL1 had only one annotated miRNA that was identified as differentially-expressed when compared to wild-type HYL1 (miR845a; FDR-corrected p=4.81e^−5^; Figure 4B); this miRNA targets retrotransposons, specifically in pollen (Borges et al. 2018). These results suggest that ancFpHYL1 is sufficient to recover near-native miRNA production when expressed in *A. thaliana.*

**Figure 4.**
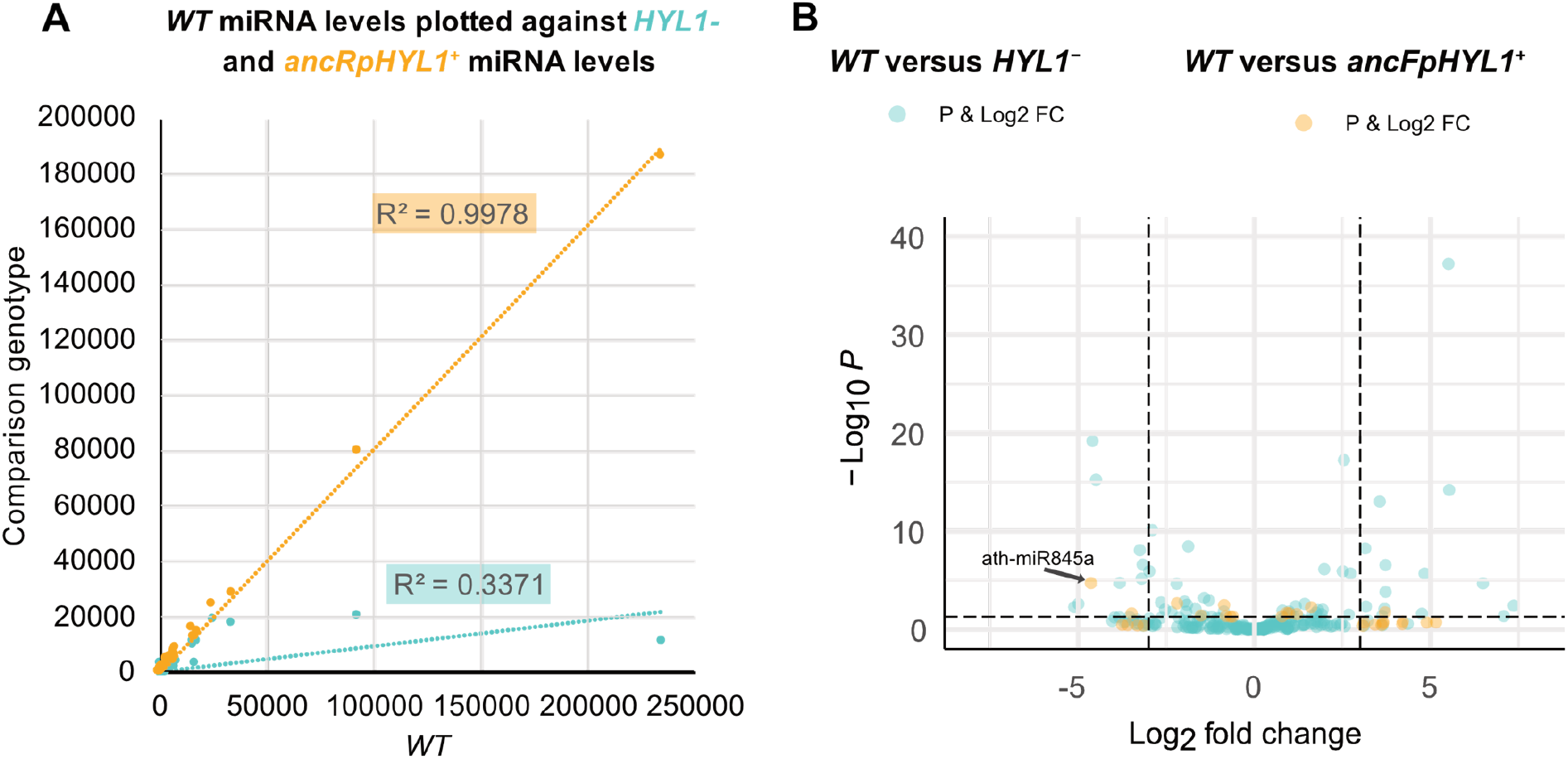
Transgenic expression of ancestral flowering-plant HYL1 (ancFpHYL1) recovers near-native production of annotated miRNAs in HYL1^−^ knockout *Arabidopsis thaliana*. We transformed replicate HYL1^−^ knockout *A. thaliana* plants to express the ancestral flowering-plant HYL1 (ancFpHYL1) as a transgene and measured miRNA production using small-RNA sequencing (see Methods). **A.** We plot the expression of each annotated miRNA (as transcripts-per-million reads, TPM) in HYL1^−^ knockout (teal) and ancFpHYL1 (orange) plants, vs wild-type Col-0 (WT, x-axis). Best-fit linear regressions are shown. **B.** We plot the Log_2_ fold-change (x-axis) and −Log_10_ p-value (y-axis) of each annotated miRNA expressed in HYL1^−^ (teal) and ancFpHYL1 (orange) plants, compared to wild-type Col-0 (WT). Points above the horizontal dotted line line have p<0.05, and points outside the two vertical dotted lines have an absolute fold-change >3.0. The microRNA, ath-miR845a (labeled), is the only differentially-expressed miRNA in ancFpHYL1 plants, compared to wild-type.

## CONCLUSION

Investigations into the evolution of animal and plant development have highlighted the importance of gene regulation for generating phenotypic diversity (Quattrocchio et al. 1999; Graham et al. 2000; Davidson and Erwin 2006; He and Deem 2010; Mansfield 2013; Hudry et al. 2014; Jill Harrison 2017: 2019; Szövényi et al. 2019), but also characterized a specific pattern of evolutionary dynamics that may underlie developmental evolution, in general. Interestingly, the basic transcription-factor driven gene regulatory networks underlying development are shockingly conserved across millions of years of evolutionary history (Peter and Davidson 2011), leading researchers to propose a model in which ‘evolutionary tinkering’ reused modular gene regulatory networks in new ways over time, contributing to both similarities and differences across species (Jacob 1977; Halder et al. 1995; de Mendoza et al. 2013; Paixão-Côrtes et al. 2015; Verd et al. 2019).

Our examination of the evolution of the HYL1-DCL1 partnership responsible for the production of miRNAs contributing to post-transcriptional gene regulation and plant development suggests the HYL1-DCL1 interaction arose very early in plant evolutionary history, possibly as early as the evolution of plant multicellularity but certainly by the time of land plant origins. Ancestral HYL1 proteins reconstructed from before the divergence of mosses from seed plants ~500 Ma (Morris et al. 2018) had high affinity for dsRNA targets and for modern *A. thaliana* DCL1 *in vitro*, and the predicted HYL1-DCL1 structural interface was the same in ancestral HYL1-DCL1 and *A. thaliana* HYL1-DCL1. Surprisingly, the ancestral flowering-plant HYL1 from ~200 Ma (Silvestro et al. 2021) was sufficient to recover many aspects of the HYL1^−^ knockout phenotype in *A. thaliana* by driving near-native miRNA production. These results suggest that the HYL1-DCL1 partnership was strongly conserved, once it evolved in early plants.

Overall, the evolution of plant miRNA production appears consistent with a ‘conserved developmental toolkit’ model, in which miRNA production - facilitated by HYL1-DCL1 - arose as a new mechanism for controlling plant development that may have contributed to key aspects of early plant phenotypic novelty. Other protein families involved in miRNA biogenesis also appear strongly conserved across species (Murphy et al. 2008), further supporting the general conclusion that miRNA-based RNA interference might represent a conserved gene-regulatory ‘toolkit’. In contrast to the relative stability of the proteins involved in RNAi, pri-miRNA genes appear to be very rapidly-evolving. Although recent studies have suggested that some miRNAs might be shared across diverse species (Arteaga-Vázquez et al. 2006), many pri-miRNA genes have very limited taxonomic distributions and appear to be relatively fast-evolving, suggesting that miRNAs and their regulatory targets might be highly evolutionarily labile (Tarver et al. 2012; Chorostecki et al. 2017; Simkin et al. 2020). Through their role in determining RNAi ‘target specificity’ via sequence complementarity with mRNAs, rapid miRNA evolution suggests an efficient mechanism through which evolutionary processes can ‘tinker with’ post-transcriptional gene regulation - and thereby development - by altering pri-miRNA gene transcription, the sequences of the mature miRNAs or specific target sites on mRNAs.

## METHODS

### Protein family identification, sequence alignment and phylogenetic analysis

Protein sequences containing at least one double-stranded RNA-binding motif (DSRM, NCBI conserved domain database id CD00048) were identified by rpsblast search of the nr database using an e-value cutoff of 0.01 (Marchler-Bauer and Bryant 2004; Kozomara and Griffiths-Jones 2011; Marchler-Bauer et al. 2015; NCBI Resource Coordinators 2016). Double-stranded RNA-binding proteins (DRBs) were identified as full-length protein sequences containing 2-3 DSRMs and no other annotated functional domains with e-value <0.01. Dicer and Dicer-like (DCL) protein sequences were identified using a similar approach, with members of the Dicer-DCL protein family being defined as having at least a PAZ domain (NCBI conserved domain database ids CD02843, CD02844 or smart00949) followed by two RIBOc domains (CD00593), with rpsblast e-value <0.01. Other functional domains were annotated by sequence search of the NCBI conserved domain database.

Full-length DRB and Dicer-DCL protein sequences were aligned using mafft-einsi v7.215 (Katoh and Standley 2013), MSAProbs v0.9.7 (Liu et al. 2010), MUSCLE v3.8.31 (Edgar 2004), Probalign v1.4 (Roshan and Livesay 2006) and ProbCons v1.12 (Do et al. 2005), with default parameters. We identified experimentally-determined DSRM structures by sequence search of the RCSB protein data bank (Rose et al. 2013), using DSRMs from annotated human, *D. melanogaster* and *A. thaliana* DRBs and Dicer-DCL sequences as queries and an e-value cutoff of 0.01. Resulting X-ray and NMR structures were aligned using the iterative_structure_align algorithm in MODELLER v9.14 (Eswar et al. 2008; Madhusudhan et al. 2009). We used the mafft --add parameter to align DSRM protein sequences to the structure-based alignment.

Initial maximum likelihood phylogenies were constructed from each alignment using FastTree v2.1.7 with default parameters (Price et al. 2010). Initial trees were used as starting trees for full maximum-likelihood reconstruction using RAxML v8.0.24 (Stamatakis 2014), with the best-fit evolutionary model selected from each alignment using AIC in ProtTest v3 (Darriba et al. 2011). Clade support was evaluated by SH-like aLRT scores (Anisimova and Gascuel 2006).

Ancestral protein sequences were reconstructed using an empirical Bayesian method to integrate over plausible tree topologies (Hanson-Smith et al. 2010). We collected all maximum-likelihood trees inferred from any sequence alignment and estimated the posterior probability of each topology - assuming a given alignment - using Bayes’ rule, assuming a flat prior over topologies. Given a topology and alignment, we inferred the marginal posterior-probability distribution over ancestral sequences at each node using RAxML, which implements an empirical Bayesian ancestral reconstruction algorithm (Yang et al. 1995). We integrated over topologies by weighting each ancestral reconstruction by the posterior probability of that tree, given the alignment. Sequence alignments were mapped to one another using the mafft --merge option. Alignment uncertainty was incorporated by combining ancestral sequence reconstructions from each alignment using a flat prior over alignments (Aadland and Kolaczkowski 2020).

Ancestral insertions and deletions (indels) were reconstructed by converting each sequence alignment to a presence-absence matrix and reconstructing ancestral presence/absence states using the BINCAT model in RAxML, which calculates the posterior probability of a ‘gap’ at each position in the alignment, for each ancestral node on the phylogenetic tree.

### Experimental measurement of protein-RNA and protein-protein affinity

We generated blunt-ended GC-rich 28-base-pair RNA molecules *in vitro* using T7 RNA reverse transcriptase and synthetic dsDNA as template. Complementary purified single-stranded RNAs were annealed to produce double-stranded RNA by combining at 1:1 ratio, heating to 95°C for 5 minutes and then cooling to 25°C. Blunt-ended dsRNA was produced by exposure to alkaline phosphatase. The 3’ end of one RNA strand was biotinylated to facilitate kinetics assays using the Pierce™ 3’ End RNA Biotinylation Kit (Thermo).

Ancestral and extant full-length DRB proteins, DCL DSRM1+DSRM2 constructs and constructs encoding individual DSRM domains were expressed in *E. coli* Rosetta™ 2(DE3)pLysS cells using pET-22b(+) constructs, which were verified by Sanger sequencing. Proteins were purified by His-affinity purification and visualized by SDS-page stained with 1% coomassie. Protein concentrations were measured using a linear-transformed Bradford assay (Zor and Selinger 1996).

We measured protein-RNA and protein-protein binding using a label-free *in vitro* kinetics assay at pH=7 (Abdiche et al. 2008; Frenzel and Willbold 2014). Biotinylated RNA molecules were bound to a series of 8 streptavidin probes for 5 minutes, until saturation was observed. Probes were washed and then exposed to 25 μg/ml biocytin to bind any remaining free streptavidin. FLAG-tagged proteins were similarly immobilized using anti-FLAG affinity probes. Each probe with bound ligand was then exposed to purified protein at increasing concentrations in 1x Kinetics Buffer (ForteBio) for 6 minutes, followed by dissociation in Kinetics Buffer for an additional 4 minutes before exposure to the next concentration of free protein (Frenzel and Willbold 2014). Molecular binding at each concentration over time was measured as the change in laser wavelength when reflected through the probe in solution, sampled every 3 milliseconds. Two probes were not exposed to free protein as controls to evaluate system fluctuation across the time of the experiment; measurements from these control probes were averaged and subtracted from each analysis probe.

For each replicate experiment, we estimated the protein concentration at which ½-maximal steady-state binding was achieved (Kd) by fitting a one-site binding curve to the steady-state laser wavelengths measured across free protein concentrations at saturation, using nonlinear regression. We additionally fit 1-site association/dissociation curves to the full time-course data in order to estimate the initial rates of binding across protein concentrations and used these rates to calculate the protein concentration at which the half-maximal binding rate was achieved (Km). Kds and Kms were –log_10_ transformed to facilitate visualization, and standard errors across 3 experimental replicates were calculated. We calculated the statistical significance of differences between Kds and Kms using the 2-tailed unpaired *t* test, assuming unequal variances.

### DSRM structural modeling and affinity prediction

We modeled individual DSRM-DSRM structural complexes using MODELLER v9.14 (Eswar et al. 2008), using the *A. thaliana* HYL1 DSRM-DSRM complex (PDB ID: 3ADI) as a template (Yang et al. 2010). For each complex, we constructed 100 potential structural models and selected the best 10 using the modeller objective function (molpdf), DOPE and DOPEHR scores (Shen and Sali 2006). Each score was re-scaled to units of standard-deviation across the 100 models, and we ranked models by the best average of re-scaled molpdf, DOPE and DOPEHR scores.

Each initial DSRM-DSRM structural model was used as a starting point for a short molecular dynamics simulation using GROMACS v5.1.2 (Pronk et al. 2013). We used the amber99sb-ildn force field and the tip3p water model. Initial dynamics topologies were generated using the GROMACS pdb2gmx algorithm with default parameters. Topologies were relaxed into simulated solvent using a 50,000-step steepest-descent energy minimization. The system was then brought to 300K using a 50-picosecond dynamics simulation under positional restraints, followed by pressure stabilization for an additional 50 picoseconds. Simulations were run using Particle-Mesh Ewald electrostatics with cubic interpolation and grid spacing of 0.12 nanometers. Van der Waals forces were calculated using a cutoff of 1.0 nanometer. We used Nose-Hoover temperature coupling, with protein and solvent systems coupled separately and the period of temperature fluctuations set to 0.1 picoseconds. Pressure coupling was applied using the Parrinello-Rahman approach, with a fluctuation period of 2.0 picoseconds. Non-bonded cutoffs were treated using buffered Verlet lists. We selected the lowest-energy complex from the last 10 picoseconds of each pressure stabilization simulation for affinity prediction.

DSRM-DSRM affinities were predicted from structural complexes using a statistical machine learning approach (Dias and Kolazckowski 2015). Simulated solvent and ions were excluded from the protein-protein complex, the binding site was identified, and protein-protein interactions were decomposed into a vector of atom-atom interaction features likely to correlate with binding affinity (Dias and Kolazckowski 2015). Affinities (reported as pKd=-log(Kd)) were predicted using a support vector regression model previously trained using a large number of protein-protein complexes with associated experimental affinity measurements (Dias and Kolaczkowski 2017). We report the mean of predicted affinities across the 10 complexes generated from each structural model. Differences in predicted pKds were assessed using the two-tailed unpaired *t* test, assuming unequal variances.

### Creation of transgenic Arabidopsis thaliana plants

Full-length ancestral flowering plant HYL1 (ancFpHYL1) synthesized DNA sequence with *XbaI* and *SalI* sites at the 5’- and 3’-ends was ligated into the *XbaI/SalI* sites of the pCAMBIA1300-Pro35S::GFP plant transformation construct. The ancFpHYL1 transgenic construct was transformed to *Agrobacterium tumefaciens* strain GV3101 and plated on LB media (5 g yeast extract, 10g tryptone and 10g NaCl; pH=7.0) containing 50mg/L Kanamycin and 25mg/L Gentamicin Sulfate. Floral dip method was used for *Agrobacterium*-mediated transformation of *hyl1*-2 mutant *Arabidopsis thaliana* Col-0 plants (Zhang et al. 2006). To select for Hygromycin resistance, sterilized and stratified T1 seeds were plated on MS media (1X Murashige and Skoog salt, 0.05% MES, 1% sucrose and 0.6% phyto Agar) supplemented with 50mg/L Hygromycin and 200mg/L Carbenicillin. Plated seeds were germinated and grown at 22°C for 2 weeks under continuous light. The Hygromycin resistant T1 seedlings were transferred to soil and grown at 26°C with a 16h light/ 8h dark cycle. T2 seedlings were further plated on Hygromycin selection plates to screen for single transgene insertion in the HYL1^−^ knockout mutant background. Replicate T3 and T4 homozygous transgenic seedlings were used for phenotypic analysis and RNA isolation.

Transgenic plants expressing ancFpHYL1 were validated by Reverse Transcription Quantitative PCR (RT-qPCR). Total RNA was extracted from 11-day old WT, HYL1^−^ and ancFpHYL1 T3 homozygous transgenic seedlings following the instructions of the plant miniRNA kit (Zymo research). RT-qPCR was carried out with a Power SYBR Green RNA-to-Ct 1-Step Kit (Applied Biosystems). UBQ10 (AT4G05320) was used as an endogenous control. The 2^−ΔΔCT^ method was used to quantify relative transcript expression (Livak and Schmittgen 2001). Statistical differences in transcript expression were evaluated by the 2-tailed unpaired *t* test, assuming unequal variances.

Transgenic plants expressing ancFpHYL-GFP fusion proteins were confirmed by confocal imaging. GFP signal in the roots of 11-day-old seedlings was imaged by a Zeiss LSM880 (Zeiss Microscopy, Oberkochen, Germany) confocal laser scanning microscope. GFP was excited at 488nm, and emission was collected between 515-550nm.

### Phenotyping Arabidopsis thaliana plants

Flowering time of Col-0 wild-type, HYL1^−^ and transgenic ancFpHYL1 plants was assessed by sowing seeds on soil, stratified at 4°C in darkness for 4 days, and then transferred to 22°C growth chamber under short day conditions (8h light/16h dark) at light intensity of 95 μmol m^−2^ s^−1^. For measuring silique length and seeds per silique, seeds were sowed on soil, stratified at 4°C in darkness for 4 days, and then transferred to 26°C under long day conditions (16h light/8h dark) at light intensity of 75 μmol m^−2^ s^−1^. One hundred siliques were obtained and measured for each genotype, from 3 replicate plant lines. For assaying seed germination in response to exogenous abscisic acid (ABA), seeds were sterilized and plated on ½ MS medium in absence or presence of 0.5 μM ABA, stratified at 4°C in darkness for 4 days, and then transferred to 22°C under continuous low light. We recorded the percentage of 100 seeds that germinated between 48 and 120 hours, at 24-hour intervals. Statistical differences of phenotypes were assessed using one-way ANOVA and Tukey’s multiple comparisons test or Dunnett’s multiple comparisons test.

### Small-RNA sequencing and analysis

Total RNA was extracted from replicate whole 11-day-old Col-0 wild-type, HYL1^−^ knockout and transgenic ancFpHYL1 seedlings using the Monarch Total RNA Miniprep kit (New England BioLabs), following the manufacturer’s instructions. All RNA analytes were assayed for RNA integrity, concentration and fragment size. Samples for total RNA-seq were quantified on a TapeStation system (Agilent, Inc. Santa Clara, CA). Samples with RNA Integrity score (RIN) >8.0 were considered high quality and used in subsequent sequencing. Input concentrations were 65-236 ng/ul. Small RNA-seq library construction was performed using the Perkin Elmer’s Bio Scientific NEXTflex Small RNA-Seq Kit v3 and bar-coded with individual tags following the manufacturer’s instructions (Bioo Scientific Corp, Austin, TX). Libraries were prepared on a Perkin Elmer’s Sciclone G3 NGS Workstation Liquid Handling System. After quality control, sequencing libraries were quantified using a Perkin Elmer’s LabChip system and size-selected on Sage Science’s PIppin Prep. The pool was sequenced on the HS4000 using single-end sequencing of 50 cycles.

The Araport11 reference genome and annotation from The Arabidopsis Information Resource (TAIR) was used for small-RNA sequence mapping (Garcia-Hernandez et al. 2002). After filtering using FastQC with default parameters for quality control, Cutadapt v.2.8 (Martin 2011) was used to remove adapters from the sequencing reads from each of three biological replicates per genotype. Cleaned reads were quasi-mapped to the Araport11 transcriptome using Salmon v.1.1 (Patro et al. 2017), with default parameters. Transcript-level abundance estimates from Salmon were integrated into the DESeq2 v.1.22.2 (Love et al. 2014) pipeline using Tximport v.1.10.1 (Soneson et al. 2016). Differentially-expressed transcripts were identified using DESeq2, which implements the Wald test based on negative binominal generalized linear models. A false discovery rate (FDR) correction (Benjamini and Hochberg 1995) was used to adjust individual p-values for multiple-testing (padj). Genes with padj<0.05 and absolute-value fold change |FC|≥1.5 were considered significantly differentially-expressed. Volcano plots were created using the EnhancedVolcano R package. Micro-RNA sequences were examined for fidelity by calculating the number of aligned reads having extensions or deletions, compared to the annotated mature miRNA. Averaging over replicates for each genotype (Col-0 wild-type, HYL1^−^ knockout and transgenic ancFpHYL1), we calculated the distribution of 3’ and 5’ extensions and deletions for each annotated mature miRNA. Differences in distributions of 3’ and 5’ extensions and deletions across genotypes were assessed using the Kolmogorov-Smirnov (K-S) test.

### Data Availability

All analyses presented in this study were performed using objective, transparent, reproducible algorithms documented in readable source code. All input data, analysis/visualization scripts and results files are freely available under the General Public License (GPL) as open-access documentation associated with this publication, including all sequence alignments, phylogenies, ancestral sequences, structural models and raw kinetics data, at: https://github.com/bryankolaczkowski/AncPlantDRB1. RNASeq data are available at the NCBI Sequence Read Archive (SRA) under bioproject ID: PRJNA608441, submission ID: SUB7022346.

## ACKNOWLEDGEMENTS

This work was supported by the National Science Foundation (Molecular and Cellular Biology, grant number 1817942).

## ABBREVIATIONS

DRB: Double-stranded RNA-binding protein
DCL: Dicer-like protein
DRBM: Double-stranded RNA-binding motif
RNAi: RNA interference
AGO: Argonaute
ASR: Ancestral sequence reconstruction
aLRT: Approximate likelihood ratio test
mRNA: messenger RNA
miRNA: micro RNA
HYL1: Hyponastic leaves 1
DCL1: Dicer-like 1

## REFERENCES

Aadland K, Kolaczkowski B. 2020. Alignment-Integrated Reconstruction of Ancestral Sequences Improves Accuracy. Genome Biol Evol 12:1549–1565.

Abdiche Y, Malashock D, Pinkerton A, Pons J. 2008. Determining kinetics and affinities of protein interactions using a parallel real-time label-free biosensor, the Octet. Anal Biochem 377:209–217.

Agrawal N, Dasaradhi PVN, Mohmmed A, Malhotra P, Bhatnagar RK, Mukherjee SK. 2003. RNA Interference: Biology, Mechanism, and Applications. Microbiol Mol Biol Rev 67:657–685.

Alberti C, Cochella L. 2017. A framework for understanding the roles of miRNAs in animal development. Development 144:2548–2559.

Anisimova M, Gascuel O. 2006. Approximate likelihood-ratio test for branches: A fast, accurate, and powerful alternative. Syst Biol 55:539–552.

Arteaga-Vázquez M, Caballero-Pérez J, Vielle-Calzada J-P. 2006. A Family of MicroRNAs Present in Plants and Animals. The Plant Cell 18:3355–3369.

Axtell MJ, Westholm JO, Lai EC. 2011. Vive la différence: biogenesis and evolution of microRNAs in plants and animals. Genome Biol 12:221.

Benjamini Y, Hochberg Y. 1995. Controlling the False Discovery Rate: A Practical and Powerful Approach to Multiple Testing. Journal of the Royal Statistical Society. Series B (Methodological) 57:289–300.

Billmyre RB, Calo S, Feretzaki M, Wang X, Heitman J. 2013. RNAi function, diversity, and loss in the fungal kingdom. Chromosome Res 21:561–572.

Bollman KM, Aukerman MJ, Park M-Y, Hunter C, Berardini TZ, Poethig RS. 2003. HASTY, the Arabidopsis ortholog of exportin 5/MSN5, regulates phase change and morphogenesis. Development 130:1493–1504.

Bologna NG, Iselin R, Abriata LA, Sarazin A, Pumplin N, Jay F, Grentzinger T, Dal Peraro M, Voinnet O. 2018. Nucleo-cytosolic Shuttling of ARGONAUTE1 Prompts a Revised Model of the Plant MicroRNA Pathway. Molecular Cell 69:709–719.e5.

Borges F, Parent J-S, van Ex F, Wolff P, Martínez G, Köhler C, Martienssen RA. 2018. Transposon-derived small RNAs triggered by miR845 mediate genome dosage response in Arabidopsis. Nature Genetics 50:186–192.

Cerutti H, Casas-Mollano JA. 2006. On the origin and functions of RNA-mediated silencing: from protists to man. Curr Genet 50:81–99.

Chorostecki U, Moro B, Rojas AML, Debernardi JM, Schapire AL, Notredame C, Palatnik JF. 2017. Evolutionary Footprints Reveal Insights into Plant MicroRNA Biogenesis. Plant Cell 29:1248–1261.

Dang Y, Yang Q, Xue Z, Liu Y. 2011. RNA interference in fungi: pathways, functions, and applications. Eukaryot Cell 10:1148–1155.

Darriba D, Taboada GL, Doallo R, Posada D. 2011. ProtTest 3: fast selection of best-fit models of protein evolution. Bioinformatics 27:1164–1165.

Davidson EH, Erwin DH. 2006. Gene regulatory networks and the evolution of animal body plans. Science 311:796–800.

Dias R, Kolaczkowski B. 2017. Improving the accuracy of high-throughput protein-protein affinity prediction may require better training data. BMC Bioinformatics 18:102.

Dias R, Kolazckowski B. 2015. Different combinations of atomic interactions predict protein-small molecule and protein-DNA/RNA affinities with similar accuracy. Proteins 83:2100–2114.

Dias R, Manny A, Kolaczkowski O, Kolaczkowski B. 2017. Convergence of Domain Architecture, Structure, and Ligand Affinity in Animal and Plant RNA-Binding Proteins. Mol Biol Evol 34:1429–1444.

Do CB, Mahabhashyam MSP, Brudno M, Batzoglou S. 2005. ProbCons: Probabilistic consistency-based multiple sequence alignment. Genome Res 15:330–340.

Dong Z, Han M-H, Fedoroff N. 2008. The RNA-binding proteins HYL1 and SE promote accurate in vitro processing of pri-miRNA by DCL1. Proc Natl Acad Sci U S A 105:9970–9975.

Eamens AL, Smith NA, Curtin SJ, Wang M-B, Waterhouse PM. 2009. The Arabidopsis thaliana double-stranded RNA binding protein DRB1 directs guide strand selection from microRNA duplexes. RNA 15:2219–2235.

Edgar RC. 2004. MUSCLE: multiple sequence alignment with high accuracy and high throughput. Nucleic Acids Res 32:1792–1797.

Eswar N, Eramian D, Webb B, Shen M-Y, Sali A. 2008. Protein structure modeling with MODELLER. Methods Mol Biol 426:145–159.

Franco-Zorrilla JM, Valli A, Todesco M, Mateos I, Puga MI, Rubio-Somoza I, Leyva A, Weigel D, García JA, Paz-Ares J. 2007. Target mimicry provides a new mechanism for regulation of microRNA activity. Nat Genet 39:1033–1037.

Frenzel D, Willbold D. 2014. Kinetic Titration Series with Biolayer Interferometry. PLOS ONE 9:e106882.

Garcia-Hernandez M, Berardini TZ, Chen G, Crist D, Doyle A, Huala E, Knee E, Lambrecht M, Miller N, Mueller LA, et al. 2002. TAIR: a resource for integrated Arabidopsis data. Funct Integr Genomics 2:239–253.

Ge W, Chen Y-W, Weng R, Lim SF, Buescher M, Zhang R, Cohen SM. 2012. Overlapping functions of microRNAs in control of apoptosis during Drosophila embryogenesis. Cell Death Differ 19:839–846.

Graham LE, Cook ME, Busse JS. 2000. The origin of plants: body plan changes contributing to a major evolutionary radiation. Proc Natl Acad Sci U S A 97:4535–4540.

Ha M, Kim VN. 2014. Regulation of microRNA biogenesis. Nat Rev Mol Cell Biol 15:509–524.

Halder G, Callaerts P, Gehring WJ. 1995. Induction of ectopic eyes by targeted expression of the eyeless gene in Drosophila. Science 267:1788–1792.

Hammond SM, Bernstein E, Beach D, Hannon GJ. 2000. An RNA-directed nuclease mediates post-transcriptional gene silencing in Drosophila cells. Nature 404:293–296.

Hanson-Smith V, Kolaczkowski B, Thornton JW. 2010. Robustness of Ancestral Sequence Reconstruction to Phylogenetic Uncertainty. Mol Biol Evol 27:1988–1999.

He J, Deem MW. 2010. Hierarchical evolution of animal body plans. Dev Biol 337:157–161.

Hiraguri A, Itoh R, Kondo N, Nomura Y, Aizawa D, Murai Y, Koiwa H, Seki M, Shinozaki K, Fukuhara T. 2005. Specific interactions between Dicer-like proteins and HYL1/DRB-family dsRNA-binding proteins in Arabidopsis thaliana. Plant Mol Biol 57:173–188.

Hudry B, Thomas-Chollier M, Volovik Y, Duffraisse M, Dard A, Frank D, Technau U, Merabet S. 2014. Molecular insights into the origin of the Hox-TALE patterning system. Elife 3:e01939.

Hutvagner G, Simard MJ. 2008. Argonaute proteins: key players in RNA silencing. Nat Rev Mol Cell Biol 9:22–32.

Jacob F. 1977. Evolution and tinkering. Science 196:1161–1166.

Jeseničnik T, Štajner N, Radišek S, Jakše J. 2019. RNA interference core components identified and characterised in Verticillium nonalfalfae, a vascular wilt pathogenic plant fungi of hops. Sci Rep 9:8651.

Jia H, Kolaczkowski O, Rolland J, Kolaczkowski B. 2017. Increased Affinity for RNA Targets Evolved Early in Animal and Plant Dicer Lineages through Different Structural Mechanisms. Mol Biol Evol 34:3047–3063.

Jill Harrison C. 2017. Development and genetics in the evolution of land plant body plans. Philos Trans R Soc Lond B Biol Sci 372.

Katoh K, Standley DM. 2013. MAFFT multiple sequence alignment software version 7: improvements in performance and usability. Mol Biol Evol 30:772–780.

Kim Y-K, Kim B, Kim VN. 2016. Re-evaluation of the roles of DROSHA, Export in 5, and DICER in microRNA biogenesis. Proc Natl Acad Sci U S A 113:E1881–1889.

Kozomara A, Griffiths-Jones S. 2011. miRBase: integrating microRNA annotation and deep-sequencing data. Nucleic Acids Res 39:D152–157.

Kurihara Y, Takashi Y, Watanabe Y. 2006. The interaction between DCL1 and HYL1 is important for efficient and precise processing of pri-miRNA in plant microRNA biogenesis. RNA 12:206–212.

Kurihara Y, Watanabe Y. 2004. Arabidopsis micro-RNA biogenesis through Dicer-like 1 protein functions. Proc Natl Acad Sci U S A 101:12753–12758.

Liu Y, Schmidt B, Maskell DL. 2010. MSAProbs: multiple sequence alignment based on pair hidden Markov models and partition function posterior probabilities. Bioinformatics 26:1958–1964.

Livak KJ, Schmittgen TD. 2001. Analysis of relative gene expression data using real-time quantitative PCR and the 2(-Delta Delta C(T)) Method. Methods 25:402–408.

Love MI, Huber W, Anders S. 2014. Moderated estimation of fold change and dispersion for RNA-seq data with DESeq2. Genome Biology 15:550.

Lu C, Fedoroff N. 2000. A mutation in the Arabidopsis HYL1 gene encoding a dsRNA binding protein affects responses to abscisic acid, auxin, and cytokinin. Plant Cell 12:2351–2366.

Madhusudhan MS, Webb BM, Marti-Renom MA, Eswar N, Sali A. 2009. Alignment of multiple protein structures based on sequence and structure features. Protein Eng Des Sel 22:569–574.

Mansfield JH. 2013. cis-regulatory change associated with snake body plan evolution. Proc Natl Acad Sci U S A 110:10473–10474.

Marchler-Bauer A, Bryant SH. 2004. CD-Search: protein domain annotations on the fly. Nucleic Acids Res 32:W327–331.

Marchler-Bauer A, Derbyshire MK, Gonzales NR, Lu S, Chitsaz F, Geer LY, Geer RC, He J, Gwadz M, Hurwitz DI, et al. 2015. CDD: NCBI’s conserved domain database. Nucleic Acids Res 43:D222–226.

Martin M. 2011. Cutadapt removes adapter sequences from high-throughput sequencing reads. EMBnet.journal 17:10–12.

Martinez J, Patkaniowska A, Urlaub H, Lührmann R, Tuschl T. 2002. Single-stranded antisense siRNAs guide target RNA cleavage in RNAi. Cell 110:563–574.

Martinez NJ, Walhout AJM. 2009. The interplay between transcription factors and microRNAs in genome-scale regulatory networks. Bioessays 31:435–445.

de Mendoza A, Sebé-Pedrós A, Šestak MS, Matejcic M, Torruella G, Domazet-Loso T, Ruiz-Trillo I. 2013. Transcription factor evolution in eukaryotes and the assembly of the regulatory toolkit in multicellular lineages. Proc Natl Acad Sci U S A 110:E4858–4866.

Moran Y, Agron M, Praher D, Technau U. 2017. The evolutionary origin of plant and animal microRNAs. Nat Ecol Evol 1:27.

Morris JL, Puttick MN, Clark JW, Edwards D, Kenrick P, Pressel S, Wellman CH, Yang Z, Schneider H, Donoghue PCJ. 2018. The timescale of early land plant evolution. PNAS 115:E2274–E2283.

Mukherjee K, Campos H, Kolaczkowski B. 2013. Evolution of animal and plant dicers: early parallel duplications and recurrent adaptation of antiviral RNA binding in plants. Mol Biol Evol 30:627–641.

Murphy D, Dancis B, Brown JR. 2008. The evolution of core proteins involved in microRNA biogenesis. BMC Evol Biol 8:92.

NCBI Resource Coordinators. 2016. Database resources of the National Center for Biotechnology Information. Nucleic Acids Res 44:D7–19.

Obbard DJ, Gordon KHJ, Buck AH, Jiggins FM. 2009. The evolution of RNAi as a defence against viruses and transposable elements. Philos Trans R Soc Lond B Biol Sci 364:99–115.

O’Brien J, Hayder H, Zayed Y, Peng C. 2018. Overview of MicroRNA Biogenesis, Mechanisms of Actions, and Circulation. Front Endocrinol (Lausanne) 9:402.

Paixão-Côrtes VR, Salzano FM, Bortolini MC. 2015. Origins and evolvability of the PAX family. Semin Cell Dev Biol 44:64–74.

Patro R, Duggal G, Love MI, Irizarry RA, Kingsford C. 2017. Salmon provides fast and bias-aware quantification of transcript expression. Nature Methods 14:417–419.

Pertea M, Salzberg SL. 2010. Between a chicken and a grape: estimating the number of human genes. Genome Biol 11:206.

Peter IS, Davidson EH. 2011. Evolution of gene regulatory networks controlling body plan development. Cell 144:970–985.

Price MN, Dehal PS, Arkin AP. 2010. FastTree 2--approximately maximum-likelihood trees for large alignments. PLoS One 5:e9490.

Pronk S, Páll S, Schulz R, Larsson P, Bjelkmar P, Apostolov R, Shirts MR, Smith JC, Kasson PM, van der Spoel D, et al. 2013. GROMACS 4.5: a high-throughput and highly parallel open source molecular simulation toolkit. Bioinformatics 29:845–854.

Quattrocchio F, Wing J, van der Woude K, Souer E, de Vetten N, Mol J, Koes R. 1999. Molecular analysis of the anthocyanin2 gene of petunia and its role in the evolution of flower color. Plant Cell 11:1433–1444.

Rajagopalan R, Vaucheret H, Trejo J, Bartel DP. 2006. A diverse and evolutionarily fluid set of microRNAs in Arabidopsis thaliana. Genes Dev 20:3407–3425.

Reis RS, Eamens AL, Roberts TH, Waterhouse PM. 2015. Chimeric DCL1-Partnering Proteins Provide Insights into the MicroRNA Pathway. Front Plant Sci 6:1201.

Romero IG, Ruvinsky I, Gilad Y. 2012. Comparative studies of gene expression and the evolution of gene regulation. Nat Rev Genet 13:505–516.

Rose PW, Bi C, Bluhm WF, Christie CH, Dimitropoulos D, Dutta S, Green RK, Goodsell DS, Prlic A, Quesada M, et al. 2013. The RCSB Protein Data Bank: new resources for research and education. Nucleic Acids Res 41:D475–482.

Roshan U, Livesay DR. 2006. Probalign: multiple sequence alignment using partition function posterior probabilities. Bioinformatics 22:2715–2721.

Ruhfel BR, Gitzendanner MA, Soltis PS, Soltis DE, Burleigh JG. 2014. From algae to angiosperms-inferring the phylogeny of green plants (Viridiplantae) from 360 plastid genomes. BMC Evol Biol 14:23.

Shabalina SA, Koonin EV. 2008. Origins and evolution of eukaryotic RNA interference. Trends Ecol Evol 23:578–587.

Shen M, Sali A. 2006. Statistical potential for assessment and prediction of protein structures. Protein Sci 15:2507–2524.

Sheu-Gruttadauria J, MacRae IJ. 2017. Structural Foundations of RNA Silencing by Argonaute. J Mol Biol 429:2619–2639.

Silvestro D, Bacon CD, Ding W, Zhang Q, Donoghue PCJ, Antonelli A, Xing Y. 2021. Fossil data support a pre-Cretaceous origin of flowering plants. Nature Ecology & Evolution:1–9.

Simkin A, Geissler R, McIntyre ABR, Grimson A. 2020. Evolutionary dynamics of microRNA target sites across vertebrate evolution. PLoS Genet 16:e1008285.

Soneson C, Love MI, Robinson MD. 2016. Differential analyses for RNA-seq: transcript-level estimates improve gene-level inferences. F1000Res 4:1521.

Stamatakis A. 2014. RAxML version 8: a tool for phylogenetic analysis and post-analysis of large phylogenies. Bioinformatics 30:1312–1313.

Szövényi P, Waller M, Kirbis A. 2019. Evolution of the plant body plan. Curr Top Dev Biol 131:1–34.

Takeda A, Iwasaki S, Watanabe T, Utsumi M, Watanabe Y. 2008. The Mechanism Selecting the Guide Strand from Small RNA Duplexes is Different Among Argonaute Proteins. Plant and Cell Physiology 49:493–500.

Tarver JE, Donoghue PCJ, Peterson KJ. 2012. Do miRNAs have a deep evolutionary history? Bioessays 34:857–866.

Vazquez F, Gasciolli V, Crété P, Vaucheret H. 2004. The nuclear dsRNA binding protein HYL1 is required for microRNA accumulation and plant development, but not posttranscriptional transgene silencing. Curr Biol 14:346–351.

Verd B, Monk NA, Jaeger J. 2019. Modularity, criticality, and evolvability of a developmental gene regulatory network. Elife 8.

Wickett NJ, Mirarab S, Nguyen N, Warnow T, Carpenter E, Matasci N, Ayyampalayam S, Barker MS, Burleigh JG, Gitzendanner MA, et al. 2014. Phylotranscriptomic analysis of the origin and early diversification of land plants. Proc Natl Acad Sci U S A 111:E4859–4868.

Willmann MR, Poethig RS. 2007. Conservation and evolution of miRNA regulatory programs in plant development. Curr Opin Plant Biol 10:503–511.

Yang SW, Chen H-Y, Yang J, Machida S, Chua N-H, Yuan YA. 2010. Structure of Arabidopsis HYPONASTIC LEAVES1 and its molecular implications for miRNA processing. Structure 18:594–605.

Yang X, Ren W, Zhao Q, Zhang P, Wu F, He Y. 2014. Homodimerization of HYL1 ensures the correct selection of cleavage sites in primary miRNA. Nucleic Acids Res 42:12224–12236.

Yang Z, Kumar S, Nei M. 1995. A new method of inference of ancestral nucleotide and amino acid sequences. Genetics 141:1641–1650.

Zhang R, Jing Y, Zhang H, Niu Y, Liu C, Wang J, Zen K, Zhang C-Y, Li D. 2018. Comprehensive Evolutionary Analysis of the Major RNA-Induced Silencing Complex Members. Sci Rep 8:14189.

Zhang X, Henriques R, Lin S-S, Niu Q-W, Chua N-H. 2006. Agrobacterium-mediated transformation of Arabidopsis thaliana using the floral dip method. Nat Protoc 1:641–646.

Zhang X, Zhao H, Gao S, Wang W-C, Katiyar-Agarwal S, Huang H-D, Raikhel N, Jin H. 2011. Arabidopsis Argonaute 2 regulates innate immunity via miRNA393(∗)-mediated silencing of a Golgi-localized SNARE gene, MEMB12. Mol Cell 42:356–366.

Zor T, Selinger Z. 1996. Linearization of the Bradford protein assay increases its sensitivity: theoretical and experimental studies. Anal Biochem 236:302–308.

