## Supplemental Information for "Direct molecular evidence for an ancient, conserved developmental toolkit controlling post-transcriptional gene regulation in land plants"

### SUPPLEMENTARY INFORMATION

#### SUPPLEMENTARY TABLES

| | Number of differentially expressed genes FDR<0.05 | | | Number of differentially expressed genes FDR<0.01 | | | Number of differentially expressed genes \|FC\|>3 and FDR<0.05 | | |
|---|---|---|---|---|---|---|---|---|---|
| | Total | Up-regulated | Down-regulated | Total | Up-regulated | Down-regulated | Total | Up-regulated \|FC\|>3 | Down-regulated \|FC\|>3 |
| *WT* vs *hyl1* | 37 | 18 | 19 | 25 | 12 | 13 | 18 | 10 | 8 |
| *WT* vs *hyl1_35S::ancHYL1* | 1 | 1 | 0 | 1 | 1 | 0 | 1 | 1 | 0 |
| *hyl1* vs *hyl1_35S::ancHYL1* | 32 | 17 | 15 | 27 | 16 | 11 | 12 | 9 | 3 |

**Table S1. Transgenic ancFpHYL1 removes differential-expression of miRNAs observed in HYL1$^-$ knockout, compared to wild-type *A. thaliana*.** We report the counts of differentially-expressed annotated miRNAs when comparing wild-type (WT) *A. thaliana* to HYL1$^-$ knockout (hyl1; top line), wild-type vs ancFpHYL1 (hyl1_35S::ancHYL1; middle line), and HYL1$^-$ knockout vs ancFpHYL1 (bottom line).

| | | | mean TPM | | SE TPM | |
|---|---|---|---|---|---|---|
| **miRNA** | **Padj** | **log2FoldChange** | **COL-0 wild-type** | **HYL1$^-$ knockout** | **COL-0 wild-type** | **HYL1$^-$ knockout** |
| ath-miR838 | 2.81E-2 | -5.12 | 7.07 | 0.55 | 1.79E-1 | 7.16E-2 |
| ath-miR169f-3p | 1.54E-2 | -5.00 | 15.18 | 1.00 | 6.84E-1 | 1.29E-1 |
| ath-miR157a-3p | 5.44E-18 | -4.60 | 980.88 | 18.68 | 4.77E+1 | 1.03E+0 |
| ath-miR162a-3p | 2.37E-14 | -4.50 | 1630.73 | 35.41 | 5.34E+1 | 2.21E+0 |
| ath-miR162a-5p | 1.68E-4 | -3.82 | 286.58 | 10.72 | 1.06E+1 | 8.38E-1 |
| ath-miR159a | 1.47E-7 | -3.26 | 235012.91 | 11469.24 | 3.19E+3 | 2.67E+2 |
| ath-miR171b-5p | 6.55E-5 | -3.21 | 96.35 | 5.41 | 4.32E+0 | 5.32E-1 |
| ath-miR173-5p | 3.96E-6 | -3.17 | 2560.84 | 145.36 | 1.16E+2 | 1.26E+1 |
| ath-miR391-5p | 1.35E-5 | -2.98 | 529.86 | 37.98 | 3.45E+1 | 2.74E+0 |
| ath-miR165a-5p | 1.85E-9 | -2.90 | 311.53 | 18.90 | 7.98E+0 | 5.66E-1 |
| ath-miR156d-3p | 1.18E-2 | -2.65 | 97.30 | 11.00 | 4.32E+0 | 1.31E+0 |
| ath-miR160a-3p | 4.56E-2 | -2.54 | 230.61 | 10.49 | 1.23E+1 | 6.69E-1 |
| ath-miR396a-3p | 1.88E-4 | -2.21 | 1895.00 | 177.68 | 7.23E+1 | 8.30E+0 |
| ath-miR164c-3p | 1.02E-2 | -1.98 | 127.99 | 16.07 | 6.51E+0 | 1.31E+0 |
| ath-miR165a-3p | 1.84E-2 | -1.92 | 1625.53 | 193.42 | 6.37E+1 | 2.25E+0 |
| ath-miR824-3p | 7.42E-8 | -1.88 | 691.81 | 89.23 | 2.28E+1 | 3.91E+0 |
| ath-miR472-3p | 4.12E-2 | -1.67 | 336.88 | 48.17 | 1.13E+1 | 1.15E+0 |
| ath-miR168b-3p | 4.67E-3 | -1.43 | 297.26 | 54.33 | 1.26E+1 | 3.15E+0 |
| ath-miR161.1 | 8.11E-3 | -1.30 | 16841.57 | 3203.49 | 5.70E+2 | 1.23E+2 |
| ath-miR841a-3p | 2.45E-2 | 1.17 | 133.45 | 149.64 | 4.59E+0 | 7.56E+0 |
| ath-miR394a | 1.54E-2 | 1.43 | 823.62 | 1145.03 | 3.26E+1 | 7.35E+1 |
| ath-miR167d | 4.90E-2 | 1.80 | 136.62 | 186.38 | 8.59E+0 | 9.86E+0 |
| ath-miR5642a | 4.56E-2 | 1.94 | 39.30 | 81.86 | 2.45E-1 | 2.19E+0 |
| ath-miR390a-5p | 9.62E-6 | 1.98 | 482.71 | 863.37 | 1.68E+1 | 3.72E+1 |

| | | | | | | |
|---|---|---|---|---|---|---|
| ath-miR396b-3p | 1.35E-5 | 2.50 | 1492.65 | 3660.72 | 8.63E+1 | 2.22E+2 |
| ath-miR5654-3p | 3.27E-16 | 2.54 | 170.49 | 431.35 | 7.16E+0 | 1.93E+1 |
| ath-miR393b-3p | 2.06E-5 | 2.72 | 836.60 | 2368.09 | 6.31E+1 | 1.50E+2 |
| ath-miR822-5p | 1.06E-7 | 3.14 | 733.26 | 3385.43 | 3.75E+1 | 2.87E+2 |
| ath-miR845a | 2.52E-2 | 3.16 | 1.98 | 4.43 | 1.30E-1 | 1.62E-1 |
| ath-miR5026 | 2.75E-12 | 3.56 | 76.59 | 451.29 | 4.58E+0 | 3.06E+1 |
| ath-miR833a-3p | 4.09E-2 | 3.69 | 4.74 | 15.77 | 4.23E-1 | 1.34E+0 |
| ath-miR822-3p | 3.99E-6 | 3.72 | 20.89 | 135.71 | 1.11E+0 | 1.14E+1 |
| ath-miR5629 | 1.18E-3 | 3.72 | 5.50 | 30.17 | 4.56E-1 | 2.11E+0 |
| ath-miR850 | 2.06E-5 | 4.83 | 11.42 | 183.83 | 4.69E-1 | 1.15E+0 |
| ath-miR840-5p | 9.80E-36 | 5.51 | 54.73 | 1045.82 | 3.41E+0 | 5.47E+1 |
| ath-miR5654-5p | 2.14E-13 | 5.53 | 3.71 | 78.31 | 2.25E-1 | 5.64E+0 |
| ath-miR865-5p | 1.78E-4 | 6.48 | 1.88 | 15.45 | 1.25E-1 | 8.58E-1 |

**Table S2. List of the 37 annotated miRNAs identified as differentially-expressed in the HYL1$^-$ knockout, compared to wild-type *A. thaliana*.** We report the adjusted p-value (Padj), $log_2$ fold change (log2FoldChange) and mean and standard-error (SE) of transcripts per million reads (TPM) for the COL-0 wild-type and HYL1$^-$ knockout genotypes, averaged over 3 replicate plants, for each of the annotated miRNAs identified as differentially-expressed. Negative fold-change values indicate reduced expression in HYL1$^-$ knockout plants, compared to wild-type.

#### SUPPLEMENTARY FIGURES

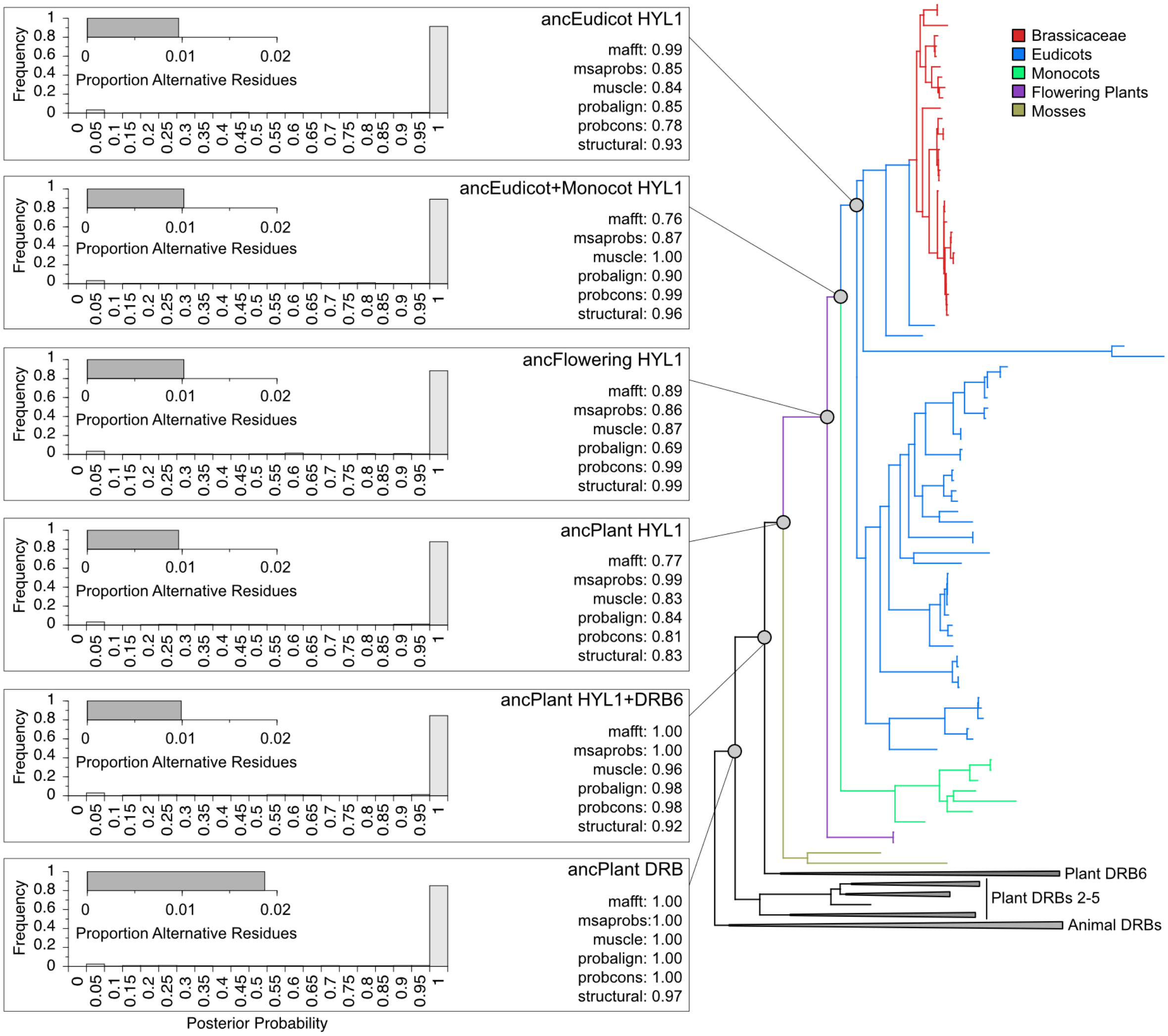


**Figure S1. Plant HYL1 protein family phylogeny and ancestral protein sequences were inferred with high statistical confidence.** We inferred the maximum-likelihood phylogeny of plant HYL1 sequences, rooted with other plant DRBs and animal DRBs, using six different sequence alignments (see Methods). We plot the maximum-likelihood consensus tree across alignments, with specific plant lineages indicated by branch colors. Support for key ancestral nodes (gray circles and corresponding boxes) from each alignment are reported as SH-like aLRT scores, with 1.0 indicating maximal support. We reconstructed maximum-likelihood ancestral sequences at each key node, using the structure-based sequence alignment (see Methods). Column graphs indicate the frequency with which ancestral residues and gap states were reconstructed with posterior probabilities binned every 0.05. Inset bar graph indicates the proportion of residues and gap states for which an alternative reconstruction had posterior probability > 0.3.

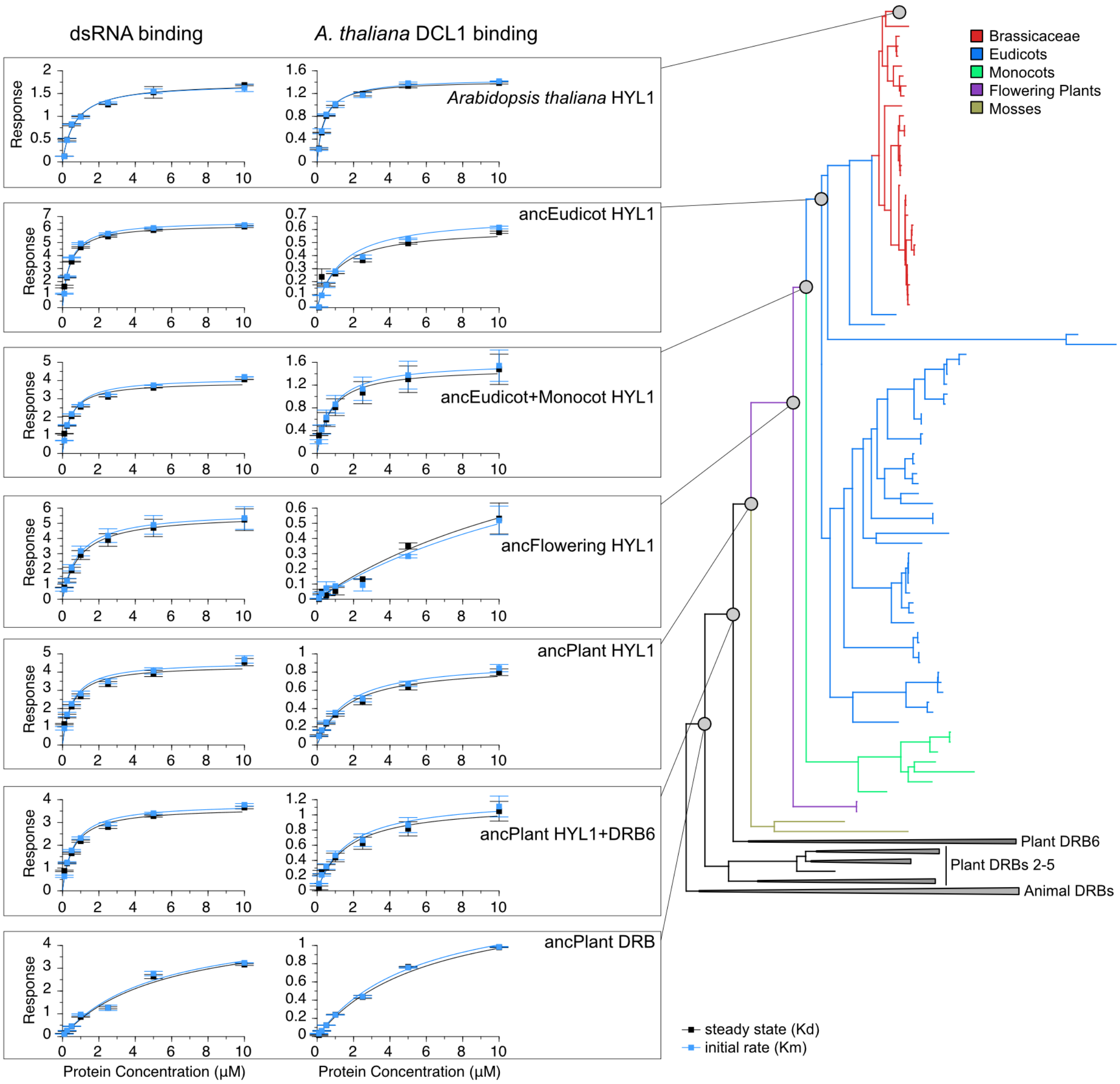


**Figure S2. Plant HYL1 evolved high affinity for double-stranded RNA and for the C-terminal DSRM+DSRM region of *Arabidopsis thaliana* DCL1 early in the plant lineage, before the HYL1-DRB6 duplication.** For each key ancestral node on the plant HYL1 phylogeny (gray circles), we reconstructed maximum-likelihood ancestral protein sequences (see Supplementary Fig. S4) and measured their affinity for dsRNA (left column) and for the C-terminal region of *A. thaliana* DCL1 (right column). We plot protein concentration-response curves for each ancestral protein and ligand and report both steady-state response (Kd; black) and initial binding rate (Km; blue), with bars indicating standard errors over three replicates (see Methods for details). We fit one-site binding curves to each data series using nonlinear regression; corresponding –log-transformed Kd and Km values are shown in Fig. 1, main text.

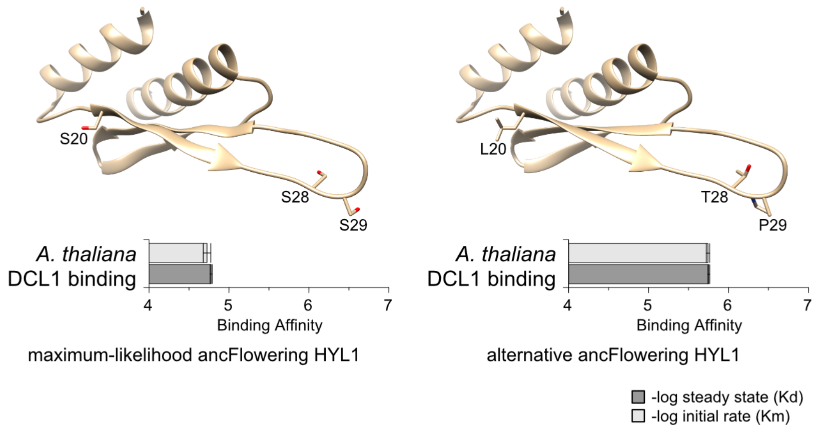


**Figure S3. Plausible alternative ancestral residues in ancFlowering HYL1 DSRM2 restore high affinity for *Arabidopsis thaliana* DCL1.** We plot the locations of three ambiguously-reconstructed Serine residues in the second DSRM of ancFlowering HYL1 (left; modeled using PDB ID 3ADJ). Right panel shows the corresponding alternative residue reconstructions (posterior probability >0.3). Bar graphs indicate the –log-transformed binding affinity of each protein to an *A. thaliana* DCL1 DSRM+DSRM construct (see Methods). Longer bars indicate tighter binding, and standard errors over three replicates are shown. We plot the –log steady-state affinity (Kd; dark bars) and the –log initial binding rate (Km; light bars). See Fig. 1 (main text) for the location of ancFlowering HYL1 on the HYL1 phylogeny.

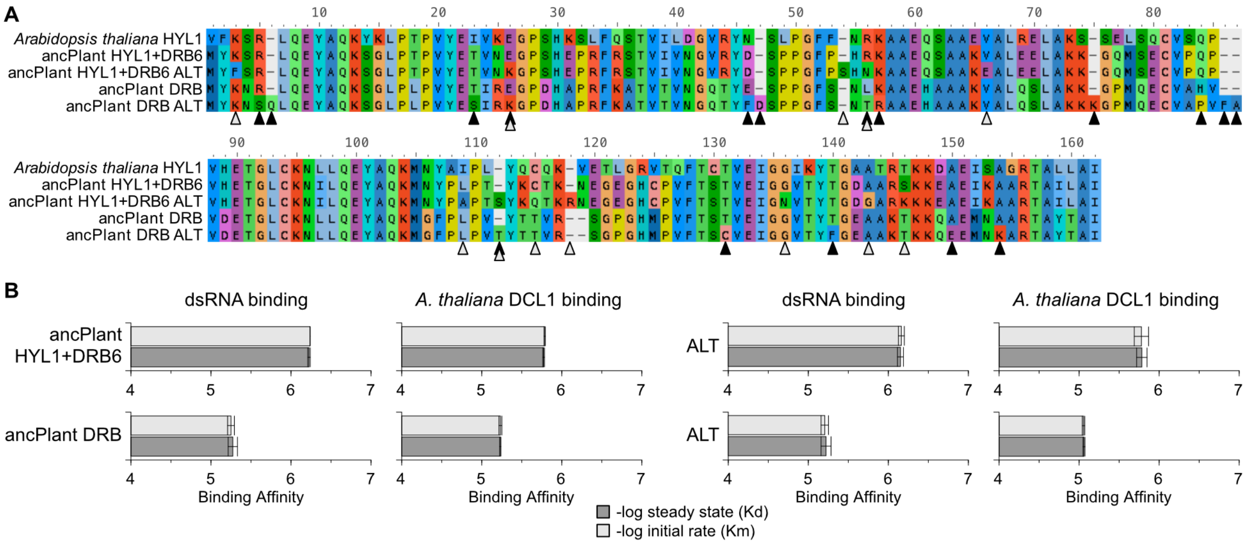


**Figure S4. The increase in affinity for dsRNA and *Arabidopsis thaliana* DCL1 observed between ancPlant DRB and ancPlant HYL1+DRB6 is robust to ancestral sequence reconstruction ambiguity. A.** We plot the plausible alternative reconstructions (ALT) in the DSRM+DSRM region of ancPlant DRB (black triangles) and ancPlant HYL1+DRB6 (gray triangles). Alternative residues and gap states are defined as a non-maximum-likelihood state having posterior probability >0.3. **B.** We measured the affinity of maximum-likelihood (left two columns) and alternative (right two columns) ancestral DRB constructs to dsRNA and *A. thaliana* DCL1 using a label-free kinetics assay (see Methods). Bars indicate the –log-transformed binding affinity, with longer bars indicating tighter binding. Means and standard errors over three replicates are shown for steady-state affinity (Kd; dark bars) and initial binding rate (Km; light bars).

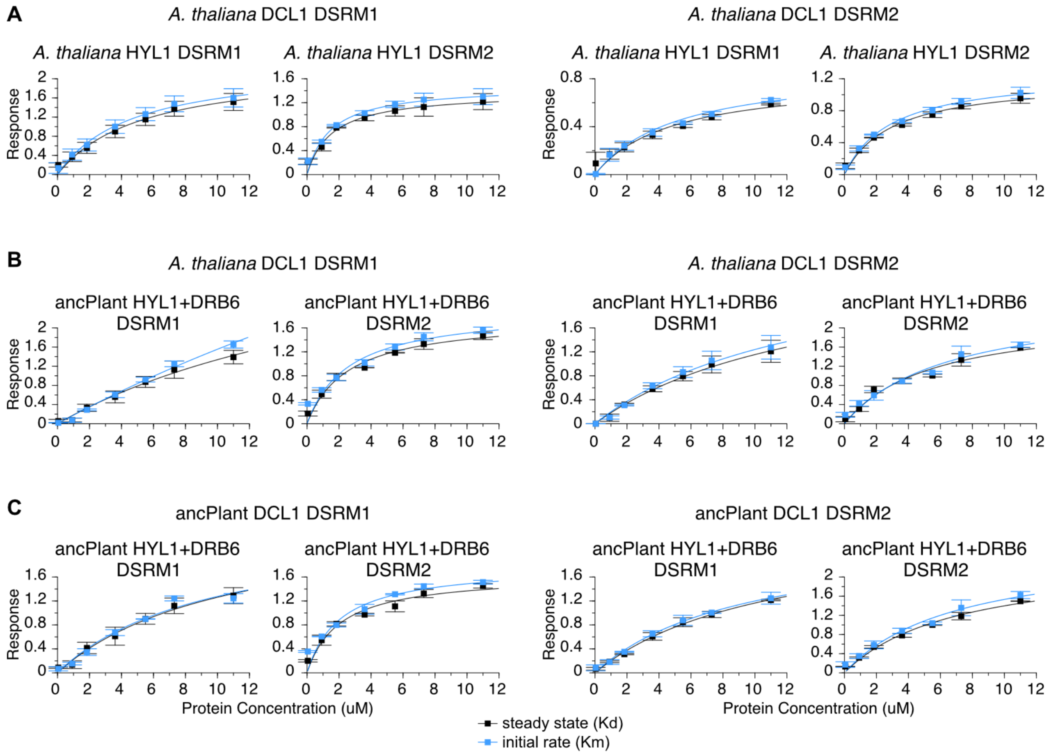


**Figure S5. The second DSRM domain of ancestral plant HYL1+DRB6 and *Arabidopsis thaliana* HYL1 binds tightly to the first DSRM of DCL1.** We measured the *in vitro* binding of individual DSRMs from the N-terminal region of *A. thaliana* HYL1 (**A**) and ancPlant HYL1+DRB6 (**B**) with individual DSRMs from the C-terminal region of *A. thaliana* DCL1. We plot concentration-response curves for each pair of ancestral and *A. thaliana* DSRMs and report both steady-state response (Kd; black) and initial binding rate (Km; blue), with bars indicating standard errors over three replicates (see Methods for details). **C** shows similar analyses of ancPlant HYL1+DRB6 DSRMs interacting with ancestral-reconstructed DCL1 DSRMs (see Supplementary Information Fig. S6). We fit one-site binding curves to each data series using nonlinear regression; corresponding –log-transformed Kd and Km values are shown in Fig. 2, main text.

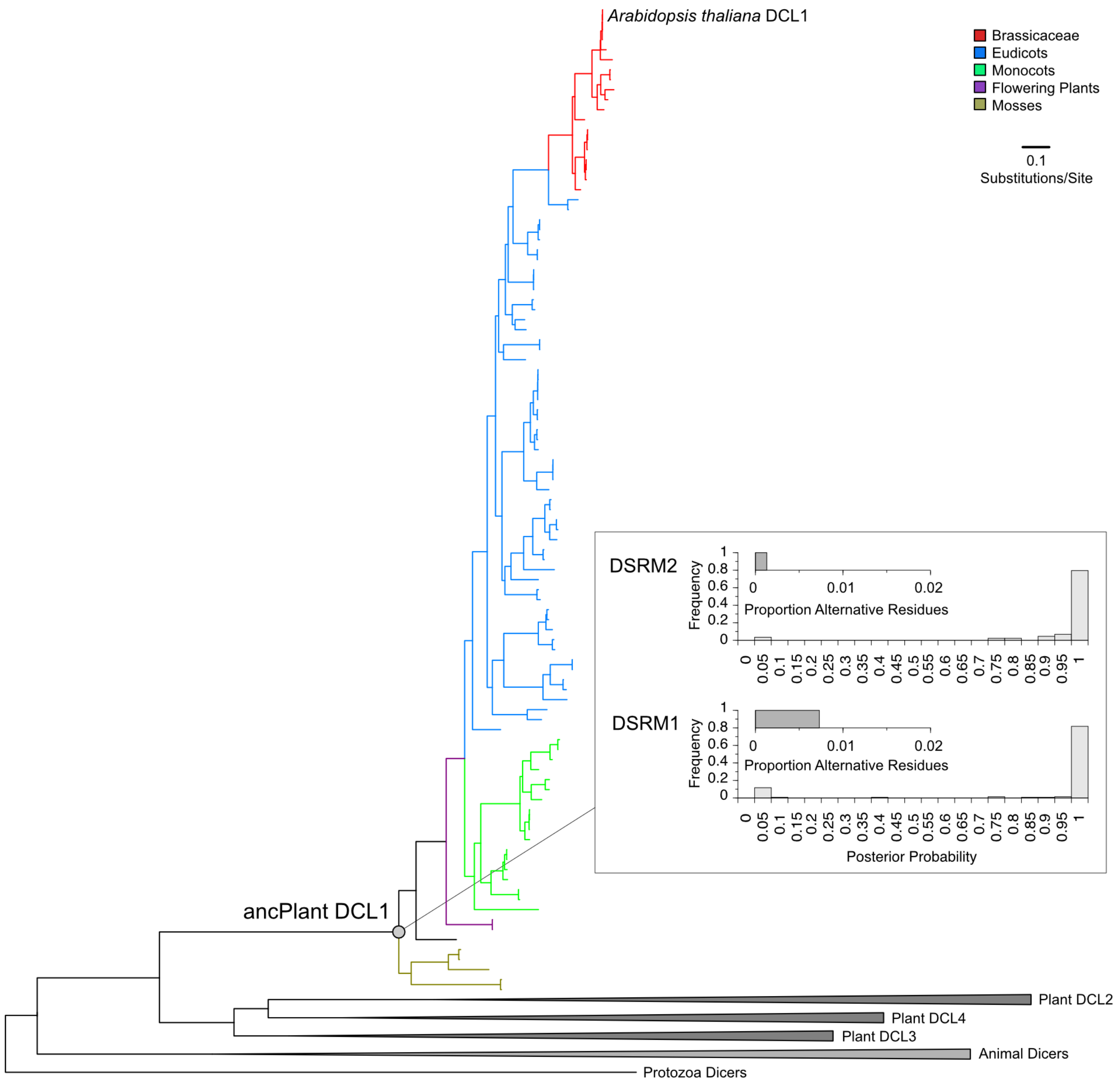


**Figure S6. Plant DCL1 protein family phylogeny and ancestral DSRM sequences were inferred with high statistical confidence.** We inferred the maximum-likelihood phylogeny of plant DCL1 sequences, rooted with other plant DCLs and animal Dicers. We plot the maximum-likelihood consensus tree across different sequence alignments, with specific plant lineages indicated by branch colors. We reconstructed maximum-likelihood ancestral sequences for the C-terminal DSRM1 and DSRM2 functional domains of the last common DCL1 ancestor (gray circle), using a structure-based sequence alignment (see Methods). Column graphs indicate the frequency with which ancestral residues and gap states were reconstructed with posterior probabilities binned every 0.05. Inset bar graph indicates the proportion of residues and gap states for which an alternative reconstruction had posterior probability >0.3.

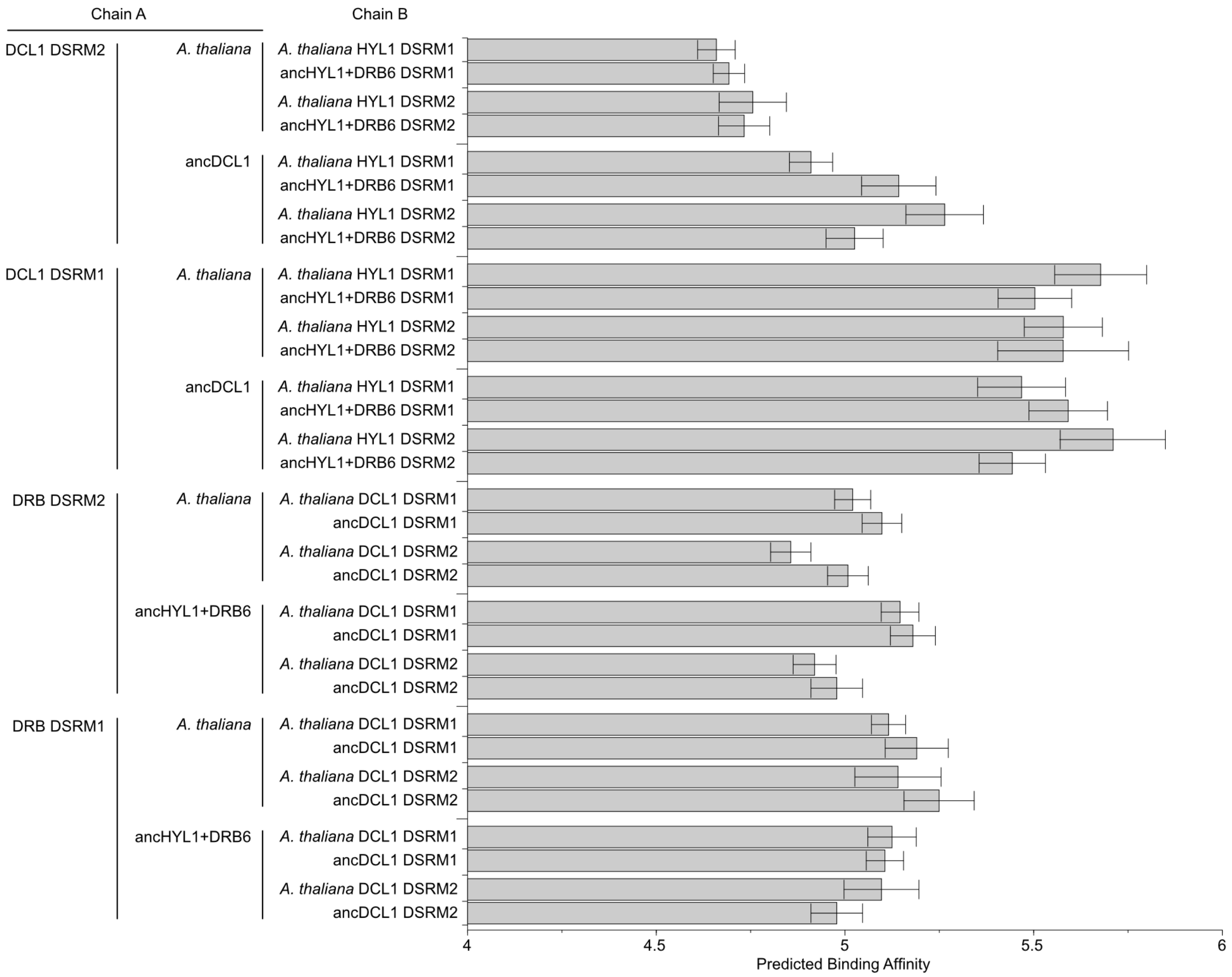


**Figure S7. Structure-based protein-protein affinity analysis predicts plant DCL1's DSRM1 interacts with DSRMs from HYL1.** We modeled the structural interface between ancestral and extant DCL1 DSRMs bound to DSRMs from ancestral HYL1+DRB6 and *Arabidopsis thaliana* HYL1 in two different conformations, based on an X-ray structure of a DSRM-DSRM dimer (PDBID 3ADI). We plot the mean and standard error in predicted binding affinity (pKd = -log(Kd)) of each protein-protein complex, inferred using a statistical machine learning approach, with longer bars indicating tighter binding (see Methods). Inferred complexes are shown in Supplementary Information Figure S8.

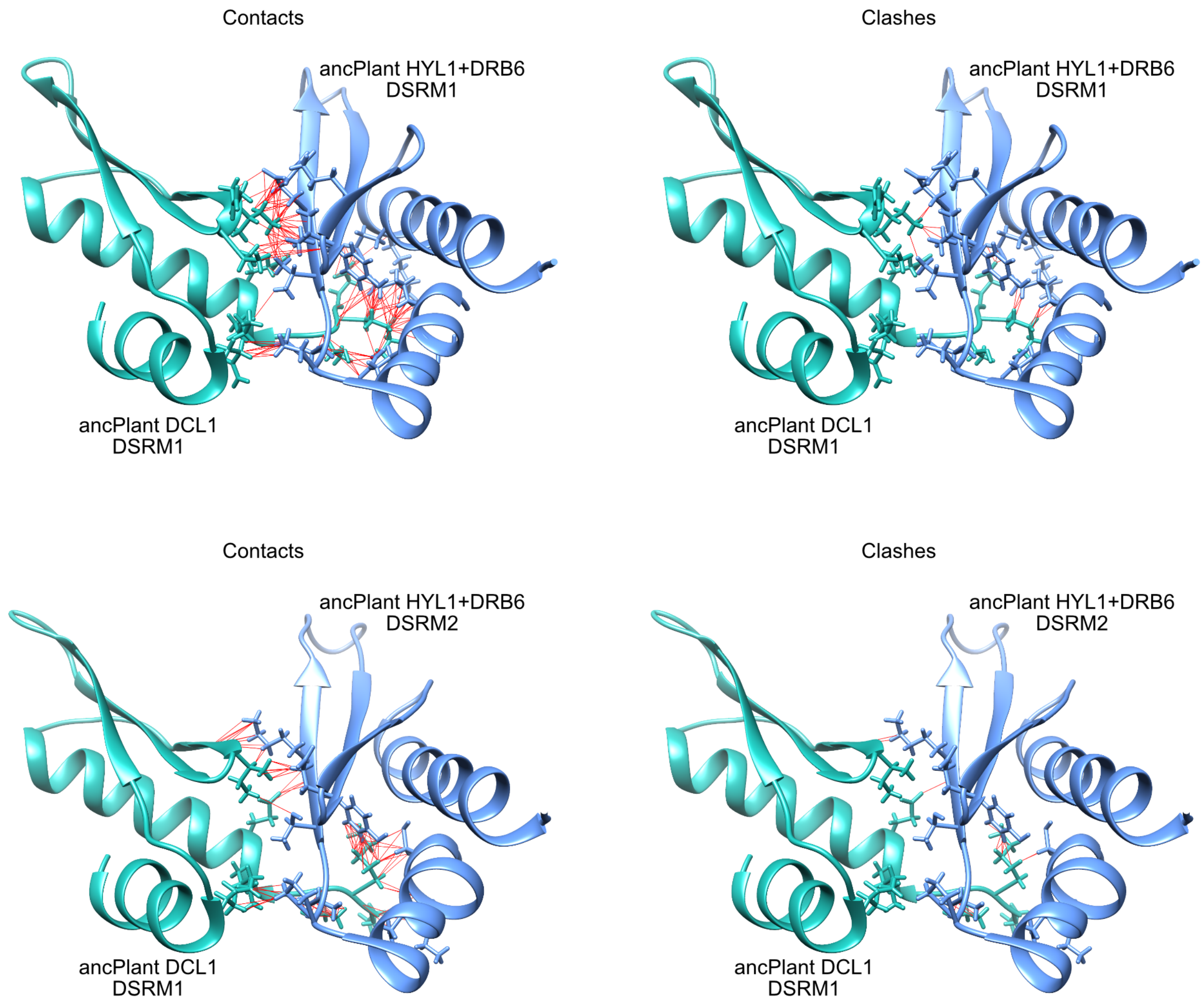


**Figure S8, part 1.**

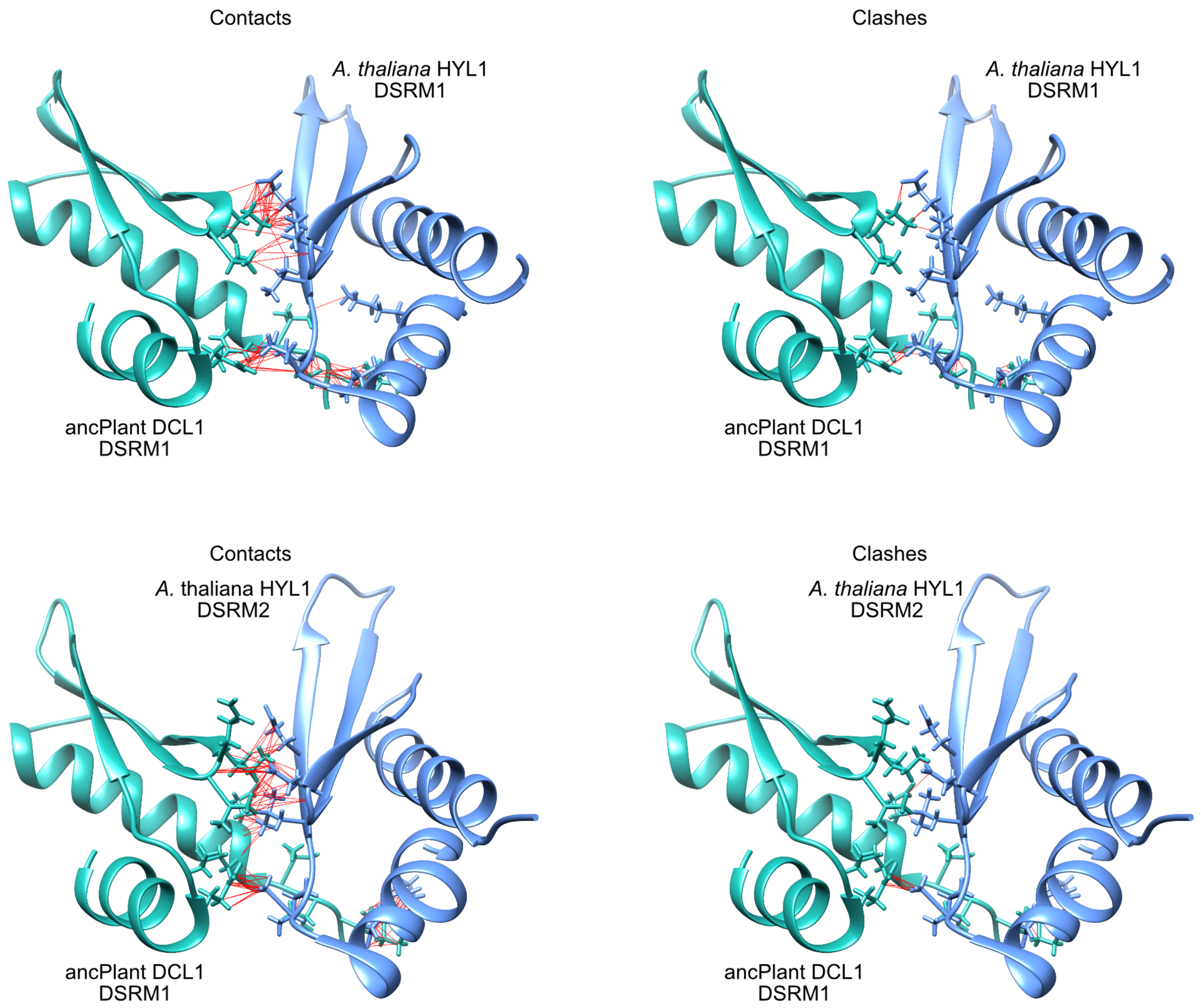


**Figure S8, part 2.**

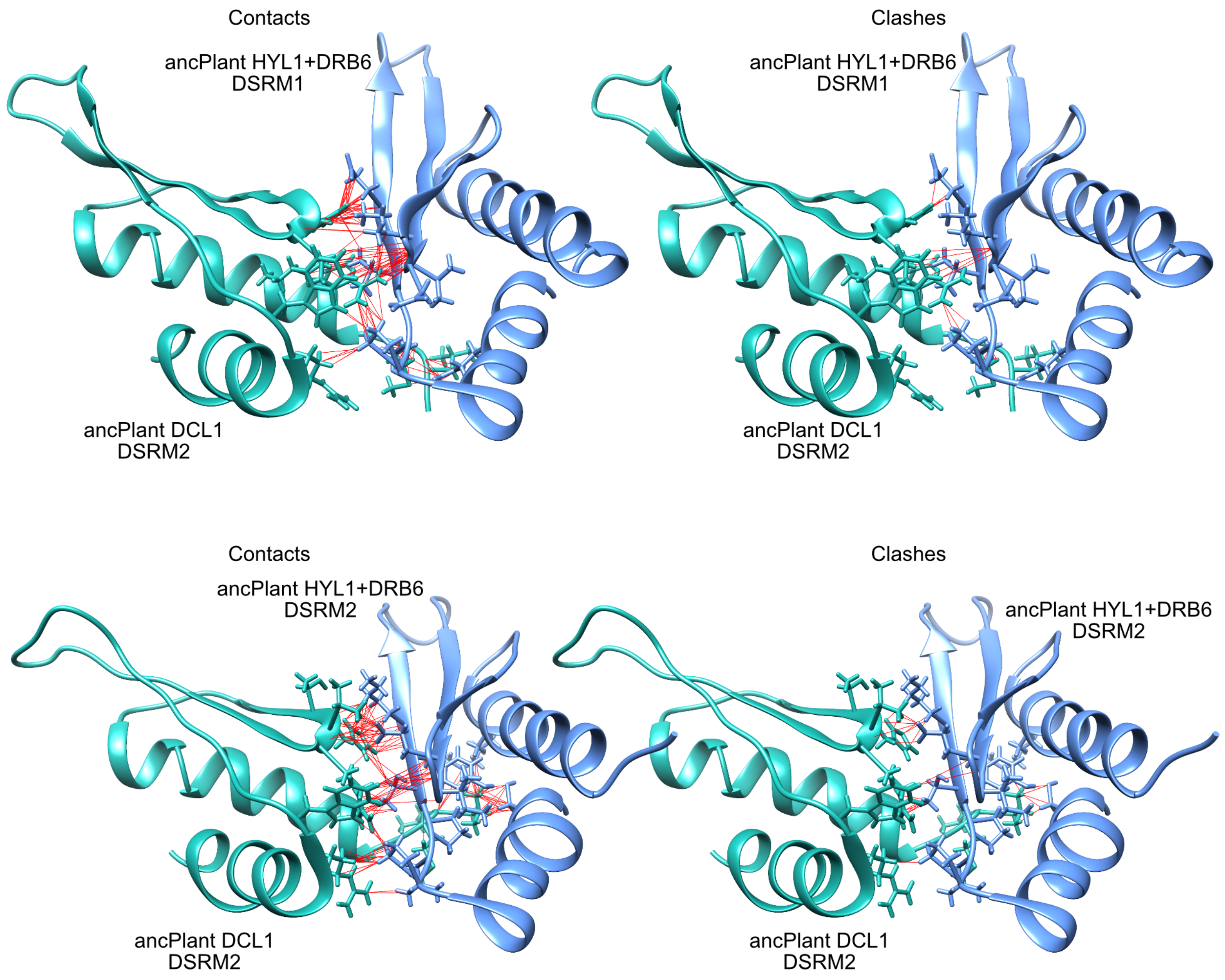


**Figure S8, part 3.**

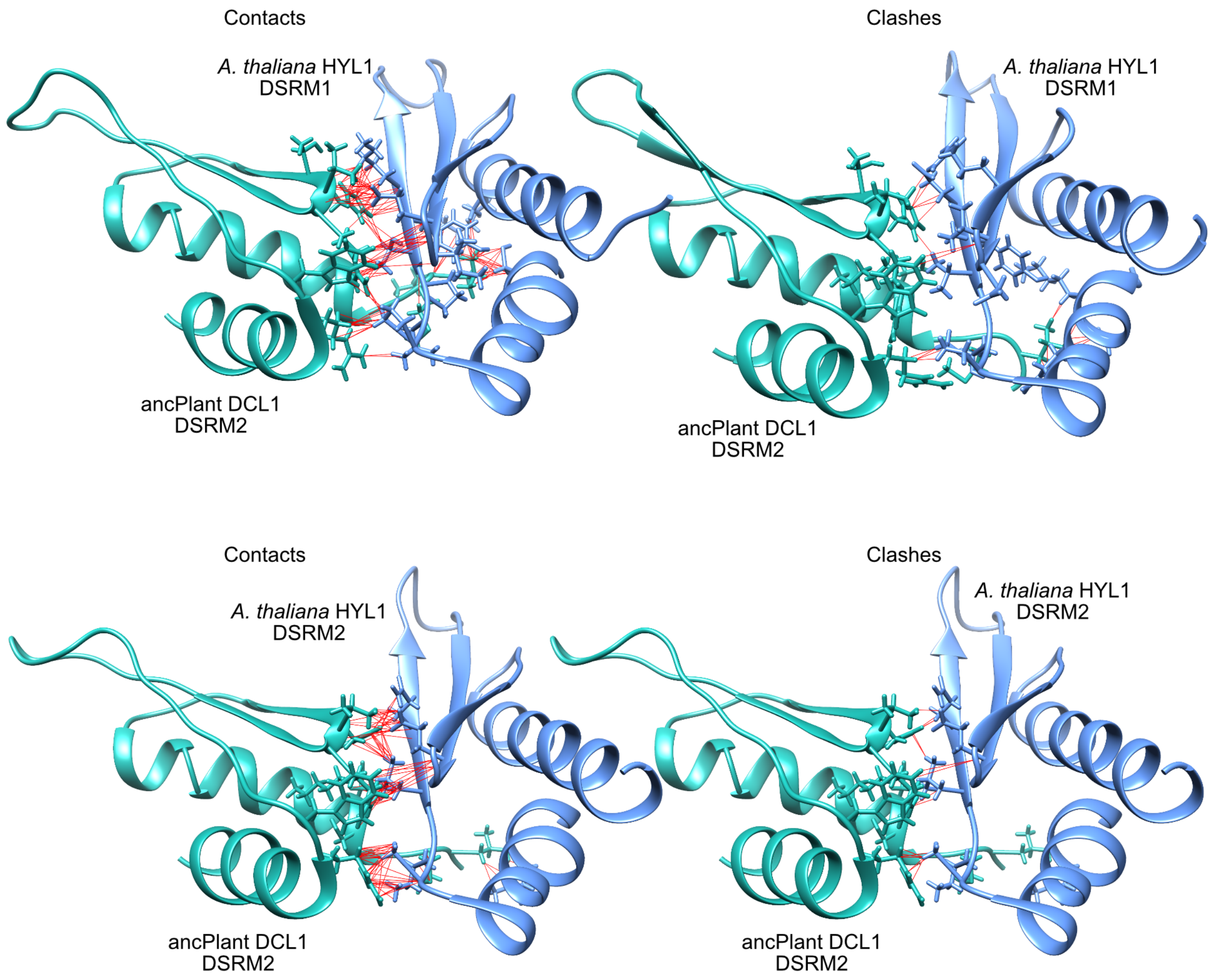


**Figure S8, part 4.**

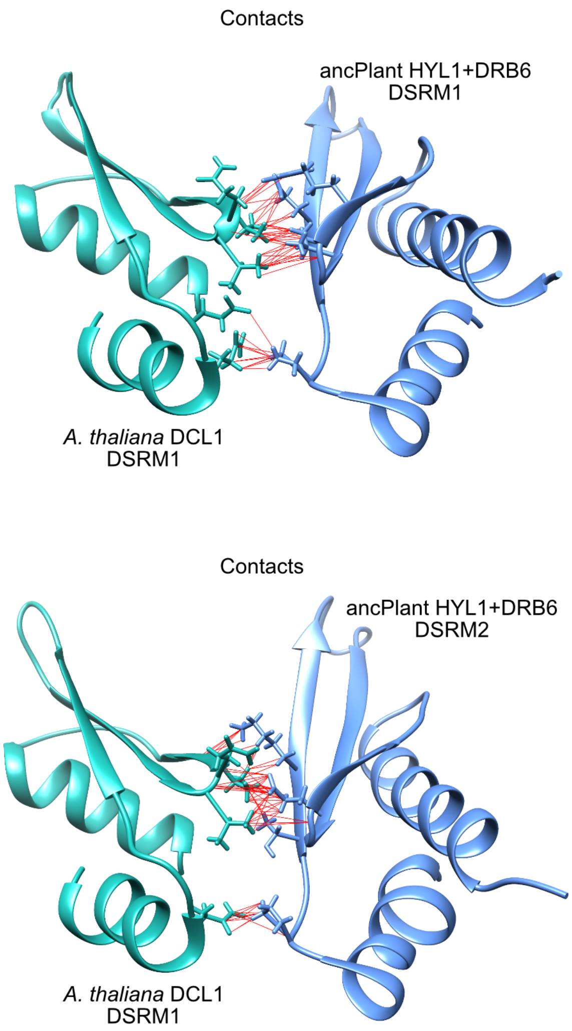


**Figure S8, part 5.**

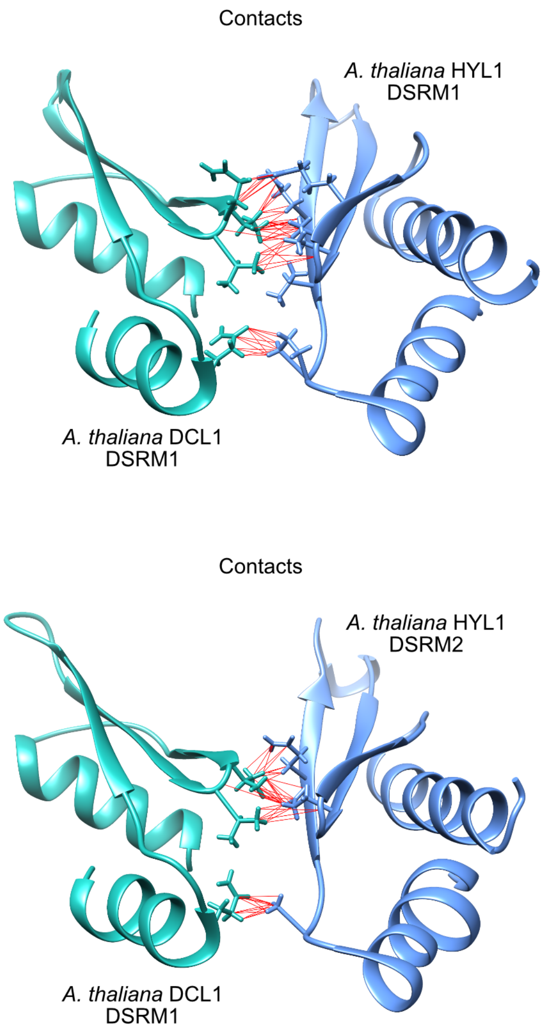


**Figure S8, part 6.**

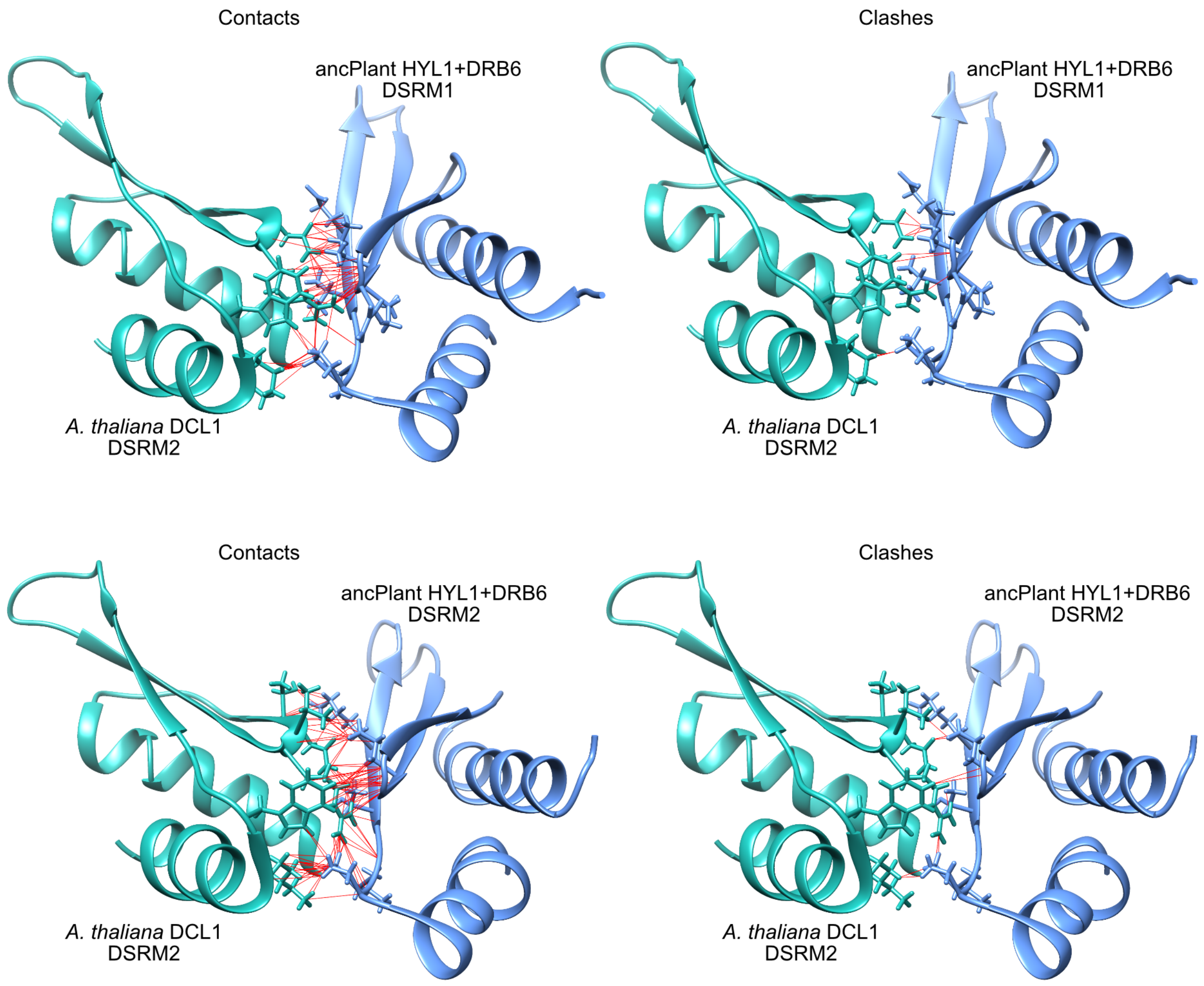


**Figure S8, part 7.**

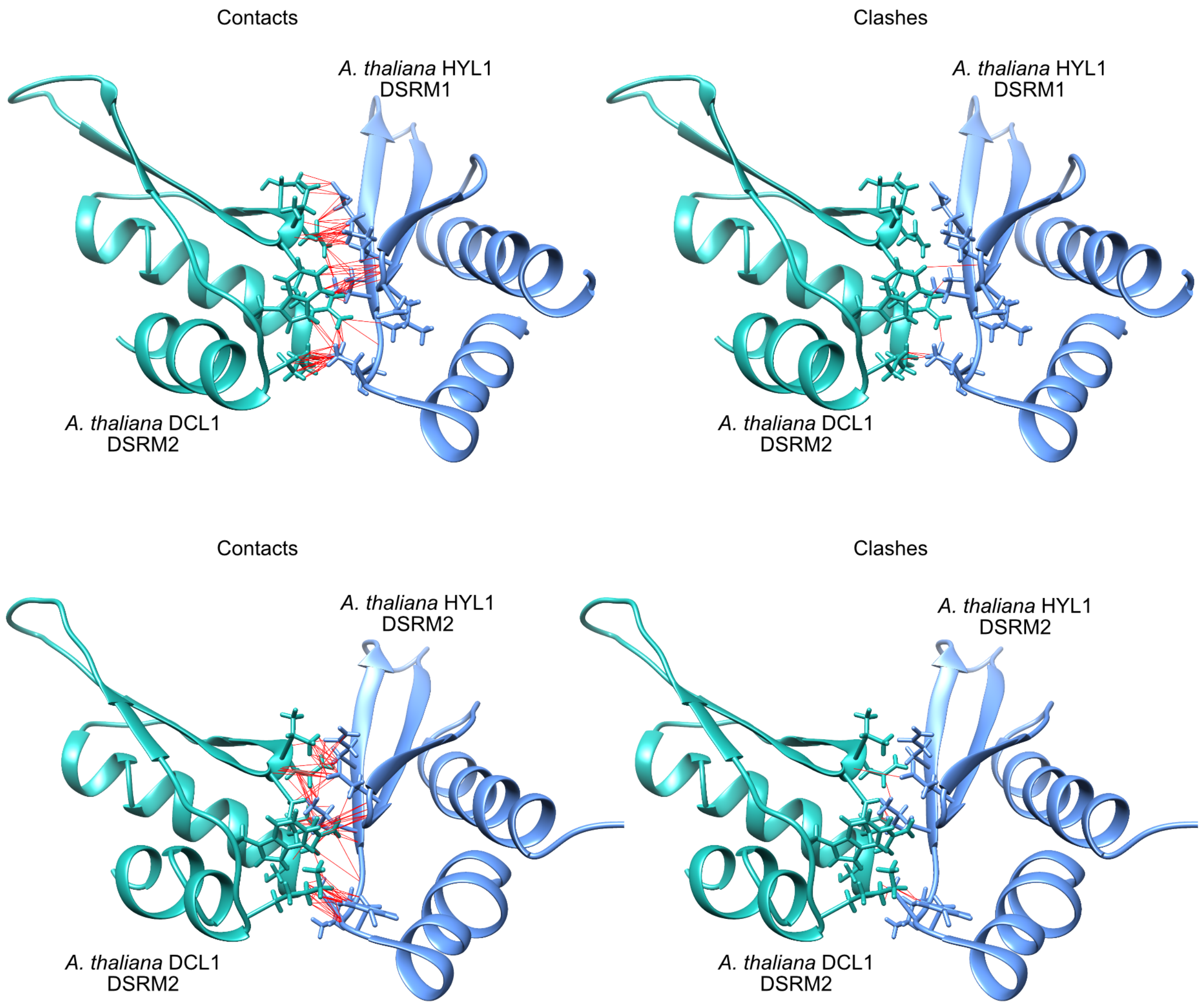


**Figure S8, part 8.**

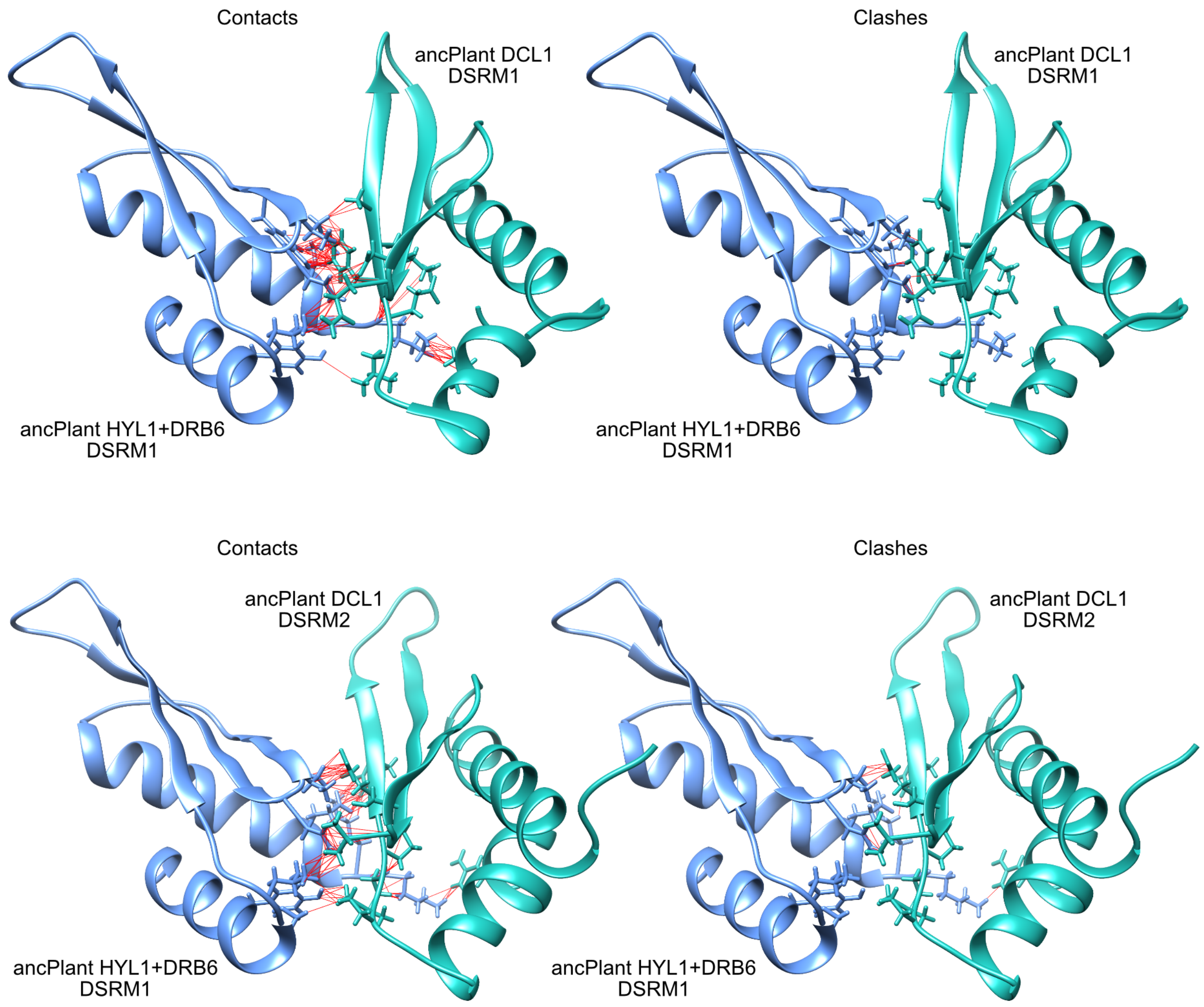


**Figure S8, part 9.**

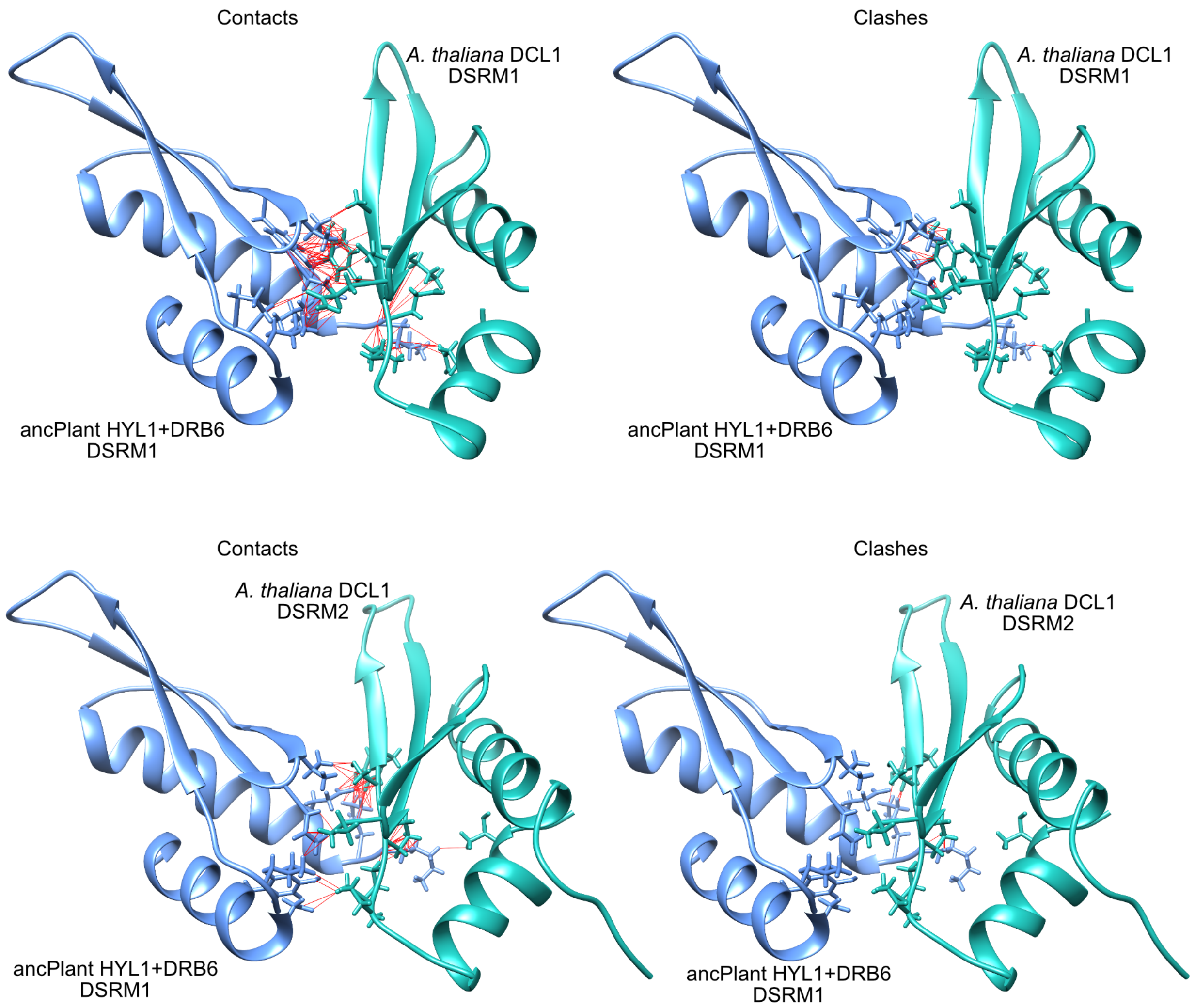


**Figure S8, part 10.**

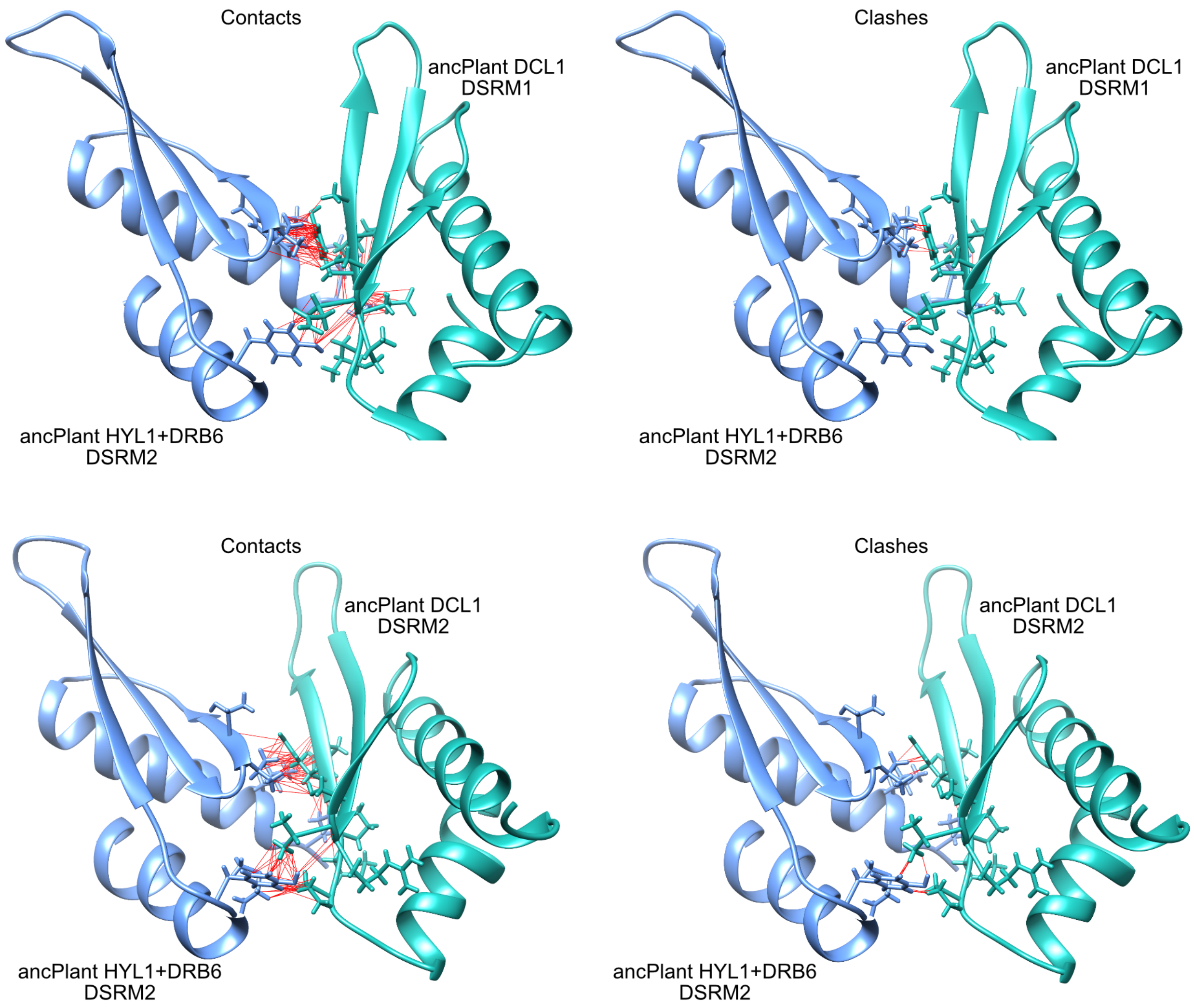


**Figure S8, part 11.**

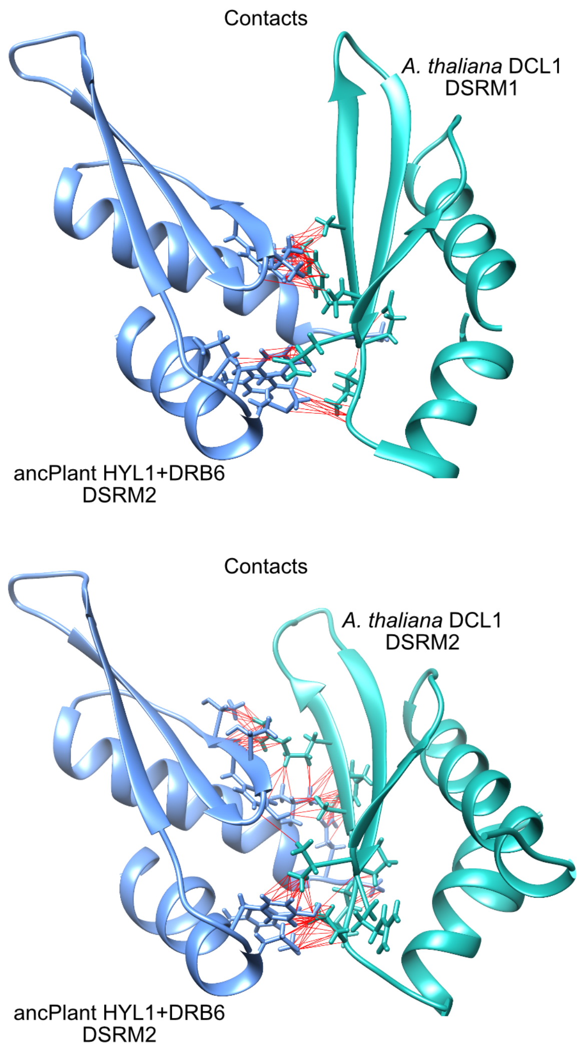


**Figure S8, part 12.**

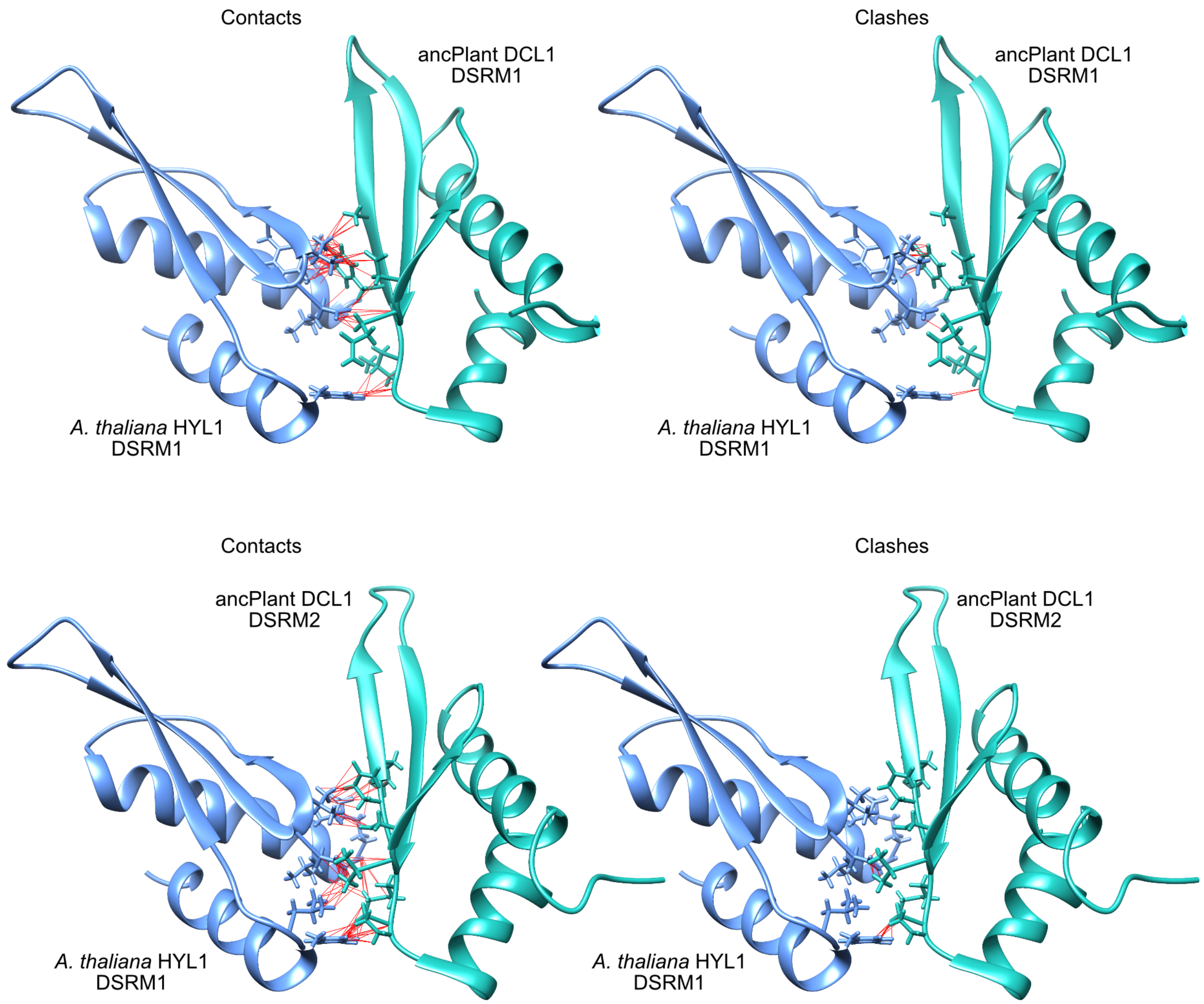


**Figure S8, part 13.**

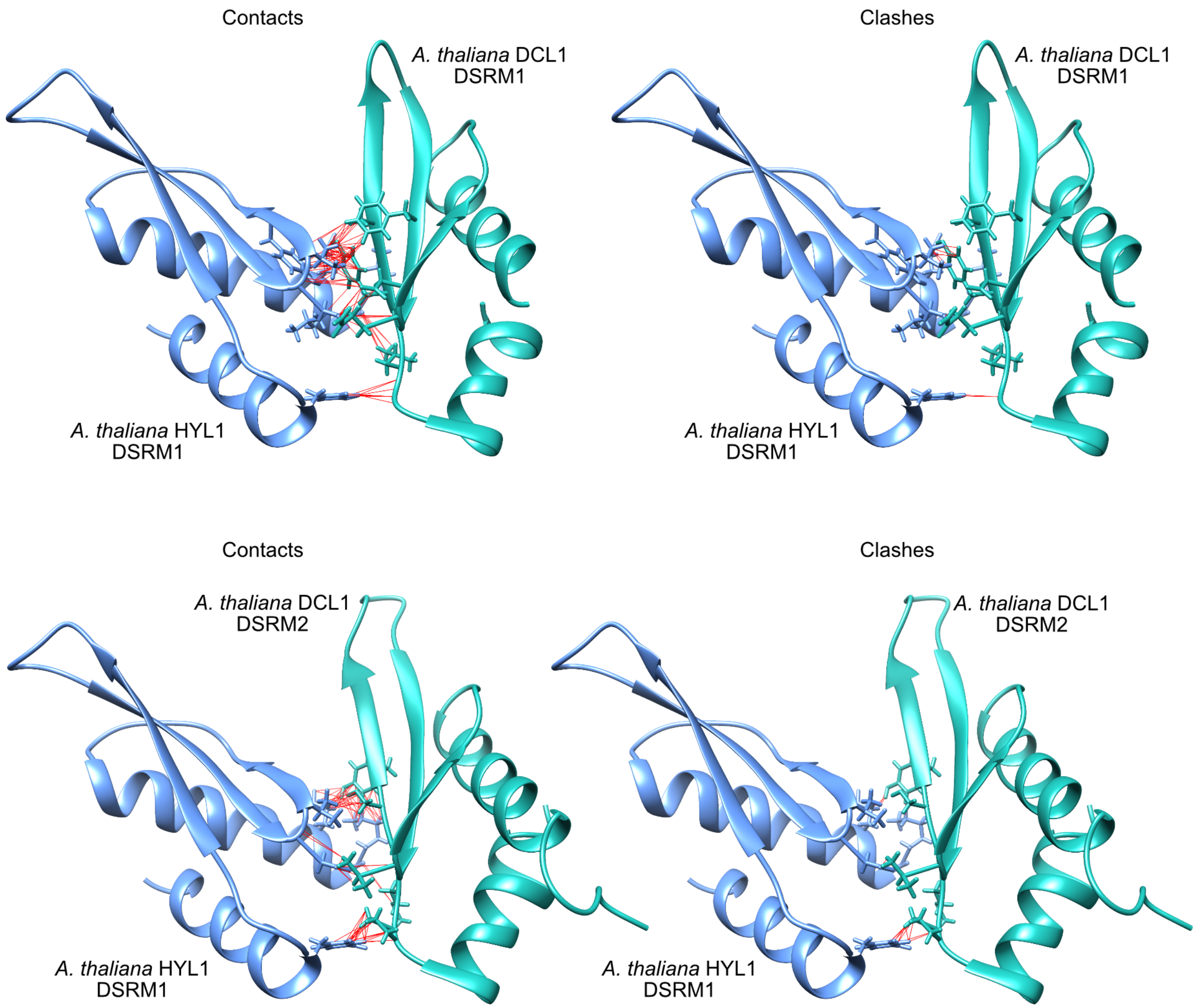


**Figure S8, part 14.**

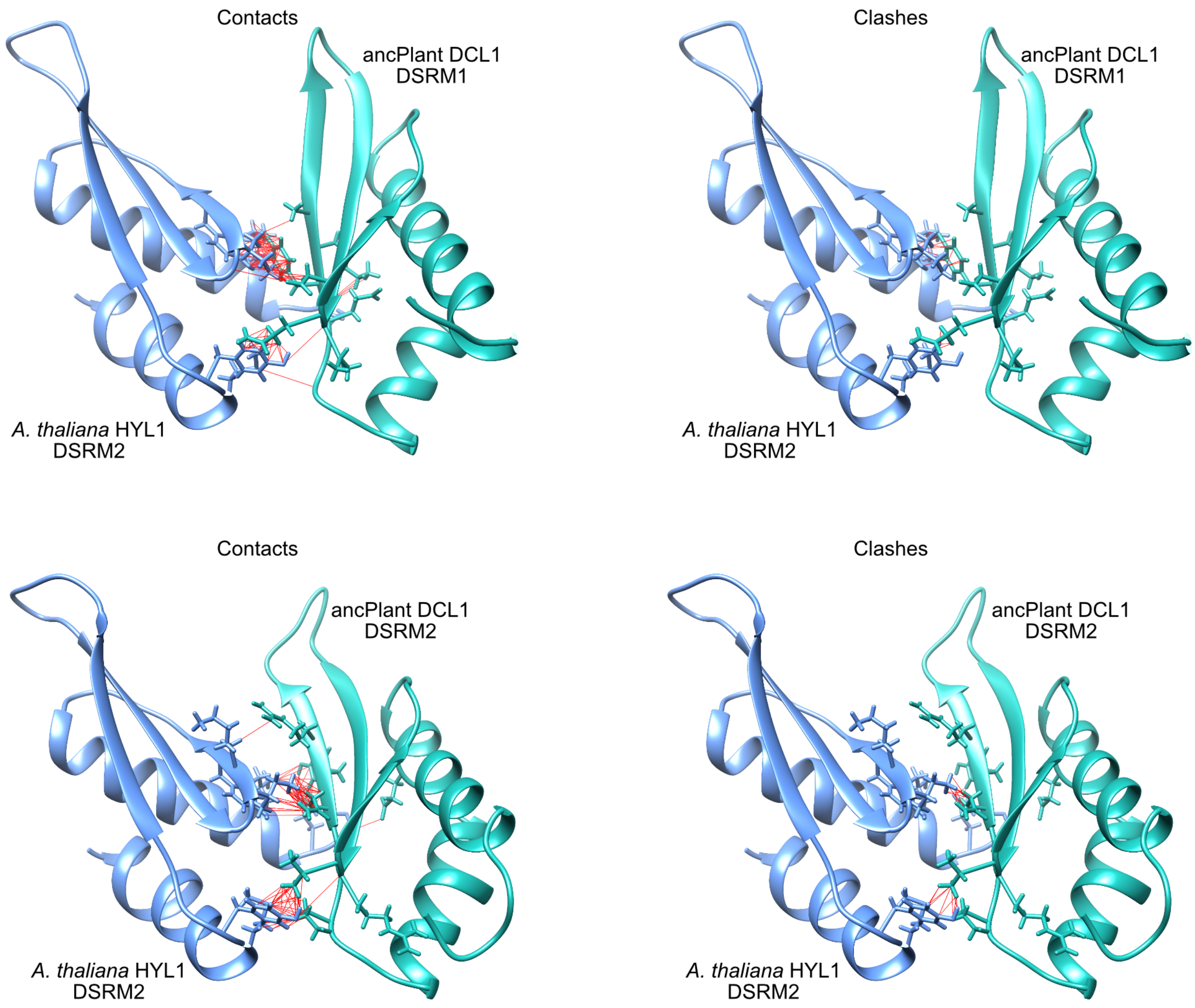


**Figure S8, part 15.**

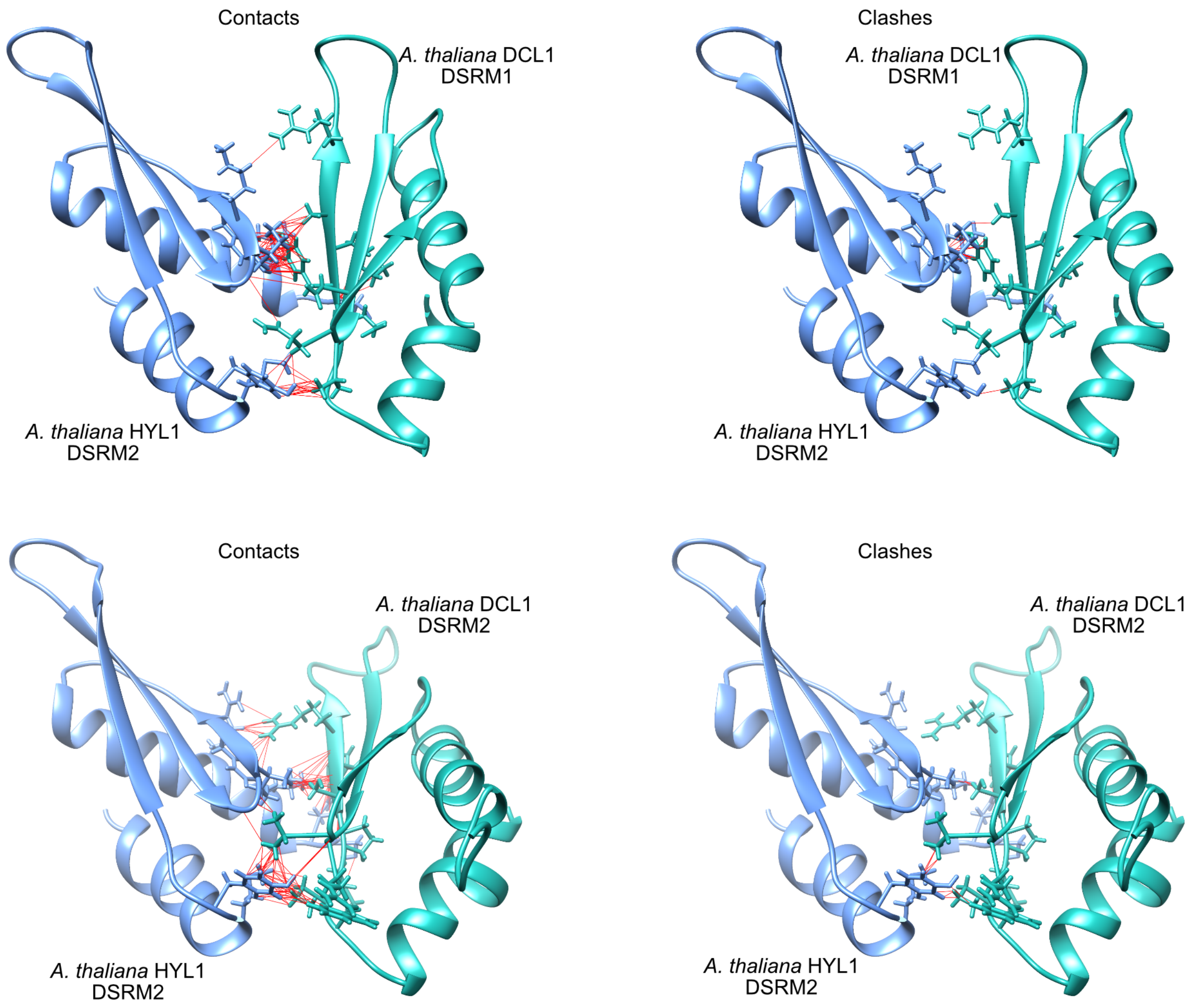


**Figure S8, part 16. Structural models of DCL1 and HYL1 DSRM-DSRM complexes.** We plot the best-scoring modeled complexes of ancestral-reconstructed and *Arabidopsis thaliana* DSRMs from the DCL1 (green) and HYL1 (blue) lineages (see Methods). For visualization, red lines indicate atom-atom contacts (left panel, defined as at least 0.4 angstroms between atomic VDW radii) or atomic clashes (right panel, defined as having >0.6 angstroms overlap between atomic VDW radii). Residues participating in potential intermolecular contacts or clashes are shown as stick models.

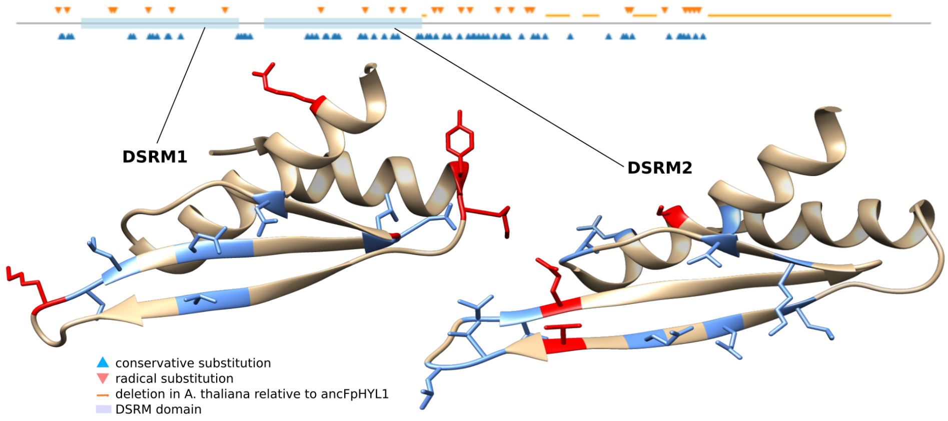


**Figure S9. Ancestral flowering plant HYL1 (ancFpHYL1) is very different from *Arabidopsis thaliana* HYL1.** We plot the amino-acid differences between *A. thaliana* HYL1 and the reconstructed ancFpHYL1 along the pairwise alignment (top panel). Blue upward-pointing arrows indicate conservative substitutions within the amino-acid classes GAVLI, SCTM, FYW, HKR and DEN; red downward-pointing arrows indicate radical substitutions between classes. Orange horizontal lines indicate deletions in *A. thaliana* HYL1, relative to ancFpHYL1. Blue transparent boxes indicate annotated DSRM domains. Structures of *A. thaliana* DSRM1 (PDB ID 3ADG) and DSRM2 3ADJ) are shown in ribbon format, with conservative (blue) and radical (red) amino-acid substitutions indicated.

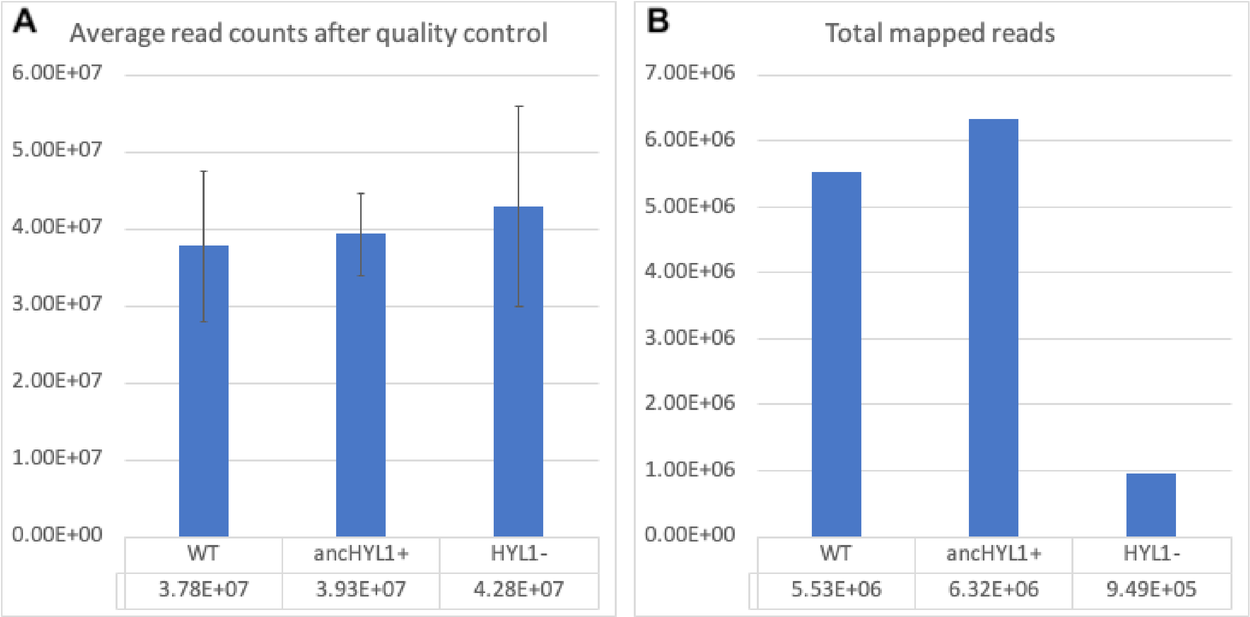


**Figure S10. Total small-RNA sequencing read counts are similar among wild-type (WT), HYL1$^{-}$ knockout and ancFpHYL1-expressiong (ancHYL1+) plants; reads mapping to annotated *A. thaliana* miRNAs is reduced in the HYL1$^{-}$ knockout. A.** We plot the mean and standard error over three replicates of total small-RNA sequencing reads obtained from each plant genotpe after quality filtering. **B.** We plot the total numbers of reads mapping to annotated miRNAs in the Araport 11 *A. thaliana* genome annotation.

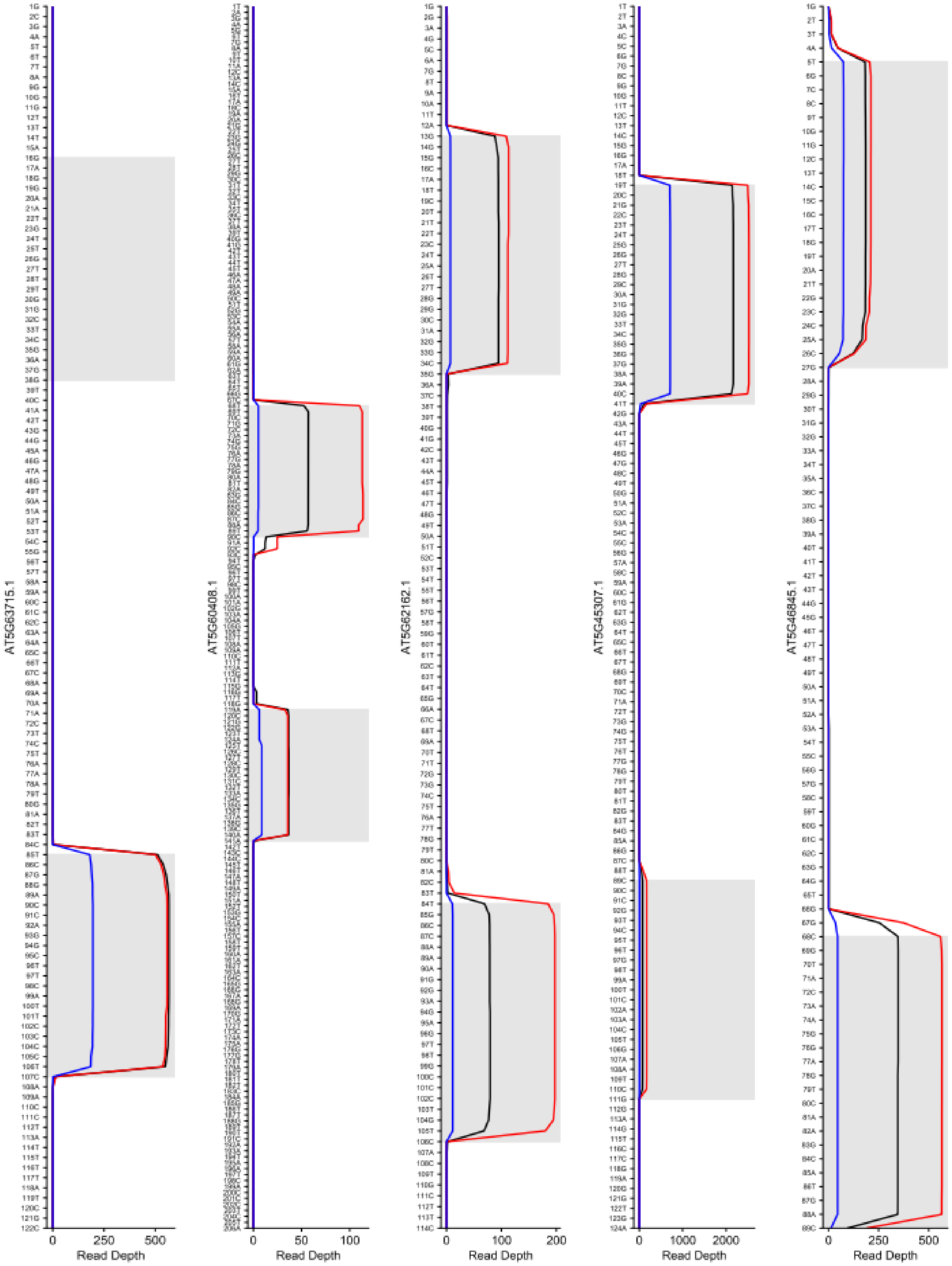


**Figure S11, part 1.**

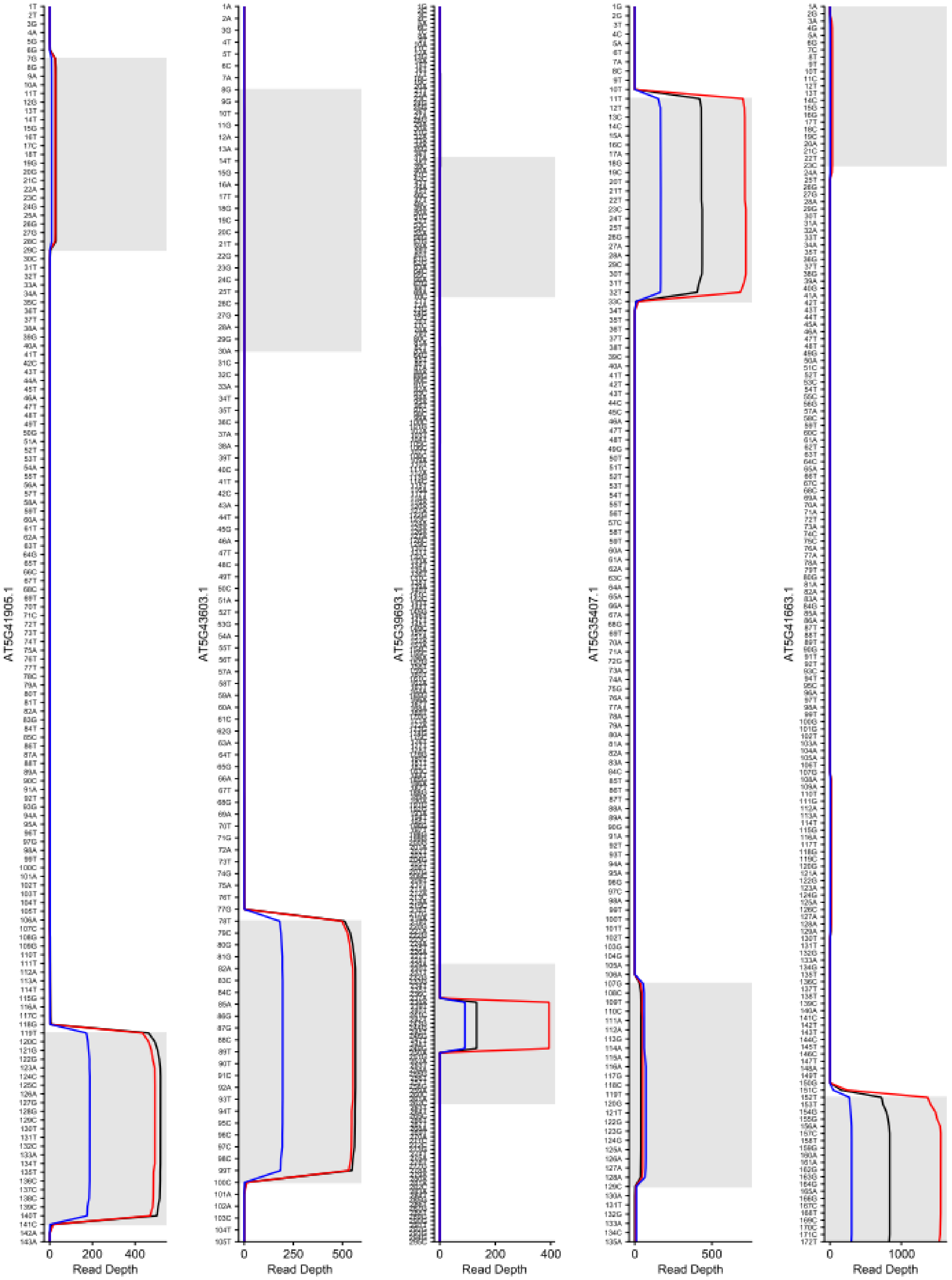


**Figure S11, part 2.**

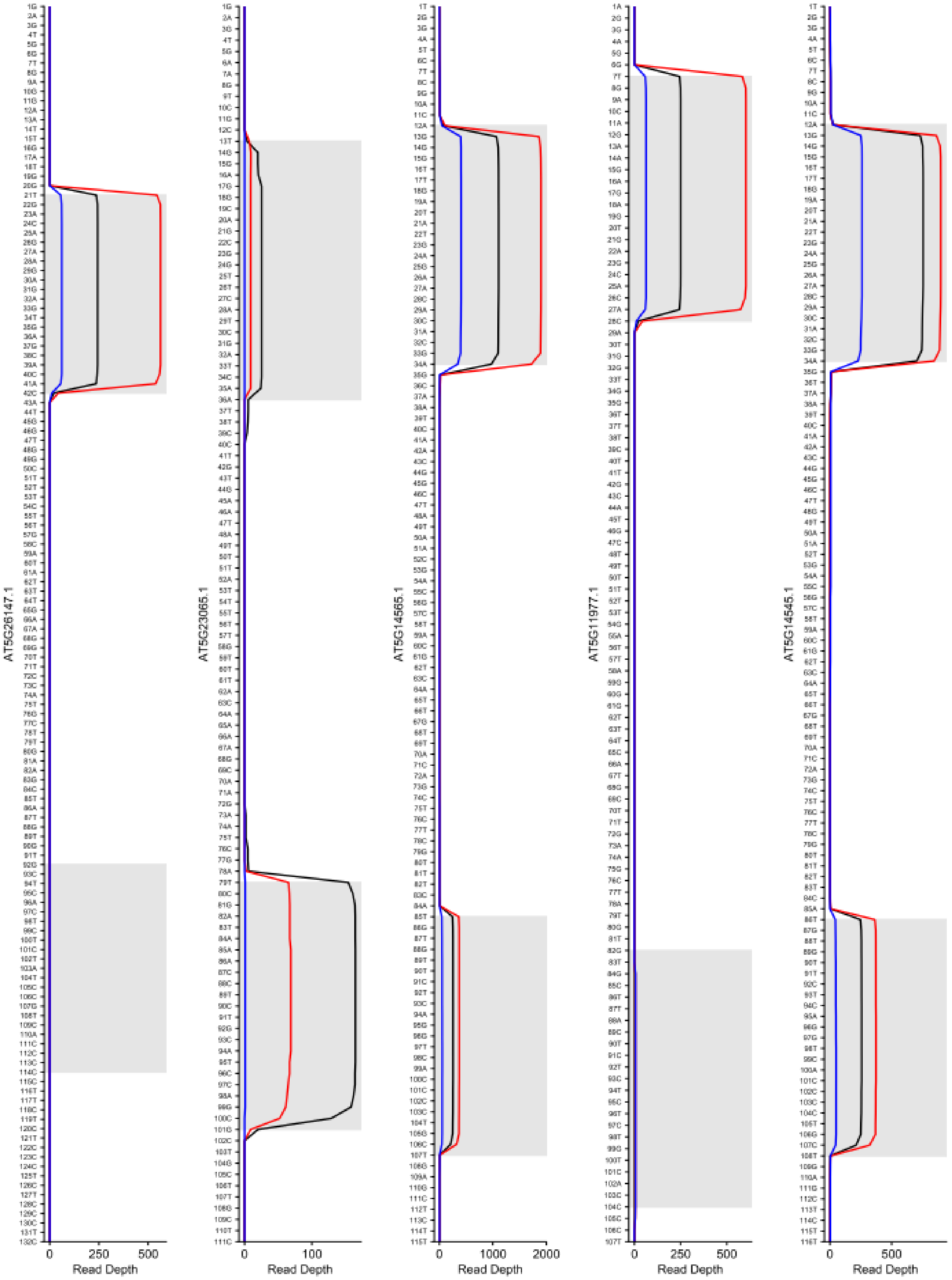


**Figure S11, part 3.**

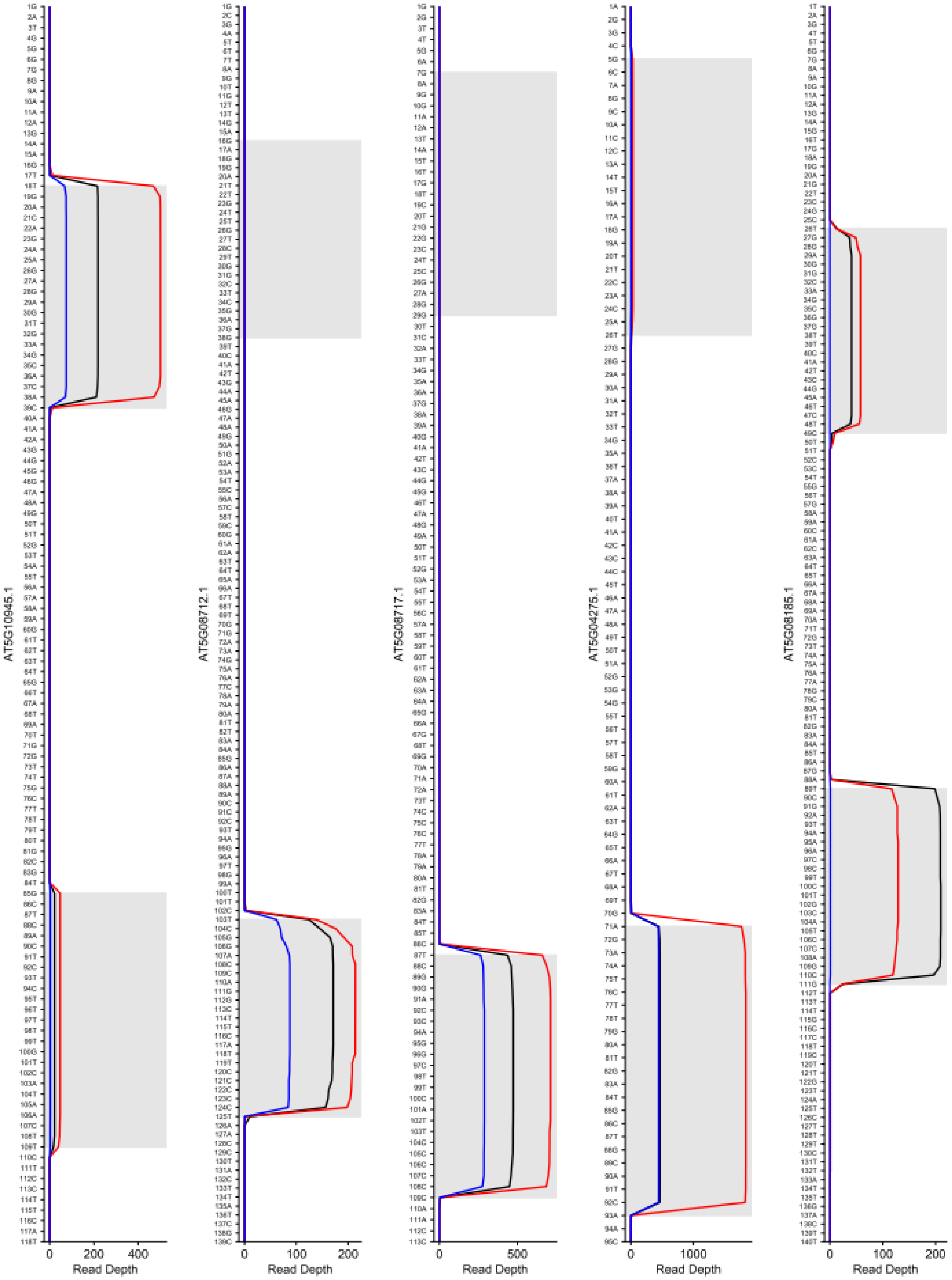


**Figure S11, part 4.**

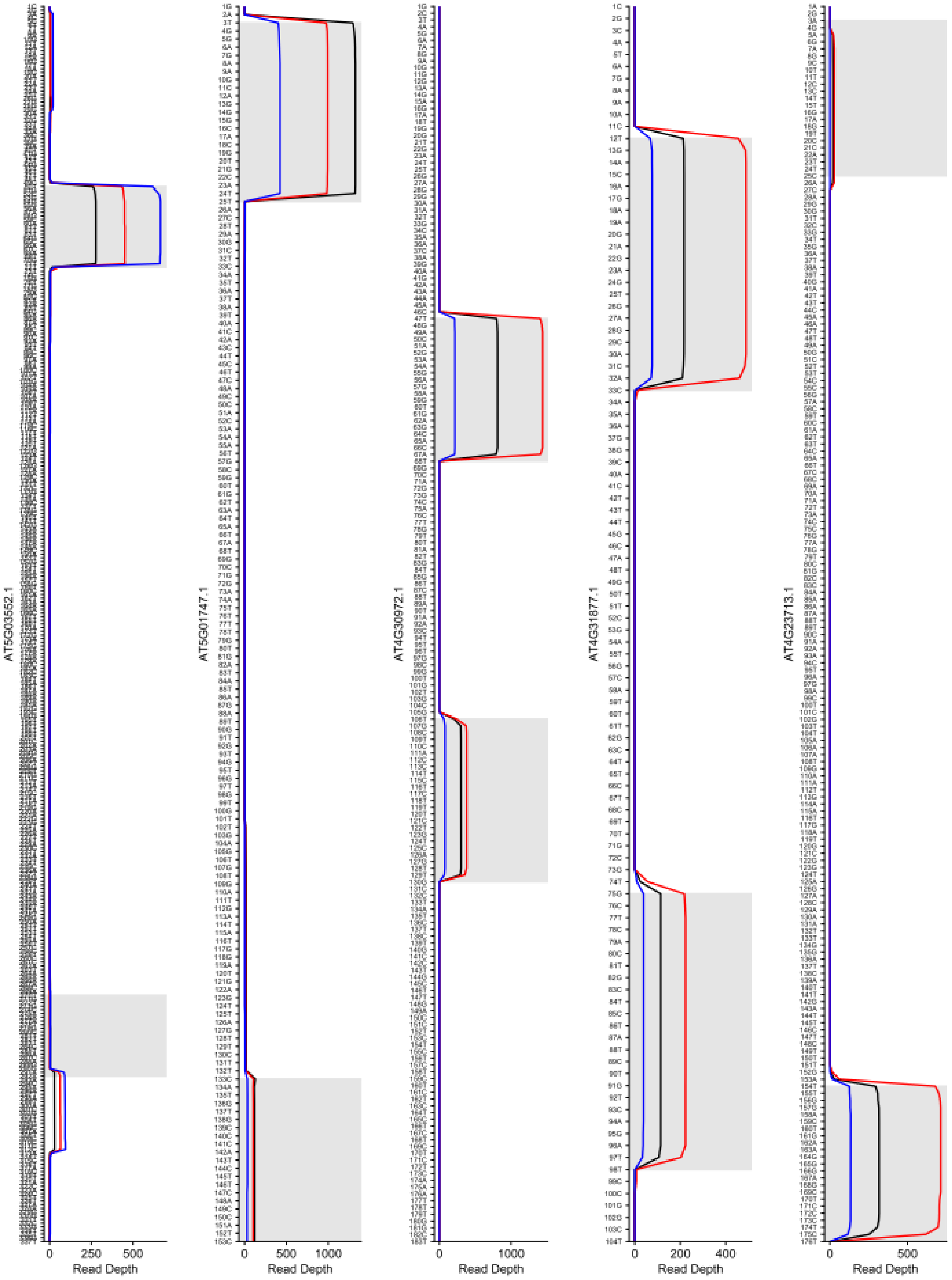


**Figure S11, part 5.**

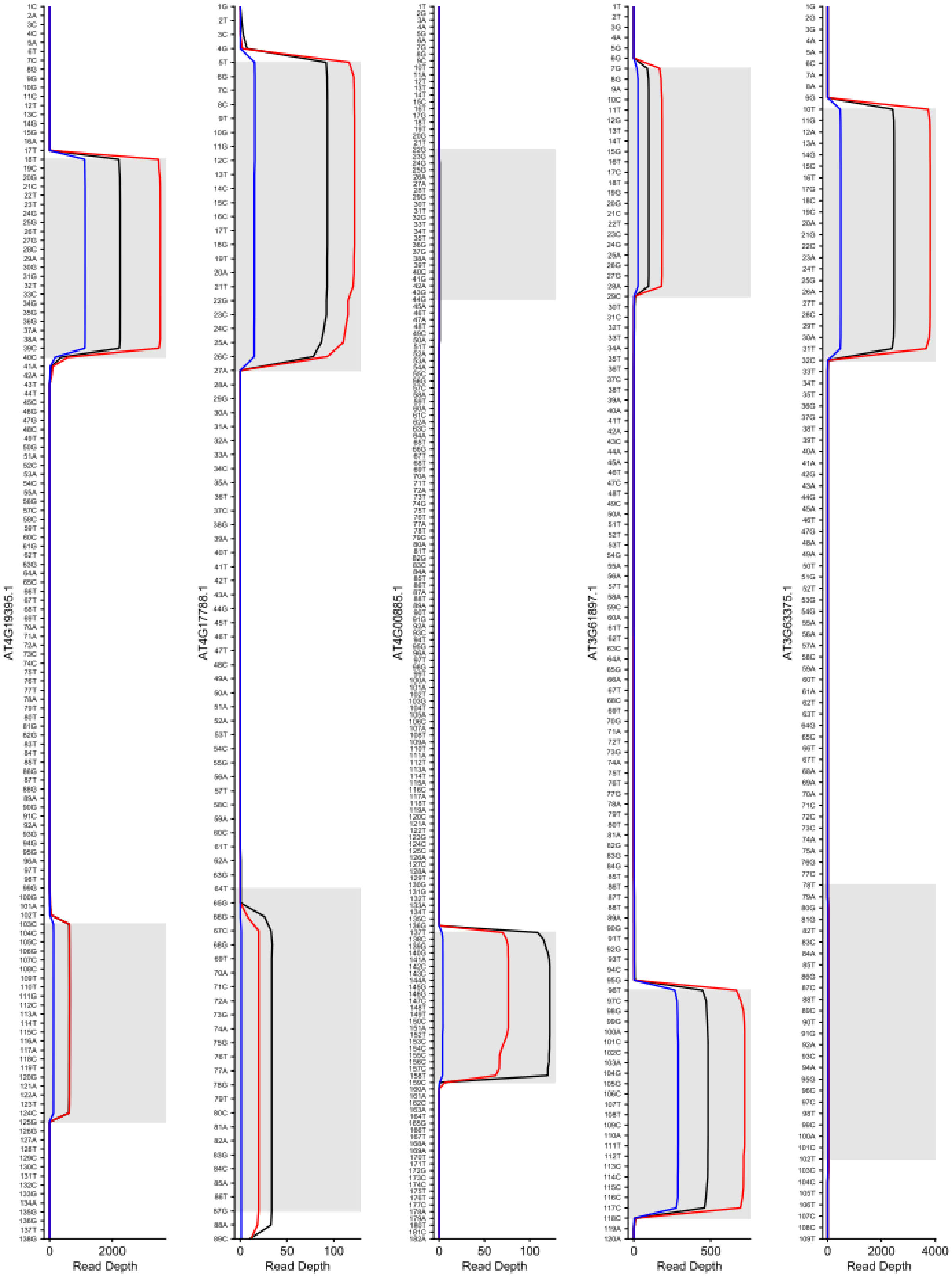


**Figure S11, part 6.**

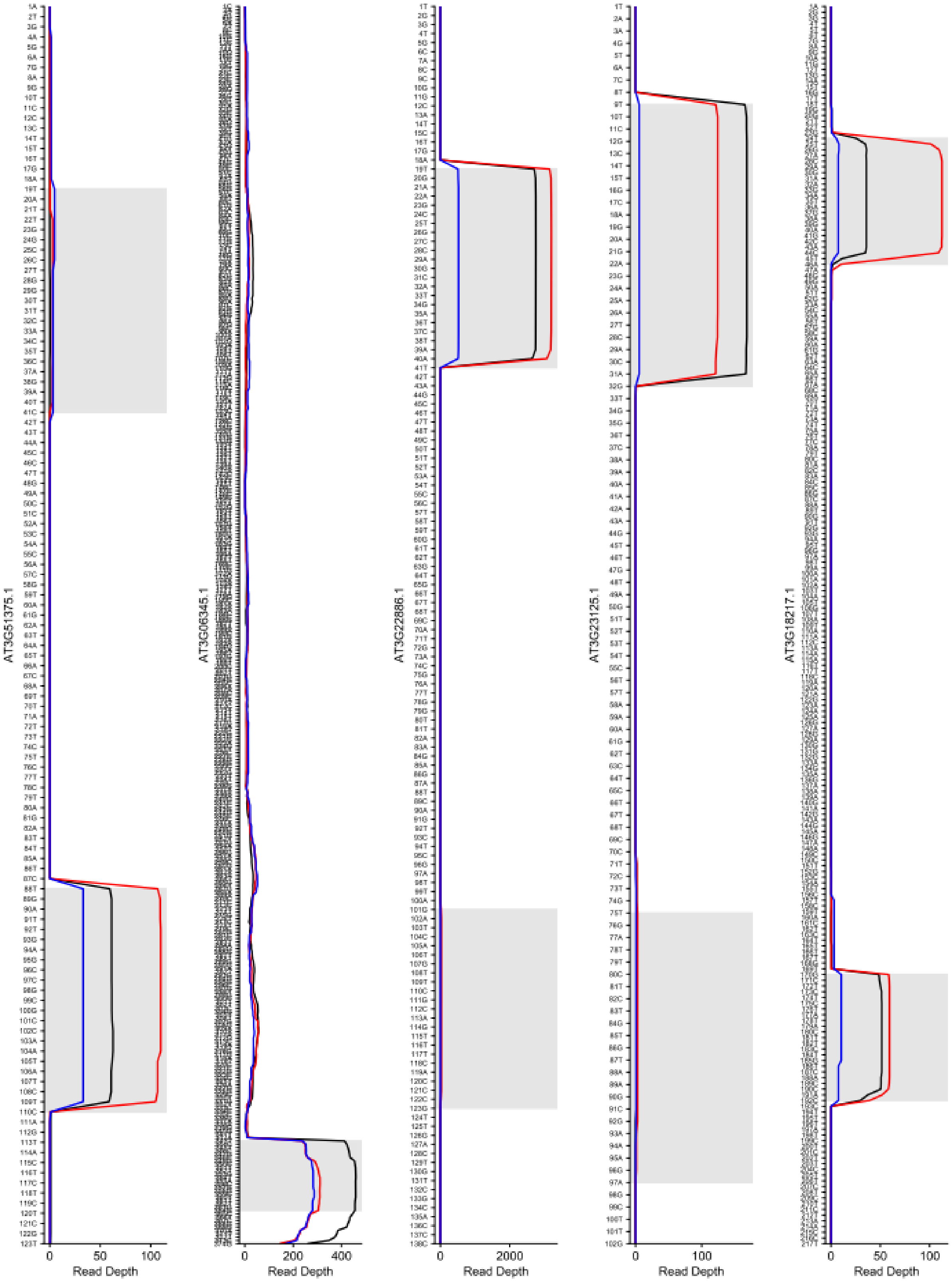


**Figure S11, part 7.**

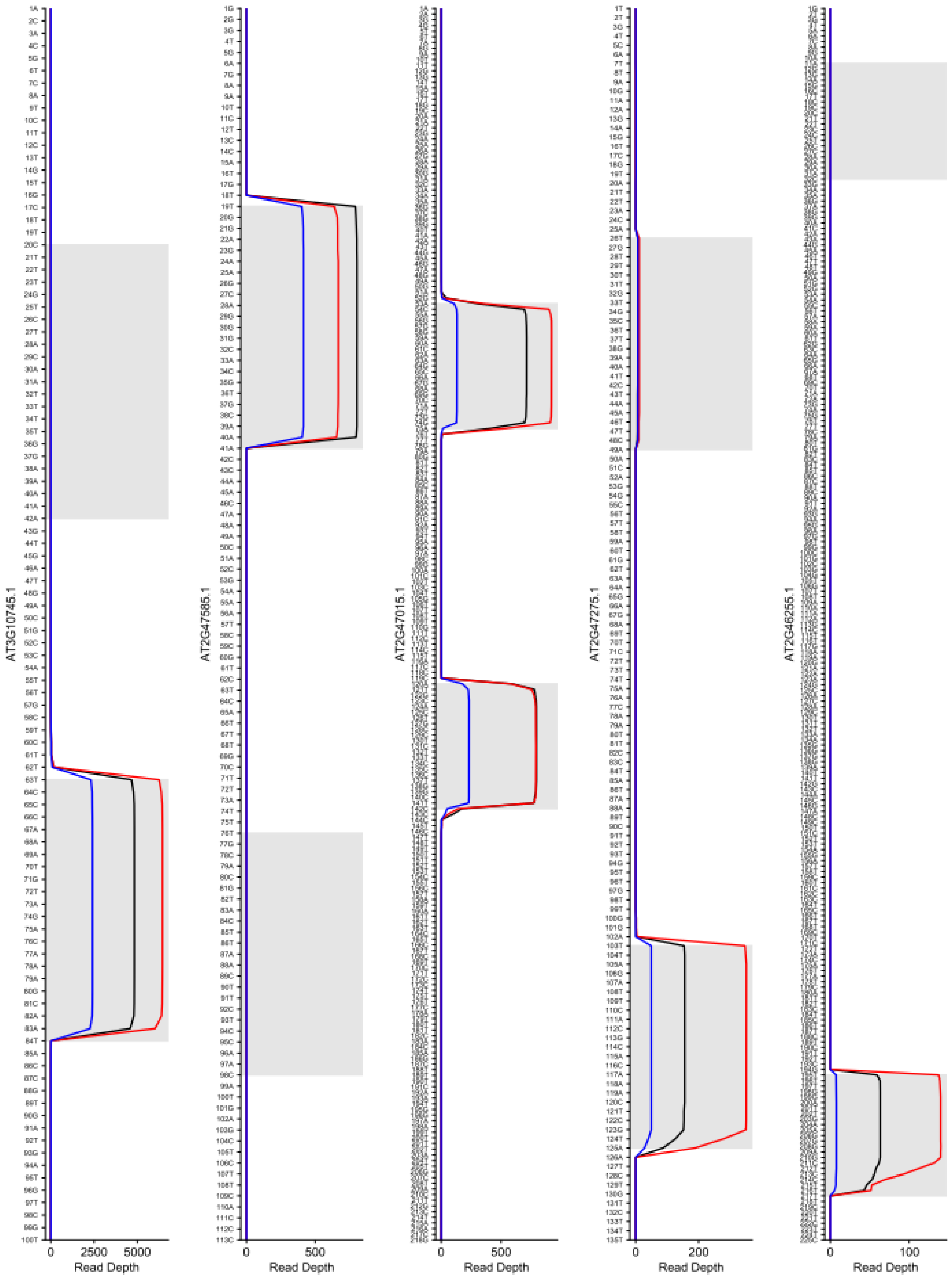


**Figure S11, part 8.**

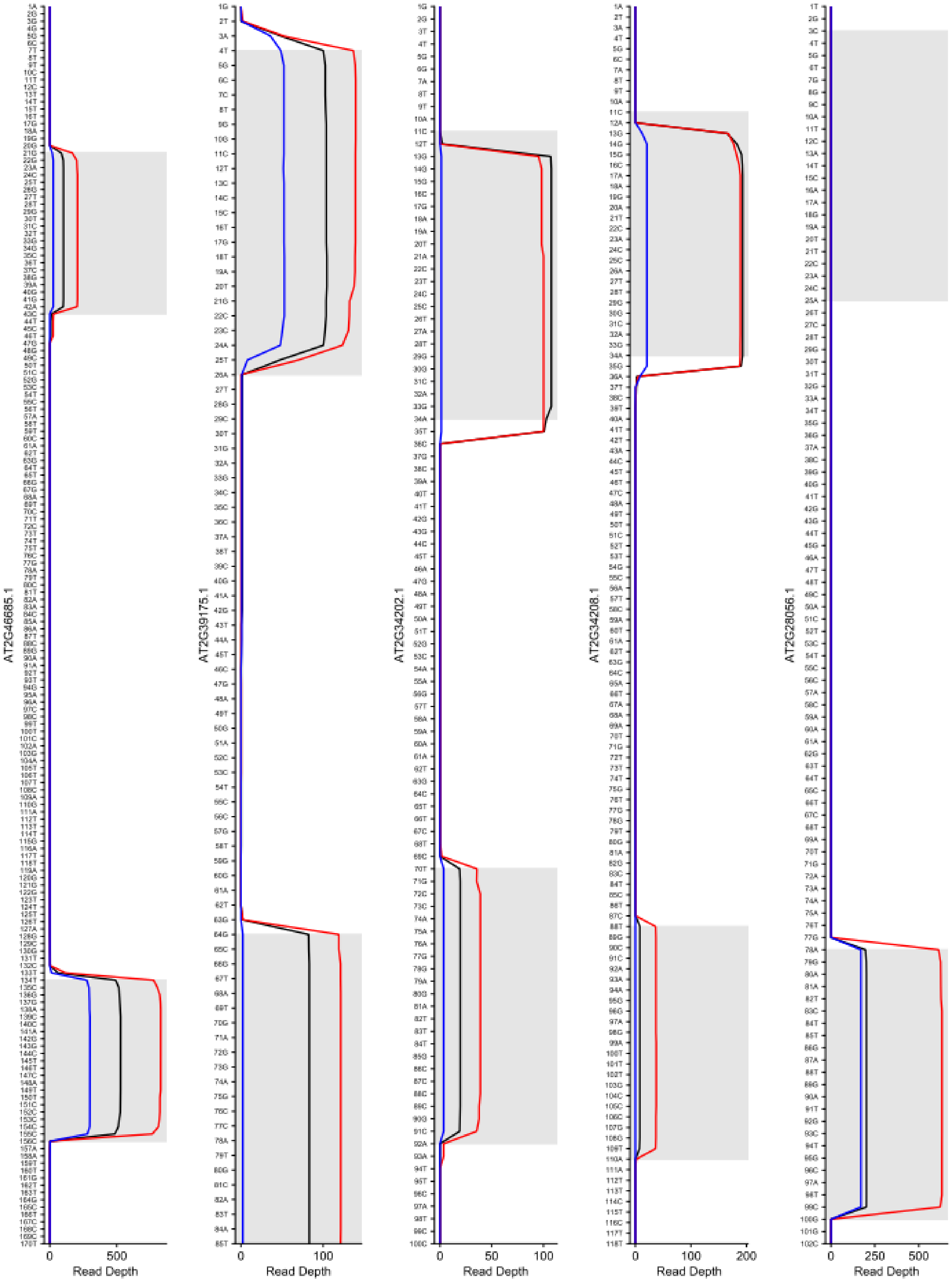


**Figure S11, part 9.**

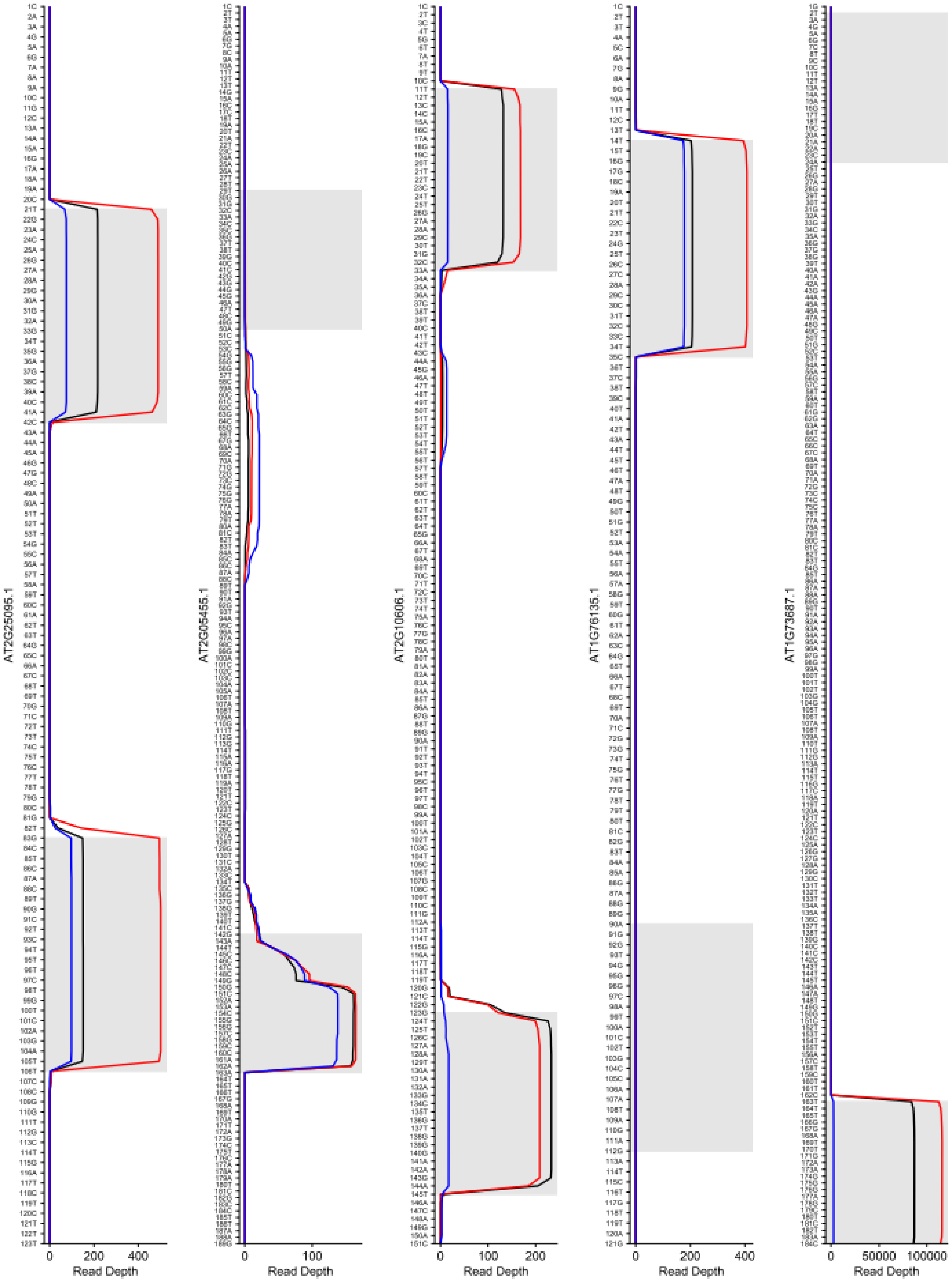


**Figure S11, part 10.**

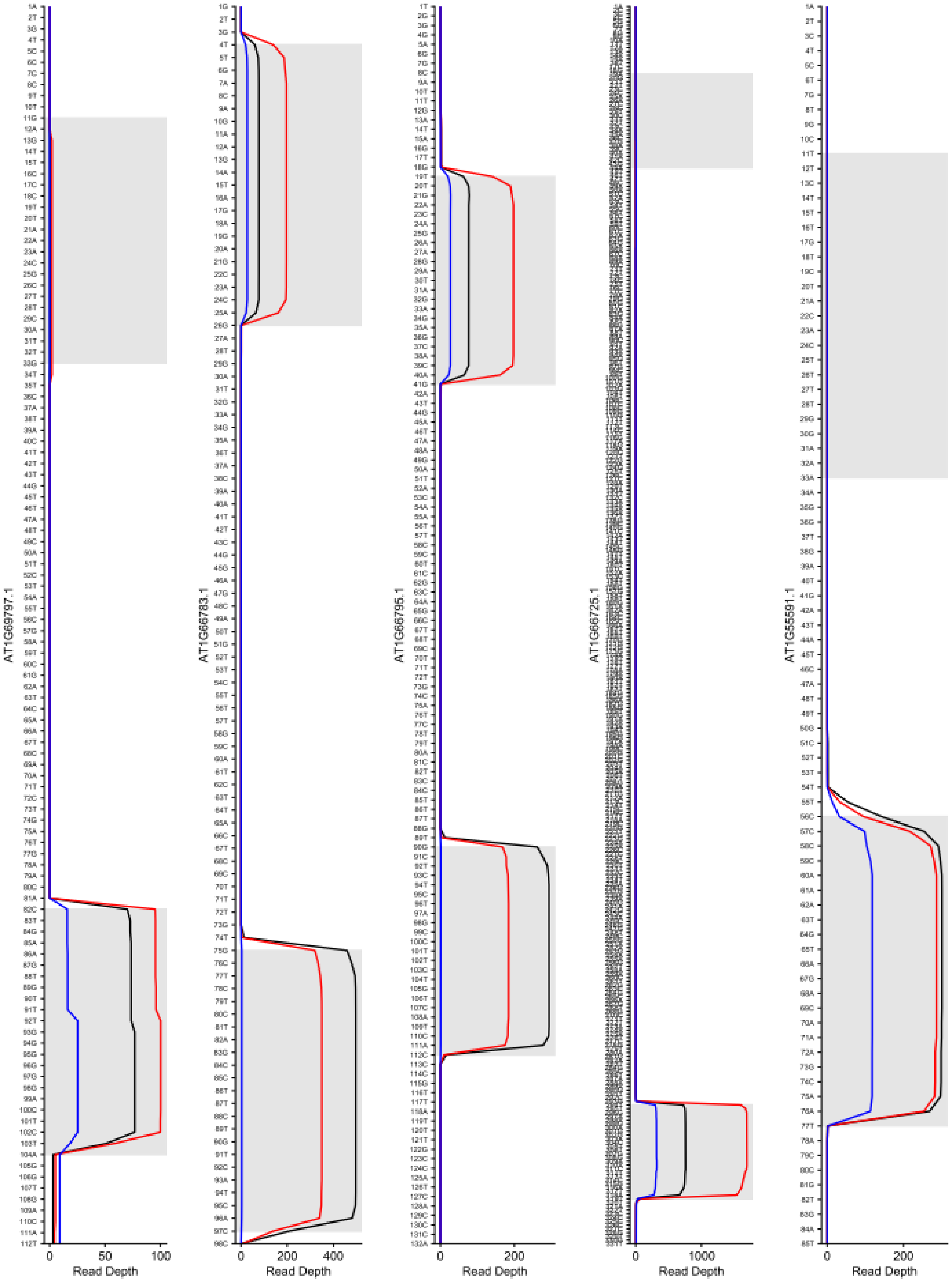


**Figure S11, part 12.**

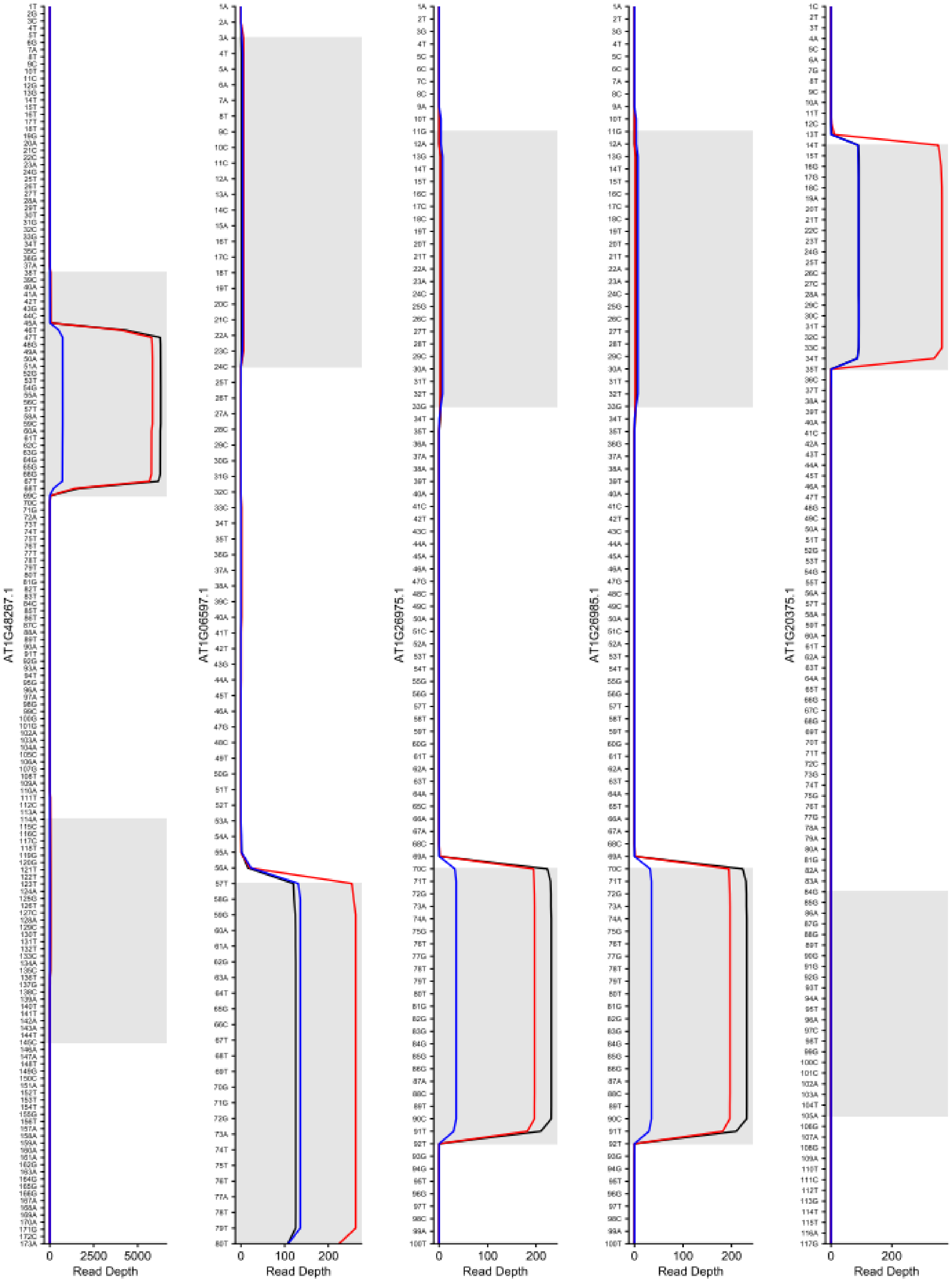


**Figure S11, part 13.**

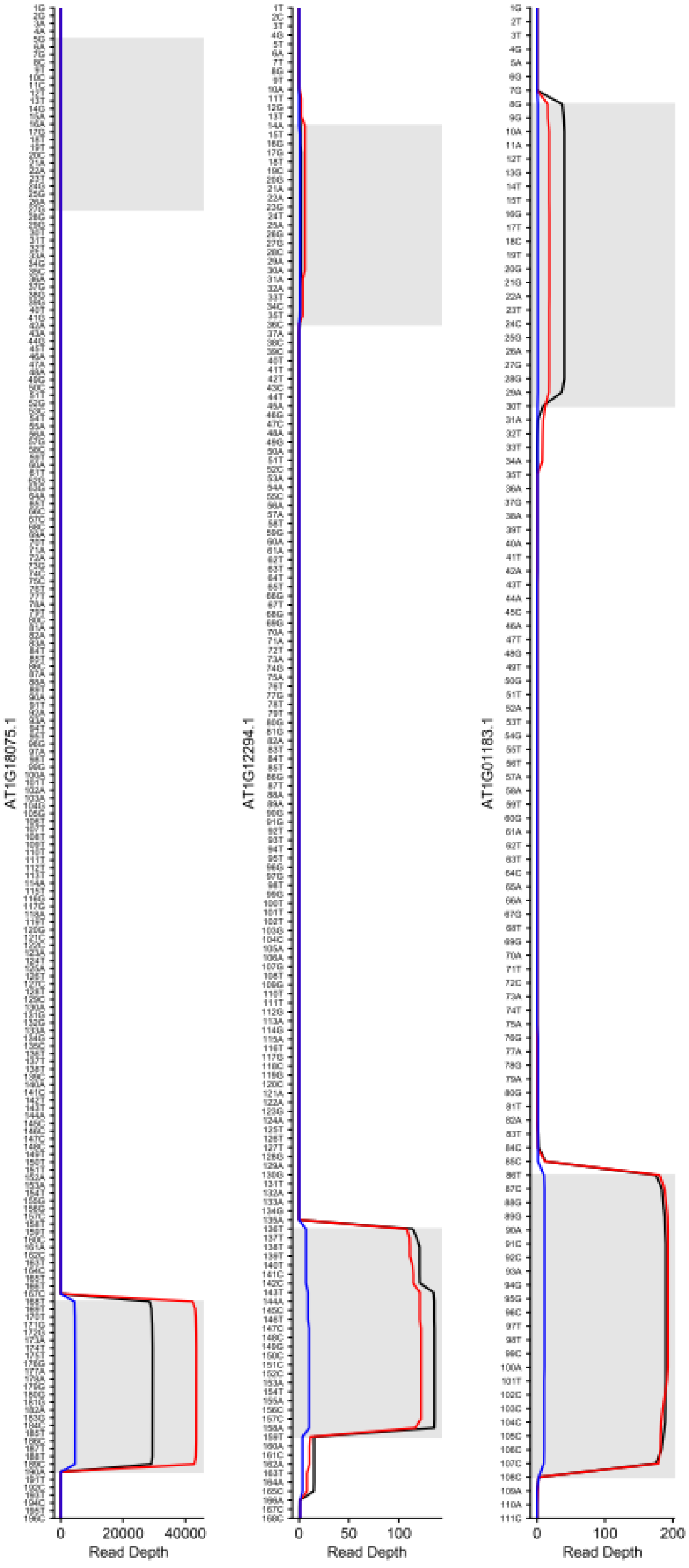


**Figure S11, part 14. Read depth of mapped reads across primary micro-RNA transcripts (pri-miRNAs) correlates with locations of annotated mature miRNAs.** Small-RNA sequencing read depth (per million) was plotted across the sequence of annotated primary micro-RNA transcripts (pri-miRNAs) for wild-type (black), HYL1$^-$ knockout (red) and ancFpHYL1 (blue) *A. thaliana* genotypes. Gray bars indicate parts of each pri-miRNA sequence annotated as mature miRNAs.

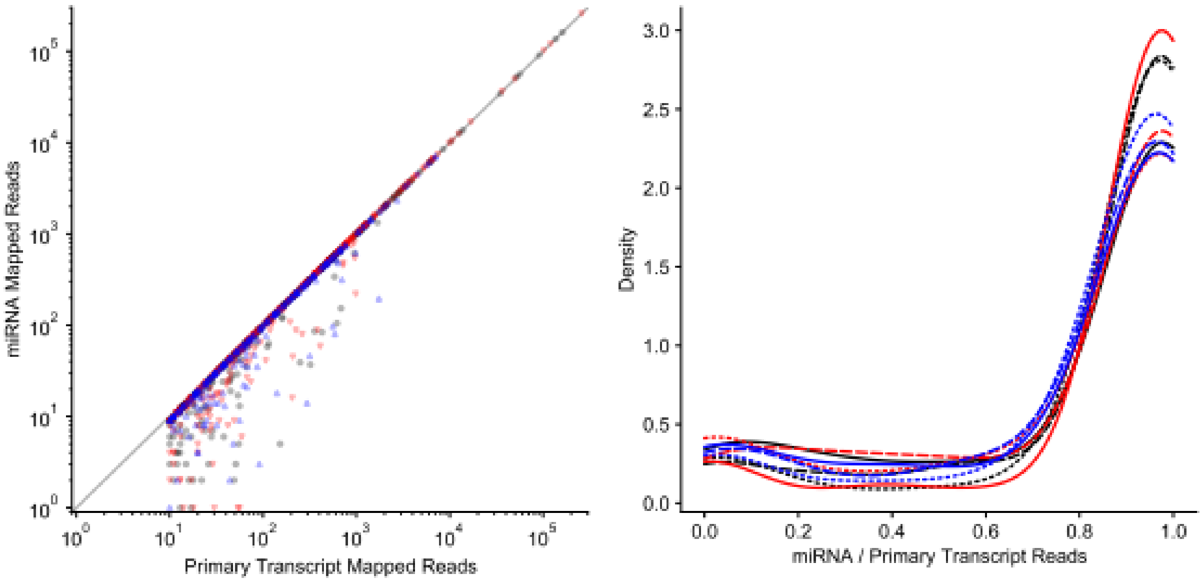


**Figure S12. Small-RNA sequencing reads mapping to primary miRNA transcripts (pri-miRNAs) overwhelming map to annotated mature miRNAs within the pri-miRNA sequence.** We counted the total numbers of small-RNA sequencing reads mapping to annotated pri-miRNA sequences within the *A. thaliana* genome from replicate wild-type (black) HYL1$^-$ knockout (red) and ancFpHYL1 (blue) plants. Left panel plots the number of reads mapping to each pri-miRNA sequence vs the number of reads mapping to annotated mature miRNAs within the pri-miRNA sequence. Data points would fall on the diagonal line if all reads mapping to the pri-miRNA fell within the mature miRNA region. Data points falling below the diagonal line indicate an excess of sequencing reads mapping to areas of the pri-miRNA outside annotated mature miRNA regions. Left panel shows a kernel density plot of the ratio of sequencing reads mapping to annotated mature miRNA regions within each pri-miRNA sequence, divided by the total number of reads mapping to the pri-miRNA. The value of 1.0 on the x-axis indicates the density of data points from each plant in which *all* of the reads mapping to the pri-miRNA fell within annotated mature miRNA regions.

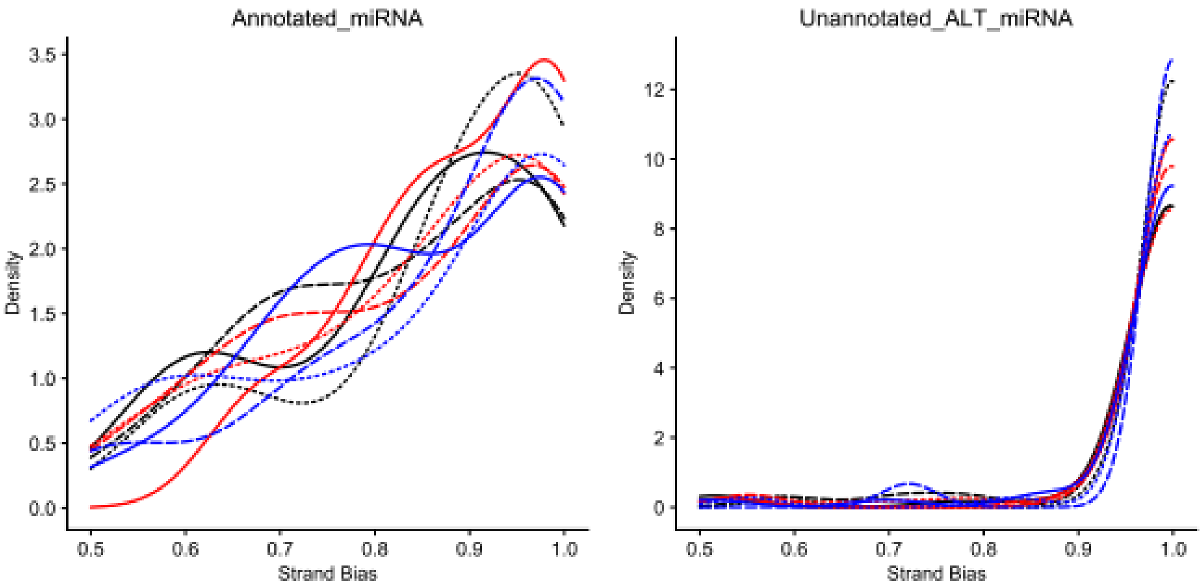


**Figure S13. The majority of small-RNA sequencing reads mapping to pri-miRNAs were obtained from one of the annotated mature miRNA guide strands, suggesting that guide strands potentially functioning in RNAi are responsible for the majority of small-RNA reads.** For each pri-miRNA sequence, we calculated the “strand bias” as the ratio of the number of reads mapping to each annotated mature miRNA guide strand, divided by the total number of reads mapping to the pri-miRNA. A strand bias of 0.5 would indicate that approximately ½ of the total reads mapped to each of the potential miRNA guide strands, which would suggest small-RNA sequencing was primarily generating sequence reads directly from miRNA duplexes. We plot kernel densities of strand biases obtained from each wild-type (black), HYL1$^{-}$ (red) and ancFpHYL1 (blue) *A. thaliana* plant. Left panel shows data from all annotated pri-miRNAs. Right panel shows data from only those annotated pri-miRNAs in which only a single mature miRNA is annotated in the Araport 11 genome annotation.
